# Mechanistic origins of temperature scaling in the early embryonic cell cycle

**DOI:** 10.1101/2024.12.24.630245

**Authors:** Jan Rombouts, Franco Tavella, Alexandra Vandervelde, Connie Phong, James E. Ferrell, Qiong Yang, Lendert Gelens

**Affiliations:** Laboratory of Dynamics in Biological Systems, Department of Cellular and Molecular Medicine, KU Leuven, Herestraat, 49, Leuven, Belgium; Cell Biology and Biophysics Unit and Developmental Biology Unit, European Molecular Biology Laboratory (EMBL), Heidelberg, Germany; Department of Physics / Biophysics, University of Michigan, Ann Arbor, MI 48109, USA; Department of Chemical and Systems Biology, Stanford University School of Medicine, Stanford, CA 94305-5174, USA; Unit of Theoretical Chronobiology, Université Libre de Bruxelles, Brussels, Belgium

**Keywords:** Cell cycle, Cell-free extract, Xenopus laevis, Temperature, Thermal limits

## Abstract

Temperature profoundly impacts organismal physiology and ecological dynamics, particularly affecting ectothermic species and making them especially vulnerable to climate changes. Although complex physiological processes usually involve dozens of enzymes, empirically it is found that the rates of these processes often obey the Arrhenius equation, which was originally proposed for individual chemical reactions. Here we have examined the temperature scaling of the early embryonic cell cycle, with the goal of understanding why the Arrhenius equation approximately holds and why it breaks down at temperature extremes. Using experimental data from *Xenopus laevis, Xenopus tropicalis*, and *Danio rerio*, plus published data from *Caenorhabditis elegans, Caenorhabditis briggsae*, and *Drosophila melanogaster*, we find that the apparent activation energies (*E*_*a*_ values) for the early embryonic cell cycle for diverse ectotherms are all similar, 75 ± 7 kJ/mol (mean ± std.dev., n = 6), which corresponds to a *Q*_10_ value at 20°C of 2.8 ± 0.2 (mean ± std.dev., n = 6). Using computational models, we find that the approximate Arrhenius scaling and the deviations from it at high and low temperatures can be accounted for by biphasic temperature scaling in critical individual components of the cell cycle oscillator circuit, by imbalances in the *E*_*a*_ values for different partially rate-determining enzymes, or by a combination of both. Experimental studies of cycling *Xenopus* extracts indicate that both of these mechanisms contribute to the general scaling of temperature, and *in vitro* studies of individual cell cycle regulators confirm that there is in fact a substantial imbalance in their *E*_*a*_ values. These findings provide mechanistic insights into the dynamic interplay between temperature and complex biochemical processes, and into why biological systems fail at extreme temperatures.

## Introduction

Living organisms are continually influenced by their environment, and embryos are particularly sensitive to environmental changes. Even subtle perturbations during critical developmental windows can significantly impact embryo viability, as well as embryonic and post-embryonic performance (1). A key aspect of the environment is temperature: changes in temperature can profoundly influence embryonic development, influencing the speeds of biochemical reactions and affecting the overall physiology, behavior, and fitness of the organism (2–6).

Ectotherms, in particular, rely critically on the ambient temperature as they have minimal ability to generate heat internally. Each ectothermic species has a specific temperature range associated with its geographic distribution on the planet (7, 8). While adult stages of ectotherms can adopt various physiological and behavioral strategies to maintain optimal temperatures, such as taking shelter when it is too hot (6), embryos possess limited mechanisms to cope with environmental challenges, making them the most vulnerable life stage to environmental stress (9, 10). Temperature plays a pivotal role in determining the fertilization rate of eggs, the growth and survival of embryos, and in certain cases, even the gender of offspring, as observed in many turtle species and all crocodiles (11). Assessing the thermal impact on and sensitivity of embryonic development across a range of temperatures provides essential insights into species’ responses and vulnerability to the challenges posed by global warming (8, 9, 12). Indeed, the impact of global warming is already evident in certain sea turtle species, where a diminishing number of male offspring is observed (13).

But even without this shifting landscape and the challenges of their changing ecosystems, ectotherms face the daunting challenge of needing to have their biochemistry function reliably over a wide range of temperatures. Given that complex metabolic networks, signaling systems, and developmental processes may involve dozens of enzymes, the question arises as to how much variation in the individual enzymes’ temperature scaling can be tolerated before the system fails.

The influence of temperature on physiology and development has been a subject of study for over a century (3, 4, 14). The relationship between temperature and the speed of many diverse biological processes is often well approximated by the Arrhenius equation (15–22). Originally formulated for simple, one-step chemical reactions, the Arrhenius equation describes the rate of a chemical reaction (*k*) as a function of the absolute temperature (*T*):

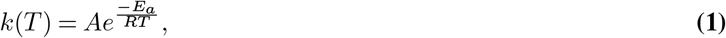

where *E*_*a*_ denotes a temperature-independent activation energy, *R* is the universal gas constant, and the pre-exponential factor *A* sets the maximal reaction rate at high temperatures. While derivable for elementary chemical reactions from thermodynamic principles, this equation is considered an empirical law applicable to various physiological rates (16). However, deviations from this Arrhenius response are consistently observed at higher temperatures (18, 22–25). The decline is often attributed to the heat denaturation of some critical enzyme (18, 25–27), and the trade-off between increasing reaction rate and increasing denaturation yields an optimal temperature. Moreover, the thermal response of processes at the cellular level is influenced not only by the reactions of enzymes but also by active cellular responses to temperature changes. For instance, in response to stress, cells may up-regulate heat shock proteins, aiding in protein refolding (28). Another example is found in budding yeast, which can up-regulate the production of viscogens like trehalose and glycogen to help maintain normal diffusion kinetics at elevated temperature (29).

Recently, the topic of biological temperature scaling has garnered renewed interest, thanks to the ability to obtain accurate, high-resolution data through time-lapse microscopy. This approach has provided fresh insights into early development in several model systems (30–33). Overall, these findings reaffirm the utility of the Arrhenius equation as a reliable approximate description of the temperature scaling of embryonic development. This was particularly clear in studies of the timing of the first embryonic cell cycle in *C. elegans* and *C. briggsae*, two closely related nematodes. In both species, the duration of the cell cycle as a function of temperature precisely agreed with the Arrhenius equation over a broad temperature range, with some deviation then occurring when the embryos were close to their maximum tolerated temperatures (31). The two nematodes also were found to have almost identical Arrhenius energies (*E*_*a*_ values), which raises the possibility that the activation energies of cell cycle regulators may be evolutionarily constrained to a single standard value (31).

*Xenopus laevis* extracts and embryos have proven to be powerful systems for the quantitative analysis of the early embryonic cell cycle. The cell cycle can be experimentally studied both *in vivo* and in extracts, and extracts can be manipulated and observed in ways that are difficult with intact embryos. In addition, much is already known about cell cycle biochemistry in this system, providing a rich and highly quantitative context for further studies. And finally, the dynamics of the cell cycle can be successfully reproduced with relatively simple mathematical models that can add depth to the understanding of experimental findings. Crapse and colleagues have begun to examine how the *Xenopus* embryonic cell cycle is affected by temperature, and they have found some striking similarities to the behaviors seen in *C. elegans* and *C. briggsae*: the cell cycle period obeys the Arrhenius equation at least approximately, and the measured *E*_*a*_ value for the cell cycle is similar to those in the two nematodes

(32).

Here we have leveraged the *Xenopus* system to address several outstanding questions on the principles of biological temperature scaling. First, we compared the *Xenopus laevis* temperature scaling to that in two other ectothermic vertebrate model systems, *Xenopus tropicalis* and *Danio rerio*, and compared the findings to previously reported data from the invertebrates *C. elegans, C. briggsae*, and *Drosophila melanogaster*. Second, we asked how well the temperature scaling is described by the Arrhenius equation, and based on ordinary differential equation modeling of the cell cycle, under what circumstances would the cell cycle be expected to exhibit Arrhenius scaling, and under what circumstances would it be expected to deviate. Finally, we asked how the different individual phases of the cell cycle vary with temperature, and found that interphase and mitosis scale differently and that this difference can be accounted for by the *in vitro* thermal properties of key cell cycle regulators. These studies provide insight into the principles that allow ectotherms to tolerate a range of temperatures, and suggest mechanisms for why the cell cycle oscillator fails at temperature extremes.

## Results

### Temperature scaling of cell division timing in the *Xenopus laevis* embryo

We measured the temperature dependence of the timing of several early cell cycle events in the developing *Xenopus laevis* embryo (Figure 1A-B), similar to the work of Crapse and colleagues (32). *Xenopus laevis* eggs were fertilized and imaged in a temperature-controlled chamber (first described in Ref. (34)) by time-lapse microscopy (Fig. 1A). We then analyzed the movies (Supplemental Videos 1-2) to visually identify various early developmental events (Fig. 1B). First, we scored the start of the fertilization wave, a ripple in the egg’s cortex that quickly spreads from the sperm entry point across the egg (at time *t*_FW_ after fertilization). This wave is due to a trigger wave of elevated intracellular calcium, and it contributes to the block to polyspermy and coordinates the start of the cell cycle (35). Next, we measured the start of the first surface contraction wave, which emanates from the animal pole and travels toward the vegetal pole (36, 37) (at time *t*_SCW_ after fertilization). This wave marks mitotic entry and has been argued to be caused by the interaction of a spherical wave of Cdk1 activation originating at the nucleus (38–40) with the cortical cytoskeleton (41–43). Finally, we assessed the cleavages that complete each of the first four cell cycles. The first cleavage begins about 95 min after fertilization at 18°C, and the next several cycles occur every 35 min thereafter (44–46). For multicellular embryos, we took the time at which the earliest cell began to divide to be the cleavage time, but note that within an embryo, these cell divisions were nearly synchronous. The timing of all of these events was recorded for about 10 different embryos at each temperature.

**Fig. 1:**
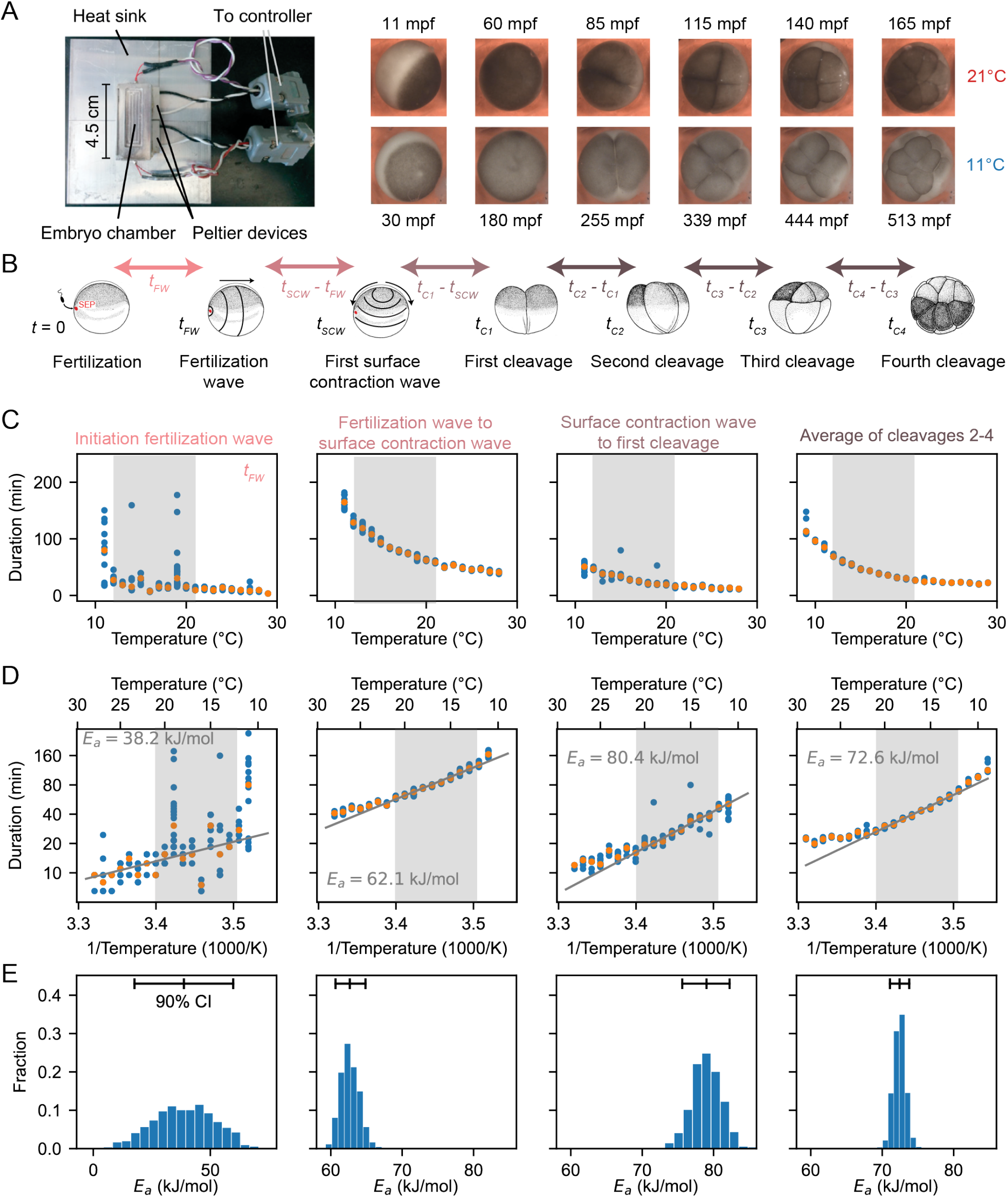
Cell division timing in early Xenopus laevis embryos scales approximately Arrhenius over a wide range of temperatures. A. *Xenopus laevis* embryonic development was imaged in a temperature-controlled chamber introduced in Ref. (34). The time unit mpf is minutes post-fertilization. B. Different early developmental events were visually identified. SEP denotes the sperm entry point. Adapted from (98). C. Duration of several early developmental periods in function of temperature in the range [*T*_min_ = 9°C,*T*_max_ = 29°C]. D. An Arrhenius fit is shown for the values between 12°C and 21°C, with the apparent activation energy indicated. E. Bootstrapping provides a probability distribution for the apparent activation energies. The mean and 90% confidence interval (CI) are also indicated.

Fertilized embryos reliably progressed through the cell cycle and divided at temperatures between 10°C and 28°C. Just outside this range (down to 9°C and up to 29°C) some cell cycles still occurred, and these data are included in Fig.1. We quantified the time intervals between these developmental events and examined their temperature dependence (Fig.1C-D), using a rearranged form of the Arrhenius equation to relate the duration of a process, Δ*t* = 1*/k*[*T*], to absolute temperature:

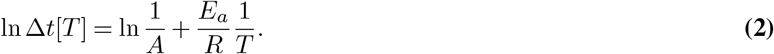

Accordingly, we replotted the data as ln Δ*t* versus 1*/T* (Fig.1D). Between 12°C and 21°C, the data were well-approximated by Eq. (2), and from the fitted slopes we extracted apparent activation energies (*E*_*a*_) of approximately 60–80 kJ/mol (Fig. 1D, Fig. S1). An exception was the onset time of the fertilization wave, which showed a lower *E*_*a*_ (*∼*40 kJ/mol) with a wide confidence interval. These values are consistent with previous reports (32) and fall within the typical enzymatic range of 20–100 kJ/mol (47–49). Outside the 12–21°C range, the durations deviated from linearity, with unexpectedly long times at both low and high extremes (Fig. 1D). To assess the robustness of these differences, we used a bootstrapping approach to generate probability distributions of the apparent activation energies (Fig. 1E, Fig. S1, Fig. S2A, Supplementary Note 2). Finally, we computed the mean square error (MSE) between the data and the Arrhenius fit, both across the full temperature range (9°C to 29°C) and within the linear range (12°C to 21°C). As expected, the MSE was substantially higher across the full range, confirming that the Arrhenius model does not adequately describe the entire dataset (Fig. S2B).

### Diverse ectothermic species yield similar temperature scaling

Next we examined the timing of the early cell cycles in two additional vertebrate model organisms, the frog *X. tropicalis* (Supplemental Videos 3-4) and the zebrafish *D. rerio* (Supplemental Videos 5-6). The three vertebrates and the two nematodes span a broad range in evolution (Fig. 2A).

**Fig. 2:**
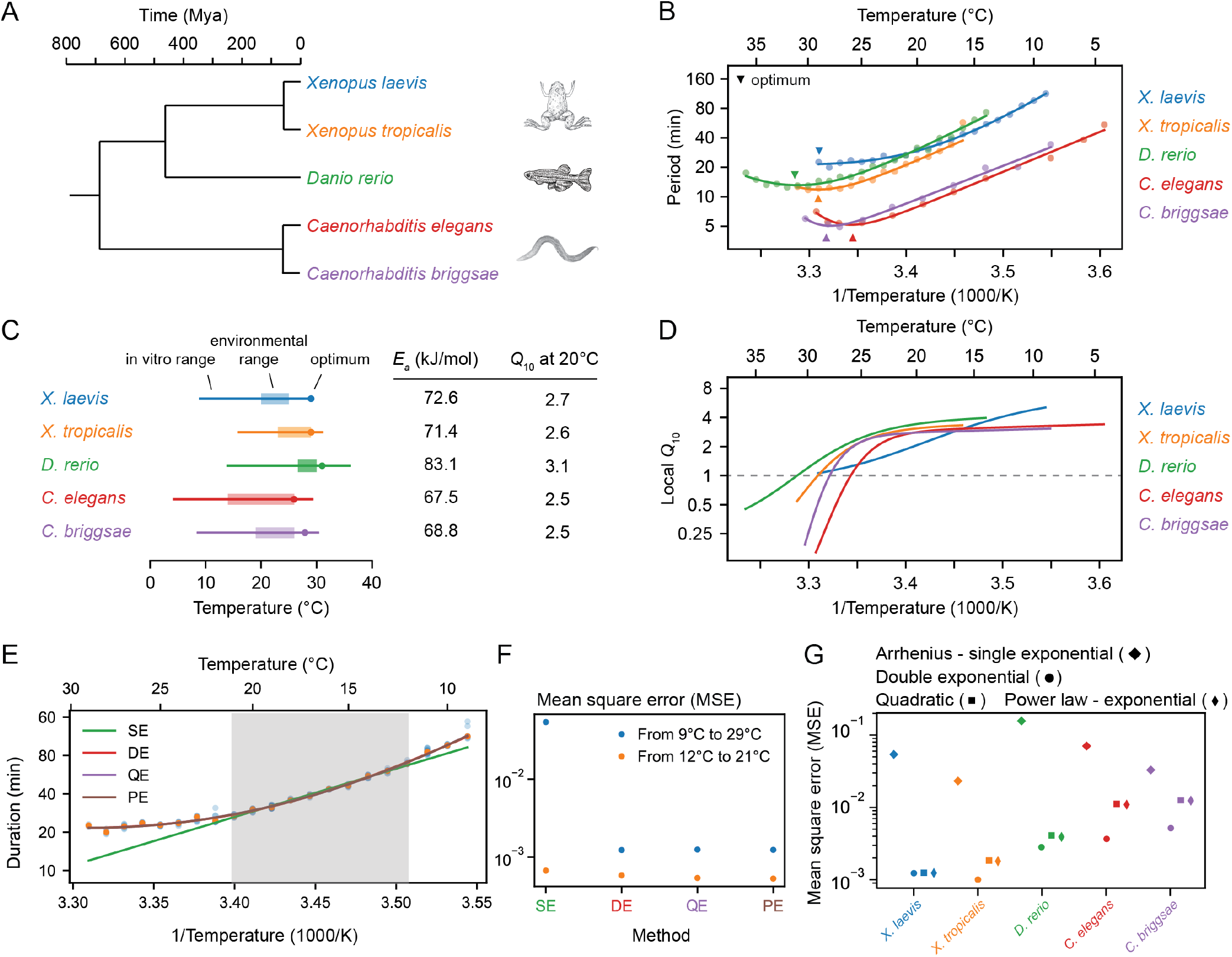
Cell cleavage period scales in a similar non-Arrhenius way across different early ectothermic embryos. A. We examined the timing of the early cell cycles in 5 different species: *C. elegans* and *C. briggsae* (from (31)), *D. rerio* (this work), *X. tropicalis* and *X. laevis* (this work). The three vertebrates and the two nematodes span a broad range in evolution. B. Median cleavage period in function of temperature for the early cell cleavages (all pooled) for the 5 different species. Optimal fits using a double exponential (DE) function are overlayed. C. The *in vivo* range of viable early cell cycles in the different species, including their thermal limits and optimal temperature at which they reach a minimum cell cycle period. Their corresponding apparent activation energies and *Q*_10_ at 20°C are shown in the table. Additionally the environmental range is indicated for all five organisms (99–102). D. Using the best DE fit, the local *Q*_10_ value is plotted in function of temperature. E. The median cleavage period in function of temperature for *X. laevis* is fitted using different functional forms: single exponential Arrhenius (SE), double exponential (DE), quadratic exponential (QE) and a power law – exponential (PE) function. F. The goodness of fit (using mean square error, MSE on the logarithms of the periods) of the alternative functional forms to the experimental data for *X. laevis* in two different temperature regions: 12-21°C and 9-29°C. G. Goodness of fit, similar as in panel F, but now for all different species over their whole measured temperature range.

The period of the early embryonic cell cycle as a function of temperature for all five organisms is shown in Fig. 2B. For simplicity, here we have pooled data for the durations of the early cell cleavages. In all cases, the early embryonic cell cycle could proceed over a 15-25°C range of temperatures. In general, the five organisms showed reasonable agreement with the Arrhenius equation, especially toward the lower end of their temperature ranges (Fig. 2B). *Xenopus laevis* was something of an outlier in this regard; its Arrhenius plot is bowed throughout the temperature range (Fig. 2B, blue). The apparent Arrhenius energies—the slopes of the Arrhenius plots—were quite similar, ranging between 68 and 83 kJ/mol, or 73 ± 6 kJ/mol (mean ± std. dev., n = 5).

As might be expected, the nominal ambient environmental temperatures for all five organisms fell within the range found to be compatible with cell cycle oscillations (Fig. 2C). The temperature ranges for *Xenopus tropicalis* and *Danio rerio* were shifted toward higher temperatures compared to *Xenopus laevis*, reflecting the fact that the former two evolved in and live in warmer regions (Fig. 2C, orange and green vs. blue). A similar shift in the viable temperature range has been noted for the nematode worms *C. elegans* and *C. briggsae* (31), replotted here in red and purple. In all cases, the maximum temperature compatible with cycling was closer to the nominal environmental temperature range than the minimum temperature was (Fig. 2C).

For four of the five organisms (*C. elegans, C. briggsae, D. rerio*, and *X. tropicalis*) there was sufficient upward deflection of the temperature curves toward the high end of the temperature range to define an optimal growth temperature corresponding to a minimal cell cycle duration (Fig. 2C). For *Xenopus laevis*, the fastest cell cycles were found at the highest temperature compatible with viability (Fig. 2C). In all cases the optimal temperature was within a few degrees of the maximal permissible temperature *T*_max_ (Fig. 2C). The optimal temperatures were generally somewhat higher than the typical environmental temperature ranges (Fig. 2C). This may reflect a trade-off between maximal speed at higher temperatures and maximal safety margins in the middle of the operating temperature range. Note that at the temperature optima, the slopes of the Arrhenius plots are zero. The curves are also shallow at the optima, which means that changes of several degrees produce little changes in the cell cycle period. The period can be regarded as temperature-invariant or temperature-compensated in this regime.

Data are also available for the temperature scaling of various embryonic processes in *Drosophila melanogaster*. Extensive data are available for the timing between the 13th and 14th cleavage (32, 50), and these data are replotted in Fig. S3A,B. Like the *Xenopus laevis* Arrhenius plot, the *Drosophila* plot is bowed throughout the temperature range, and, at the cold end of the temperature range, it yielded an Arrhenius energy of 109 kJ/mol. Note, however, that this cycle differs from the earlier *Drosophila* cycles and the other embryonic cycles examined here as they have a longer cell cycle due to lengthening of S-phase approaching the mid-blastula transition (MBT) (50, 51). Some data are available for the 11th nuclear cycle (NC11) in the syncytial *Drosophila* embryo (Fig. S3A) (33); this cycle is more similar to the other organisms’ cycles analyzed here. From the published data, we calculated an Arrhenius energy of 84 kJ/mol for the duration of NC11, slightly higher than the energies calculated for the early embryonic cycles of the other 5 model organisms. Taken together, the six organisms yielded an average Arrhenius energy for the early embryonic cell cycles of 75 ± 7 kJ/mol (mean ± std. dev.). Although the nominal periods of the cell cycles varied greatly, from about 5 min for *C. elegans* and *C. briggsae* to 25 min for *X. laevis* at room temperature, the temperature scaling factors for these organisms varied by only about 9%.

The temperature sensitivity of the early embryonic cell cycle can also be characterized by *Q*_10_ values, which capture how reaction rates change over 10°C intervals (52). While linear Arrhenius fits give a global *Q*_10_ value around 2.8 across diverse organisms (Fig. 2C), local *Q*_10_ values (see Supplementary Note 1) reveal important deviations, particularly at temperature extremes (Fig. 2D). For most species, local *Q*_10_ values plateau around 2.8 *±* 0.4 at low temperatures and decrease at higher temperatures. However, in *Xenopus laevis*, local *Q*_10_ values vary more broadly, ranging from about 1 to 4 across the full temperature range (Fig. 2D, Fig. S3E).

To better capture the deviations from idealized Arrhenius behavior, we explored three commonly used generalizations of the Arrhenius equation—the double exponential (31, 53, 54), quadratic exponential (32, 55), and power law–exponential forms (56, 57) (see Methods for details on the fitting). All three models provided markedly better fits to the experimental data than the classical Arrhenius relationship, capturing the nonlinearities in the temperature dependence with high accuracy (Fig. 2E-G). However, their comparable performance makes it difficult to identify a single best functional form based on fitting alone. This underscores the limitations of descriptive models and points to the need for mechanistic frameworks to explain the origin of temperature scaling curves in biological systems.

### A simple relaxation oscillator model for the early embryonic cell cycle can account for approximate Arrhenius scaling as well as the observed deviations from Arrhenius scaling

We next took a computational approach to the question of why cell cycle periods at least approximately obey the Arrhenius equation, and why they sometimes deviate from Arrhenius scaling. We used a differential equation model of the embryonic cell cycle oscillator to investigate how the period would be expected to vary with temperature, given either single or double exponential equations for the individual enzymes’ temperature scaling. This allowed us to examine how systems-level properties of the cell cycle oscillator circuit, rather than just variations from the Arrhenius relationship in the behaviors of individual enzymes, might be expected to affect the temperature scaling of the oscillations.

The cell cycle regulatory network consists of many complex interactions involving dozens of species, which makes it extremely challenging to construct a complete mathematical model, let alone study and interpret the influence of temperature on the cell cycle. The early embryonic cell cycle of insects, worms, amphibians and fish is, however, much simpler: the cycle consists of a rapidly alternating sequence of synthesis (S) phase and mitotic (M) phase, without checkpoints and without G1 and G2 gap phases. Transcription is negligible at this point in embryogenesis, and the number of protein species involved is smaller than in the somatic cell cycle. As a result, simpler mathematical models can be constructed. This greatly simplifies the analysis of temperature scaling.

At the heart of the early embryonic cell cycle lies the protein complex cyclin B – Cdk1, consisting of the protein cyclin B and the cyclin-dependent-kinase Cdk1 (Fig. 3A), plus a phospho-epitope-binding subunit, Suc1/Cks, that is not separately considered here. When this protein kinase complex is enzymatically active, it phosphorylates hundreds of other proteins, bringing about the entry of the cell into mitosis (58, 59). The oscillations in Cdk1 activity result from three interlinked processes: (1) cyclin B synthesis, which dominates in interphase and causes Cdk1 activity to gradually rise; (2) the flipping of a bistable switch, due in this model to the Cdk1-Cdc25 positive feedback loop and the Cdk1-Wee1 double negative feedback loop, which results in an abrupt rise in Cdk1 activity; and (3) cyclin B degradation by the anaphase-promoting complex/cyclosome APC/C and the proteasome, which dominates during M-phase and causes Cdk1 activity to fall (Fig. 3B) (60–62). This type of oscillator, consisting of a rapid bistable switch plus a negative feedback loop, is referred to as a relaxation oscillator (63–65). Relaxation oscillators are common in biology, and all relaxation oscillators share similar qualitative behavior, irrespective of the exact molecular details: there is a slow ramp up in activity, which then triggers an abrupt burst in activity through the positive feedback loop(s), and finally the negative feedback restores the system back to its low activity state. These three distinct phases can be distinguished in experimental data on the activity of Cdk1 as a function of time in the early embryonic cell cycle (Fig. 3C; see also below).

**Fig. 3:**
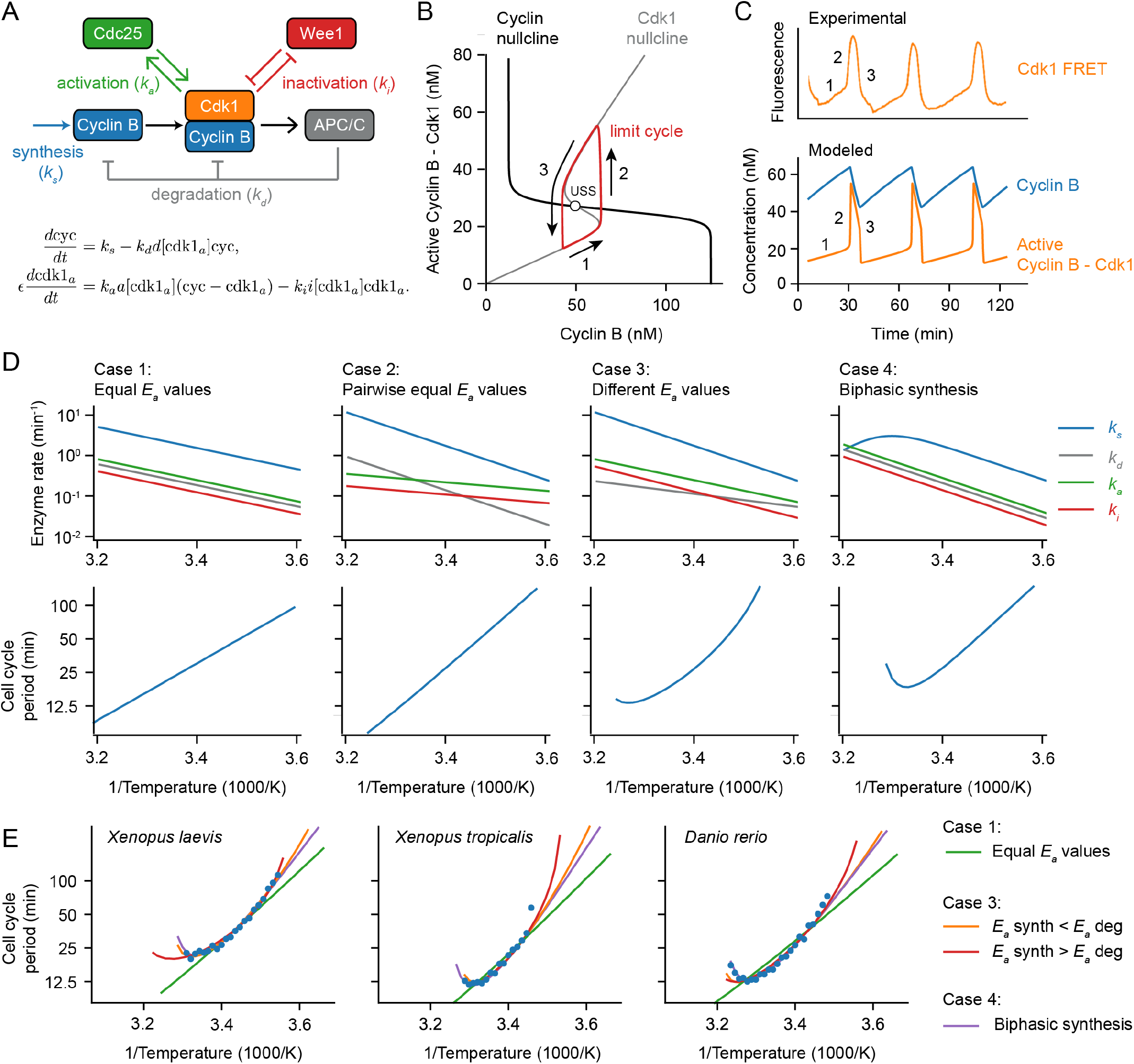
A simple relaxation oscillator model for the early embryonic cell cycle can reproduce the observed non-Arrhenius scaling. A. Sketch of key reactions in the early cell cycle regulatory network. B-C. Phase plane representation (B) and time series (C) of cell cycle oscillations in Eq. (17). D. Scenarios showing how different temperature scaling of cell cycle regulatory processes can lead to Arrhenius scaling and/or thermal limits in the scaling of the cell cycle period. E. Best fits of models presented in panel D to the measured data for the early cell cycle duration for *X. laevis, X. tropicalis* and *D. rerio* shown in Fig. 2. For case 3, the apparent activation energies for *k*_*s*_ and *k*_*d*_ need to be different to fit the data well. For Case 4, we introduced a biphasic response in cyclin B synthesis (*k*_*s*_). For parameter values and more details about the model, see Supplementary Note 3.

We described the changes in cyclin concentration and Cdk1 activity with a model consisting of two ordinary differential equations (ODEs) (66):

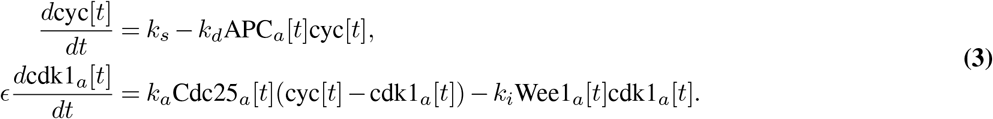

The first equation describes how cyclin B (cyc) is synthesized throughout the cell cycle at a rate *k*_*s*_ (nM/min) and how it is degraded at a rate *k*_*d*_ (1/min) by the proteasome after ubiquitination by active APC/C (APC_*a*_[*t*]). The second equation describes the conversion of cyclin B-Cdk1 complexes between an inactive form and an active form by Cdc25 (Cdc25_*a*_[*t*]) and Wee1 (Wee1_*a*_[*t*]). For simplicity, we do not directly include degradation of the bound-form of cyclin B (see Supplementary Note 3.A). If we assume that the Cdk1-mediated phosphorylation reactions that regulate APC/C, Wee1, and Cdc25 are essentially instantaneous, we can eliminate three of the time-dependent variables from the right hand side of the ODEs:

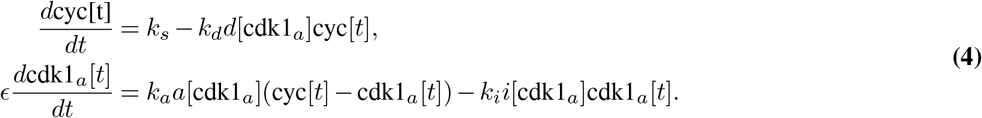

The terms *d*[cdk1_*a*_], *a*[cdk1_*a*_], and *i*[cdk1_*a*_] are assumed to be Hill functions of the instantaneous values of cdk1_*a*_, and were parameterized based on experimental measurements of steady-state responses in *Xenopus laevis* extracts (66–68). We thus have two ODEs in two time-dependent variables, cyc[*t*] and cdk1_*a*_[*t*], and five parameters that define the speeds of cyclin synthesis (*k*_*s*_), cyclin degradation (*k*_*d*_), Cdk1 activation by Cdc25 (*k*_*a*_), and Cdk1 inactivation by Wee1 (*k*_*i*_), as well as the relative time scales of cyclin synthesis and degradation versus Cdk1 activation and inactivation (*ϵ*). Even though this simplified model omits Greatwall/PP2A-B55 regulation and a number of other interesting aspects of *Xenopus* cell cycle regulation (62, 65, 69–72), it nevertheless captures the dynamics of Cdk1 activation and inactivation well and has been used to successfully describe various aspects of cell cycle oscillations (34, 66, 73). For this reason it seemed like a good starting point for understanding how the output of the cell cycle oscillator circuit would be expected to scale with temperature.

Using experimentally motivated parameters (66), the model (Eq. (4)) reproduced cell cycle oscillations with a realistic period of approximately 30 min (Fig. 3C). These oscillations manifested as a closed trajectory, a limit cycle, in the (cyc, cdk1_*a*_) phase plane (Fig. 3B, red), which orbits around an unstable steady state (Fig. 3B, USS) at the intersection of the system’s two nullclines (Fig. 3B). When the time scale for Cdk1 activation/inactivation is fast relative to the time scale of cyclin synthesis and degradation (*ϵ ≪* 1), typical relaxation oscillations occur: in interphase, the orbit slowly creeps up the low Cdk1 activity portion of the S-shaped nullcline (Fig. 3B, denoted 1), then abruptly jumps up to the high Cdk1 activity portion of the same nullcline (Fig. 3B, denoted 2), crawls down the nullcline due to active APC/C (Fig. 3B, denoted 3), and abruptly falls back down to the lower portion of the nullcline to begin the cycle again. The result is sawtooth-shaped oscillations in cyclin B levels and periodic bursts of Cdk1 activity that resemble experimentally-measured Cdk1 activities (Fig. 3C).

Next we examined how making the model’s parameters temperature-dependent affected the persistence and period of the oscillations over a range of temperatures. We started by assuming Arrhenius scaling for the four key rates in the oscillator model: *k*_*s*_, *k*_*d*_, *k*_*a*_, and *k*_*i*_. As expected, when all of the apparent activation energies were assumed to be equal, the oscillation period obeyed the Arrhenius equation with the same activation energy (Case 1 in Fig. 3D). Similarly, if the activation energies for cyclin synthesis and degradation were assumed to be equal, and the energies for Cdk1 activation and inactivation were assumed to be different from those but equal to each other, the period also scaled in an Arrhenius fashion (Case 2 in Fig. 3D), with *E*_*a*_ equal to that of cyclin synthesis and degradation. This Arrhenius scaling arises out of three properties of the model: (1) the cyclin nullcline’s position depends solely on the ratio *k*_*s*_*/k*_*d*_, and these two parameters were assumed to scale identically with temperature; (2) the location of the S-shaped Cdk1 nullcline depends upon the ratio *k*_*a*_/*k*_*i*_, which likewise was assumed to scale identically with temperature; and (3) as long as *ϵ* is very small, the cell cycle period is determined only by the rates of cyclin synthesis and degradation, which scale identically with temperature.

However, if the four kinetic parameters were not constrained to scale identically (Case 1) or pair-wise identically (Case 2) with temperature, the results were more like what is seen experimentally. This is shown as Fig. 3D, Case 3. The Arrhenius plot was bowed concave up, instead of being straight, and oscillations ceased if the temperature was too high or too low. The cessation of oscillations can be rationalized from the positions of the nullclines in the phase plane. If the *E*_*a*_ value for cyclin synthesis is smaller than that for cyclin degradation, the ratio *k*_*s*_*/k*_*d*_ decreases as temperature rises. With increasing temperature, the cyclin nullcline shifts down until it no longer intersects the middle portion of the S-shaped Cdk1 nullcline. At this point, the steady state becomes stable and oscillations cease, leaving the system in an interphase-like steady state with low Cdk1 activity (Fig. S4A). Conversely, if cyclin synthesis scales more strongly with temperature than degradation does, the cyclin nullcline shifts upward, leading to a stable M-phase-like steady state with high Cdk1 activity (Fig. S4C). Thus, if the temperature scaling of cyclin synthesis and degradation differ, at extremes of the temperature range the oscillator will fail, and in between the extremes the cell cycle period would be expected to deviate from the Arrhenius relationship. This could provide an explanation for the temperature scaling observed experimentally (Fig. 1).

An alternative assumption could also explain the experimental results. As shown in Fig. 3D, Case 4, if at least one process shows a biphasic dependence of rate on temperature, perhaps due to enzyme denaturation at high temperatures, the result will be a bowed Arrhenius plot and a high temperature limit to oscillations. As an example, here we have assumed a biphasic dependence of cyclin synthesis on temperature. Thus, in principle it seemed like either variation in the individual enzymes’ Arrhenius energies (Case 3), or denaturation at high temperatures (Case 4), or both, could account for the observed temperature scaling of the early embryonic cell cycle.

To test this hypothesis further, we asked how well model Cases 3 and 4 could replicate the observed cell cycle duration scaling in early frog and zebrafish embryos. We adjusted the apparent activation energies of cyclin synthesis and degradation, and, for simplicity, kept the *E*_*a*_ values for Cdk1 activation and inactivation constant. Through minimizing the error between simulations and data across the oscillation range, employing the mean sum of squares on logarithms of periods, we obtained optimal fits. Fig. 3E displays these fits across various model scenarios introduced in Fig. 3D. Although perfect Arrhenius scaling (Cases 1 and 2) did not align well with the data, model Cases 3 and 4 approximated the measured data well. Notably, the experimental data could be accounted for by assuming that cyclin synthesis scaled either more strongly or more weakly with temperature than did cyclin degradation.

To further reinforce these findings, we conducted an exhaustive parameter scan over the activation energies of all four key rates in the oscillator model using a fitting algorithm. We employed the approximate Bayesian computation method (74), implemented in Python using pyABC (75), for sequential Monte Carlo sampling of parameter sets, gradually improving fits to the data (for details, see Supplementary Note 3.D and Fig. S5). This broader analysis underscored that optimal fits occurred when there was a distinct difference in apparent activation energies between cyclin synthesis and degradation, while the activation energies of Cdk1 activation and inactivation remained similar (Fig. S5). Moreover, the quantification of fitting errors revealed that it is most probable that the experimental data is due to cyclin synthesis being more sensitive to temperature changes than cyclin degradation (*E*_*a*_(*k*_*s*_)>*E*_*a*_(*k*_*d*_)).

Next, we asked whether the results obtained were specific to the two-ODE cell cycle model. To explore this, we turned to a structurally distinct model: a five-ODE, mass-action-based system that includes interactions among Cdk1, Greatwall, and PP2A—elements that collectively form a mitotic switch as well (62, 65, 76). In contrast to the two-ODE model, which featured a bistable switch between Cdk1 and Cyclin B and included feedback through Cdc25 and Wee1, this model implements a bistable switch between APC/C and Cdk1, and thus omits the Cdc25/Wee1-mediated feedback loops entirely. It also differs in its use of strictly mass-action kinetics, avoiding the highly nonlinear Hill functions of the two-ODE model, and in its dimensionality, expanding from two to five ODEs. Despite these structural differences, both models share key features: cyclin synthesis and Cdk1-activated degradation, the presence of a bistable switch, and a separation of timescales enabling relaxation oscillations. The five-ODE model could be parameterized to yield realistic cell cycle oscillations (77) (Fig. S6B, Supplementary Note 3.B). Due to the model’s increased complexity—ten kinetic parameters—we relied exclusively on the ABC algorithm for parameter inference. This approach produced satisfactory fits (Fig. S6E), and analysis revealed that highly correlated activation energy pairs typically corresponded to antagonistic reaction rates (Fig. S6F,G, Supplementary Note 3.B). These results again show that well-fitting parameter sets tend to exhibit similar activation energies for faster reactions. Moreover, they support the idea that thermal limits can arise from imbalances in the apparent activation energies of cyclin synthesis and degradation, reinforcing the conclusions drawn from the two-ODE model.

In summary, computational modeling revealed that thermal limits and non-Arrhenius scaling like those seen in early embryos can arise from (at least) two different mechanisms. Firstly, in cases where all rates follow Arrhenius-like scaling but possess varying activation energies, an imbalance emerges, culminating in a thermal limit and a bowed Arrhenius plot. We can call this behavior ‘emergent’, since the limit and the bowing are not inherent to any individual reaction but arise collectively. Secondly, thermal limits can arise if one or more underlying reactions exhibit a thermal optimum and deviate from Arrhenius scaling. Here, the system’s behavior is predominantly dictated by the dynamics of the particular biphasic component(s).

### The durations of interphase and M-phase scale differently with temperature

To test whether the emergent imbalance model (Fig. 3C, Case 3) contributes to the temperature scaling of the *Xenopus laevis* embryo, we set out to determine how the durations of interphase and M-phase individually scaled with temperature. Both phases contribute to the overall duration of the cell cycle, and the durations of the two phases are largely determined by different processes, cyclin synthesis for the former and cyclin degradation for the latter. Due to the opacity of the *Xenopus* embryo, it is difficult to assess these cell cycle phases by *in vivo* microscopy. We therefore turned to cycling *Xenopus* egg extracts, which are transparent and highly amenable to microscopy.

Cycling extracts were prepared and supplemented with a Cdk1 FRET sensor, whose emission increases upon Cdk1 activation and/or inactivation of opposing phosphatase(s) (78). Unlike the original PBD (polo-box domain)-based sensor used in human cells (79) and *Drosophila* embryos (80), this redesigned WW-based sensor uses a Cdc25C substrate motif, WW phospho-binding domain, and EV linker to enhance signal-to-noise performance in *Xenopus* extracts (78). The extracts were then encapsulated in oil droplets, as previously described (81, 82) (Fig. 4A). The encapsulated extract droplets were then loaded into Teflon-coated imaging chambers, which were immersed in mineral oil and placed on a microscope stage equipped with a custom Peltier element-based heating/cooling device, similar to a setup tailored for embryos (44). The FRET sensor enabled real-time visualization of oscillations in Cdk1 activity in hundreds of droplets situated at different positions within the temperature gradient (Fig. 4A, Supplemental Video 7). As shown in Fig. 3C (top), Cdk1 activity first rose slowly (phase 1), then spiked to high levels (phase 2), then fell to low levels to allow a new cycle to begin (phase 3). The three phases of the Cdk1 activity cycle correspond well to the phases seen by direct biochemical assays of Cdk1 activities in cycling extracts (62, 83, 84), and to the phases of Cdk1 activation and inactivation seen in the computational models (Fig. 3C, bottom). The cell cycle was found to proceed most rapidly at temperatures of around 25°C, and to slow down at both colder and warmer temperatures (Fig. 4B-C). One unanticipated finding was that the cell cycle proceeded fairly normally at temperatures as high as 32°C, even though in intact embryos, temperatures above 28°C typically killed the embryos and halted the cell cycle. This allowed us to probe a wider range of temperatures in extracts than was possible *in vivo*. The period of the extracts’ cell cycles increased over time, consistent with previous findings (39, 78, 81, 85). In addition, the overall response of the cell cycle to temperature was consistent across different biological samples and experimental days (Fig. S7). We confined our analysis to cycles 2–4, characterizing the duration of each cycle in individual droplets (Fig. S8A-C) and after pooling (Fig. 4C-D). Similar trends could be seen in both the individual and pooled data (Fig. 4C-D). Alternatively, we analyzed all cell cycles that occurred during the first 300 minutes rather than the first three cycles. This procedure yielded essentially identical results (Fig. S8D).

**Fig. 4:**
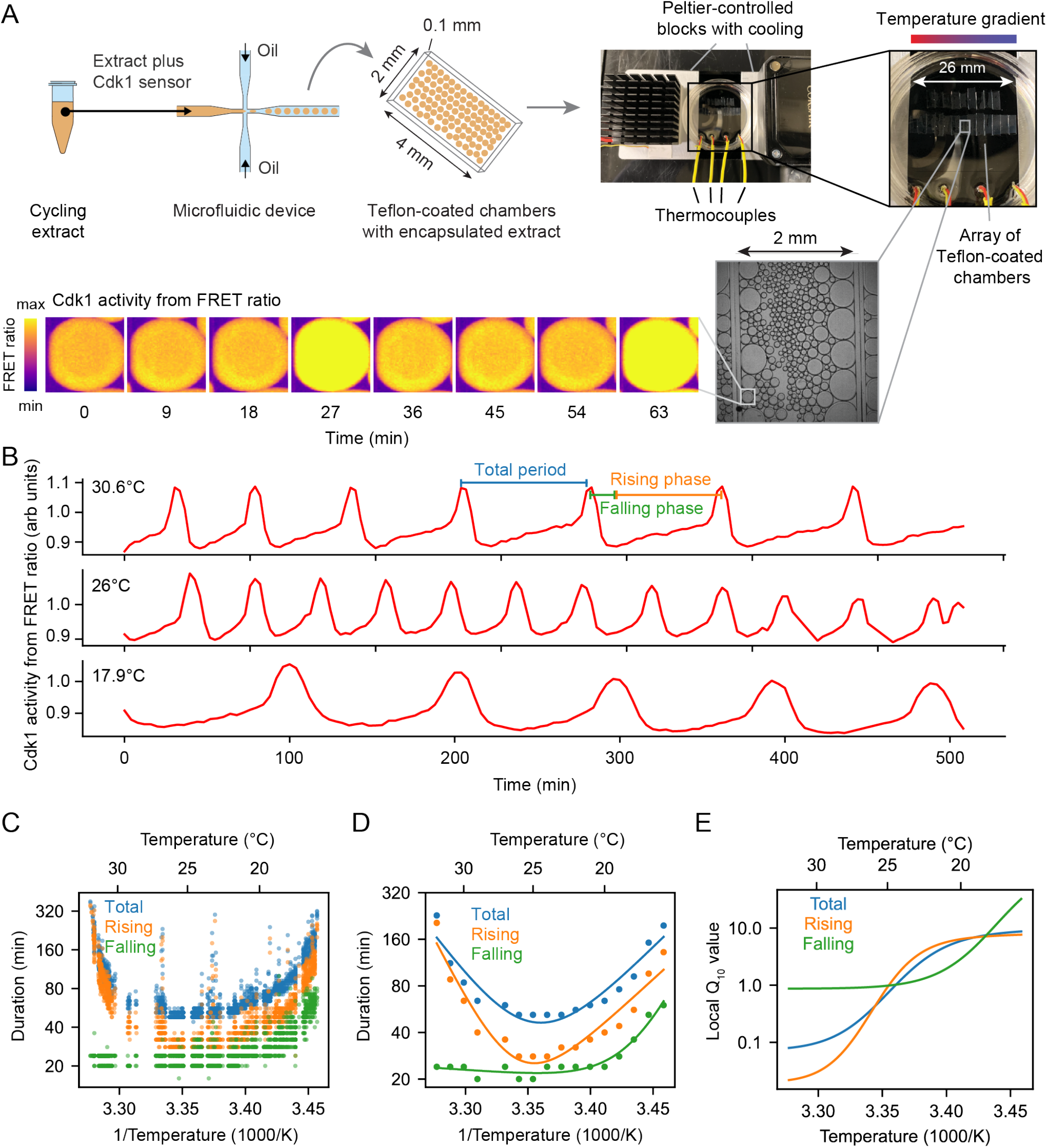
The durations of interphase and M-phase scale differently and non-Arrhenius with temperature in cycling frog egg extracts. A. Sketch of the setup to encapsulate cycling frog egg extracts in droplets surrounded by oil, including pictures of the customized device to control temperature of extract droplets with snapshots of measured FRET ratios in an example droplet. B. Representative time series of measured FRET ratios at different temperatures. C. Analysis of the duration of the total cell cycle (blue), the rising phase (orange), and the falling phase (green) in function of temperature. D. Median per temperature bin (rounded to integers) of the data shown in panel C. Optimal fits using a double exponential function are overlayed. E. Local *Q*_10_-value as function of temperature, calculated from the fitted double exponential function.

We analyzed time series data from hundreds of droplets at temperatures from 16°C to 32°C, and plotted the temperature dependence of the cell cycle durations as well as the durations of the rising and falling phases, which correspond approximately to interphase-through-metaphase and metaphase-through-mitotic exit, respectively. This analysis showed that both the total cell cycle duration and the duration of the rising phase exhibited a U-shaped dependence upon temperature—these durations decreased steeply as the temperature rose from 16°C to 20°C, then plateaued, and then increased steeply at temperatures above 30°C (Fig. 4C-D). In contrast, the duration of the falling phase decreased with temperature and then plateaued beginning at about 20°C, but did not slow down to a measurable extent at higher temperatures. These trends can be seen both from the raw data (Fig. 4C) and from binned, averaged data (Fig. 4D). Thus, the rising phase, whose duration is mainly due to the rate of cyclin synthesis, and the falling phase, whose duration is mainly due to APC/C activity, are differently affected by temperature. Interestingly, while the durations of transitioning into and out of M phase increased at low temperatures, the duration of mitotic exit remained constant at high temperature (Fig. S9). Conversely, at high temperatures, the duration of mitotic entry substantially increased.

Double exponential curves, which assume a biphasic dependence of enzyme activity upon temperature, accounted for the shapes of the Arrhenius plots (Fig. 4D). We computed local *Q*_10_ values from the fitted curves, which revealed significant changes with temperature (Fig. 4E). Generally, the *Q*_10_ for total cell cycle duration was close to that of rising phase duration (Fig. 4E), underscoring the fact that interphase constitutes a majority of the cell cycle (Fig. 4B).

### The non-Arrhenius scaling results from both the biphasic temperature sensitivity of cyclin synthesis and an imbalance in the Arrhenius constants for cyclin synthesis and degradation

We next asked how well the 2-ODE computational model could account for how the Cdk1 activity cycle varied with temperature in cycling extracts, and whether the scaling of the activation energies for key regulatory processes (*k*_*s*_, *k*_*d*_, *ϵ*) could be inferred. We utilized the temperature dependence of the measured durations of the rising and falling phases of the Cdk1 time series (Fig. 4D) to optimize our computational model. Employing the approximate Bayesian computation method (details in Supplementary Note 3.D and Figs. S10-S11), we sought optimal values that described the scaling curves for cyclin synthesis rate (*k*_*s*_), cyclin degradation rate (*k*_*d*_), and time scale separation (*ϵ*), which relates to Cdk1 activation (*a*) and inactivation (*i*). The temperature dependence of each these parameters is described by a double-exponential scaling curve (for details, see Supplementary Note 3.D). Leveraging sequential Monte Carlo sampling, the method gradually improved fits to the data. Rather than a single optimal value, the method produces a distribution of well-fitting parameters (gray lines in Fig. 5A, with the optimal fit highlighted in color). The temperature dependence of the fitted model parameters (Fig. 5B) revealed significant temperature-induced changes in cyclin synthesis rate (*k*_*s*_) and time scale separation (*ϵ*), up to five-fold across the temperature range, whereas changes in the cyclin degradation rate (*k*_*d*_) were much smaller (see Figs. S10–S11 for the distributions of the parameters yielding good fits). Additionally, the temperature dependence of *k*_*d*_ was well described by a single Arrhenius equation, while *k*_*s*_ and *ϵ* required a double exponential function for accurate description. Figs. S10–S11 further demonstrate that the model successfully captures the observed changes in oscillation dynamics as long as the cyclin synthesis rate is more temperature-sensitive than the degradation rate. This means that the fitted values of these activation energies are not tightly constrained by the data. Specifically, the model remains consistent with the data as long as the cyclin synthesis rate increases more steeply with temperature than the degradation rate. This is supported by simulation results and by the ABC-inferred parameter distributions (Figs. S10–S11), which show that the apparent activation energy for synthesis, *E*_*a*_(*k*_*s*_), is centered around 113kJ/mol. In contrast, *E*_*a*_(*k*_*d*_) for degradation is very broadly distributed, and although the peak is at around 12kJ/mol, the mean is closer to 49kJ/mol. This suggests that a range of degradation temperature sensitivities is compatible with the data, provided synthesis remains more sensitive.

**Fig. 5:**
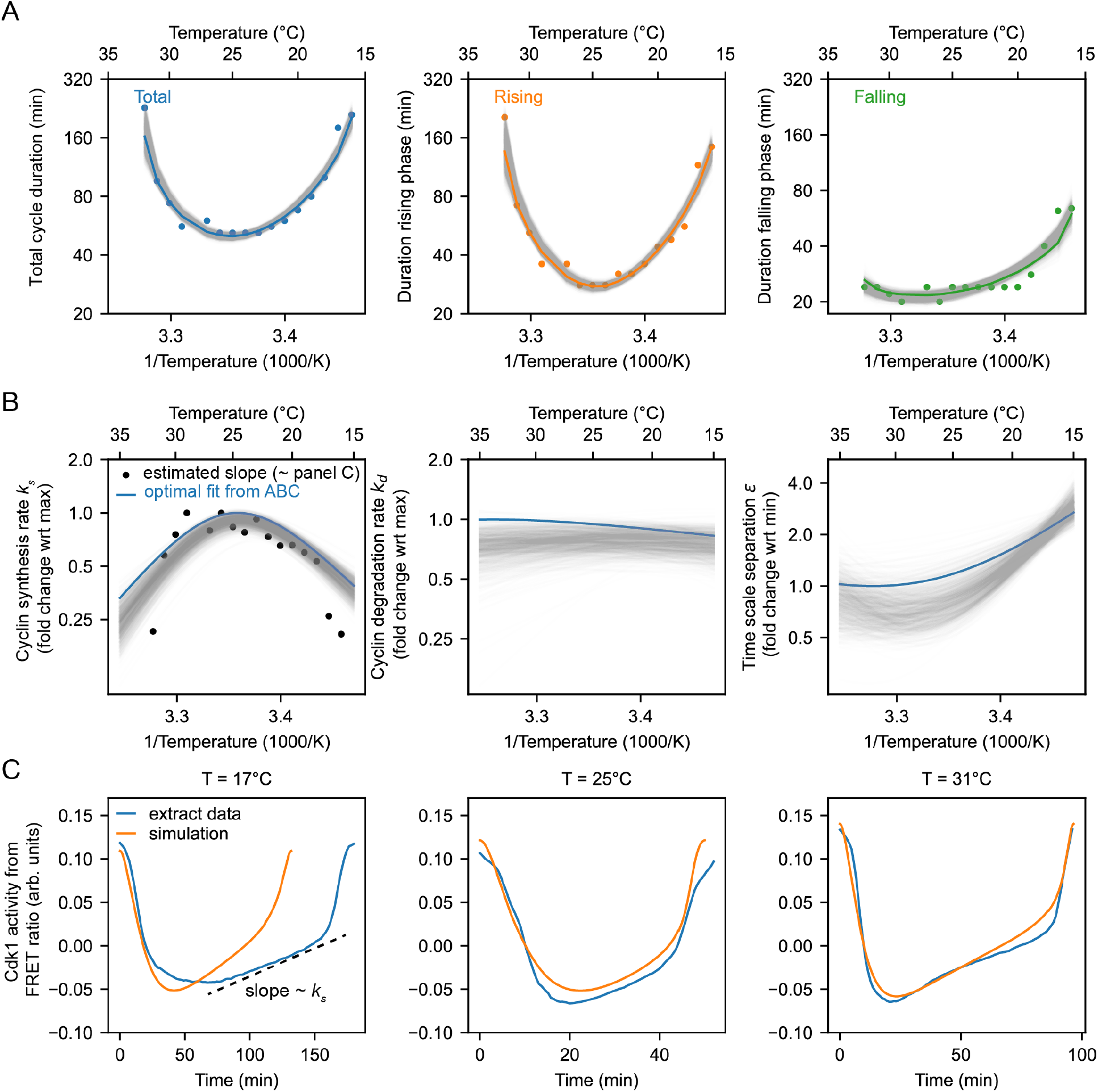
Non-Arrhenius scaling as a result of biphasic cyclin synthesis rate and an imbalance in the temperature scaling of cyclin synthesis and degradation. A. Using the ABC algorithm, we minimize the mean square error (MSE) between the measured and simulated (using the two-ODE model) durations of rising phase and falling phase. The measurements are shown with the dots, while the gray lines show results from the model with the colored line the best fit (smallest MSE). B. Optimal temperature scaling of parameters, i.e. the cyclin synthesis rate, the cyclin degradation rate, and the time scale separation, resulting from the ABC algorithm as shown in A. The black dots correspond to the cyclin synthesis rate *k*_*s*_ (nM/min) directly estimated from the FRET ratio time series (Fig. S12 and Supplementary Note 4). C. Blue line: averaged time series of Cdk1 activity (measured FRET ratio) at three different temperatures (*T* = 17°C, *T* = 25°C, *T* = 31°C). See Fig. S13 and Supplementary Note 5 for the method to compute the average waveform. Orange line: time series of the computational model, computed using the optimal parameter scaling shown in Panel B.

The three phases of the Cdk1 activity cycle correspond well to the phases seen by direct biochemical assays (61, 62, 84), and they represent the accumulation of low activity cyclin-Cdk1 complexes (phase 1), followed by the activation of Cdc25 and the inactivation of Wee1 and PP2A-B55 (phase 2), and finally the APC/C-Cdc20-mediated degradation of cyclin and re-activation of PP2A-B55 (phase 3). During the first phase, both cyclin levels and Cdk1 activity increase approximately linearly over time (61, 62) (Fig.5B-C). To obtain an independent estimate of how *k*_*s*_ varies with temperature, we computed the slope of the interphase segment of the Cdk1 FRET time series (Fig.5C, dashed line; Supplementary Note 4, Fig. S12). These empirical slopes revealed scaling trends consistent with those obtained from model fitting via the ABC algorithm (Fig. 5B, overlaid black dots), reinforcing the robustness of the inferred biphasic temperature dependence. While the Cdk1 FRET signal primarily reflects cyclin accumulation, it could in principle also be influenced by other regulatory processes, such as phosphatase inactivation or post-translational modifications that modulate Cdk1 activity independently of cyclin levels. Therefore, fitting the slope of the FRET signal does not necessarily isolate cyclin synthesis alone. Nonetheless, the strong agreement between the model’s predictions and the slope-based estimates suggests that the FRET signal serves as a useful proxy for cyclin synthesis over this regime. This interpretation is further supported by previous studies showing that cyclin synthesis is the primary driver of Cdk1 activation during interphase. Pomerening *et al*. demonstrated that both cyclin levels and Cdk1 activity rise approximately linearly with time throughout interphase (61), consistent with a model in which increasing concentrations of phosphorylated, pre-activated cyclin–Cdk1 complexes underlie the gradual activation of the oscillator. Moreover, other key regulators of Cdk1, including Cdc25C and Wee1, exhibit minimal changes in their abundance or phosphorylation state during interphase and only transition into their mitotic forms immediately before mitotic entry (see Fig. 7 in (62)).

As a final check of our fitting procedure, we computed the average shape of the Cdk1 time series experimentally (from the FRET signal, Fig. S13, Supplementary Note 5), and compared it to the model’s predicted time series, given the fitted scaling (Fig. 5B). The simulated time series closely recapitulated the experimental oscillations (Fig. 5C). The optimization was done only on durations. Thus, the match between simulated and experimentally observed waveforms provides another argument that the fitted scaling curves for the rates explain the scaling observed in the droplets.

In summary, the comparison of apparent activation energies highlights the greater temperature sensitivity of cyclin synthesis rate compared to cyclin degradation rate. This sensitivity aligns with scenarios predicted to yield non-Arrhenius scaling across a wide temperature range (Fig. 3D, Case 3). Furthermore, experimental findings indicate that cyclin synthesis rates decreased at elevated temperatures, corroborating another scenario leading to non-Arrhenius scaling (Fig. 3D, Case 4). Our analysis indicates that both mechanisms contribute to the non-Arrhenius scaling properties of the early embryonic cell cycle oscillator.

### *In vitro* enzyme assays confirm the imbalance in the cyclin synthesis and degradation *E*_*a*_ values

To further test the inference that cyclin synthesis and degradation scale differently at the low end of the temperature range, we carried out direct measurements of the two processes in X. laevis frog egg extracts (86) (Fig. 6A–B). The synthesis of one mitotic cyclin, cyclin B2, in cycling extracts was monitored by quantitative Western blotting, using the cyclin B2 levels present in CSF extracts as a normalization standard (Fig. S14). APC/C activity was gauged by introducing securin-CFP, translated in wheat germ extracts, as a fluorescent reporter of APC/C activity into CSF extracts, then driving the extracts out of CSF arrest with calcium plus cycloheximide and into mitotic arrest with non-degradable cyclin B (Fig. S15). Experimental protocols are detailed in Supplementary Note 6. These measurements were conducted across temperatures ranging from 16°C to 26°C and did not extend into the high temperature range where the rate of cyclin synthesis as inferred in Fig. 5B began to drop. These rate data were consistent with the Arrhenius equation (Fig. 6B, green line), and cyclin synthesis was more sensitive to temperature than cyclin degradation, with fitted apparent Arrhenius energies of 87 and 51 kJ/mol, respectively. Bootstrapping supported the statistical significance of this difference (Fig. 6C). This provides direct support for the hypothesis that the different scaling of opposing enzymes contributes to the non-Arrhenius character of the cell cycle period.

**Fig. 6:**
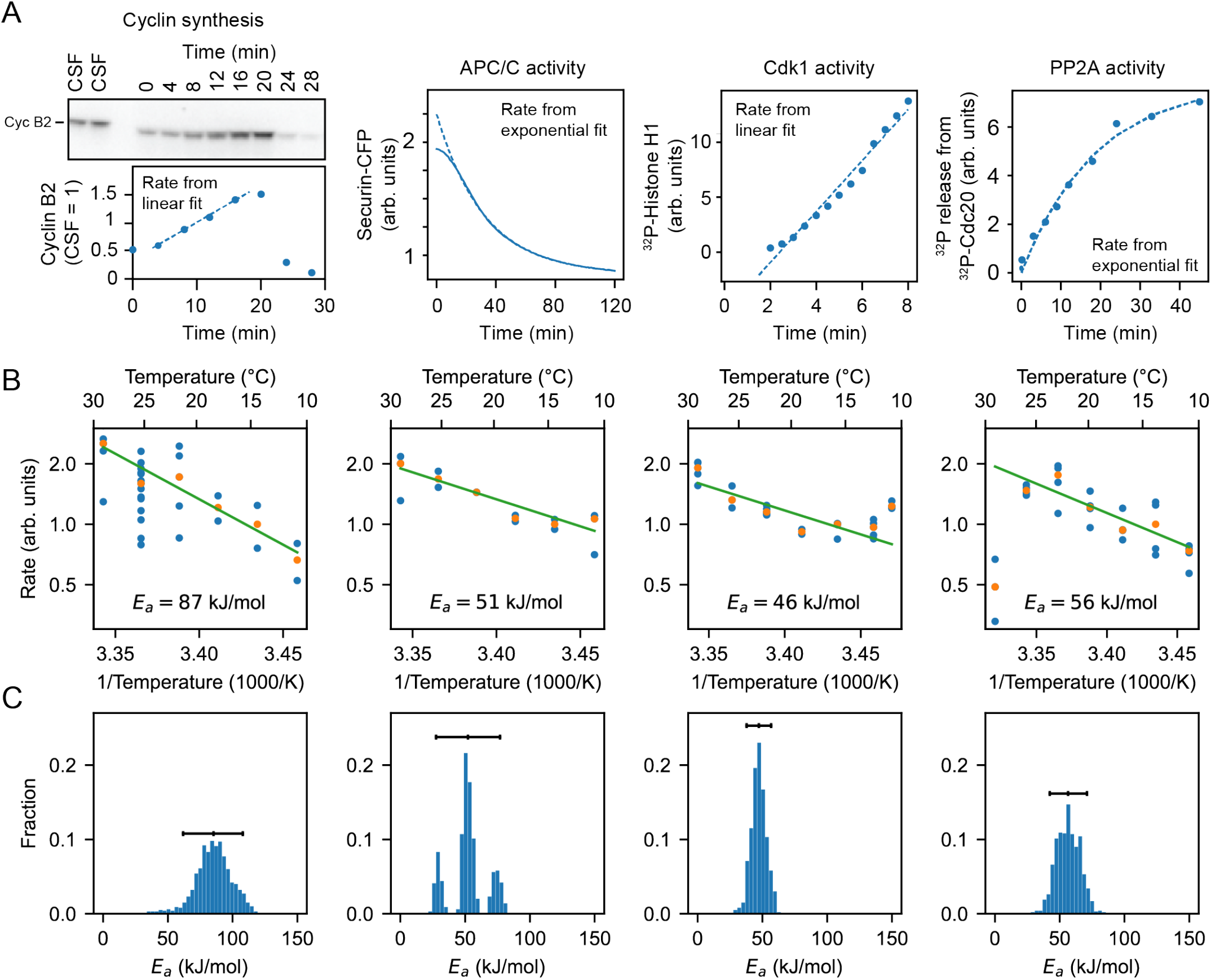
Frog egg extract measurements reveal temperature dependence of cell cycle regulators. A. Examples for how the rates for Cyclin B synthesis, APC/C activity, Cdk1 activity, and PP2A activity were fitted from time series of different biochemical assays using frog egg extracts at constant temperatures (here for *T* = 24°C), see Supplementary Note 6. B. The assays were repeated for temperatures in the interval 16 − 26°C, and (apparent) activation energies were extracted. Blue dots represent data of individual fitted time series, while the orange dots are the medians per temperature. C. Probability distribution of fitted (apparent) activation energies using bootstrapping with 90% confidence intervals (see Supplementary Note 2 for details on the bootstrap procedure).

**Fig. 7:**
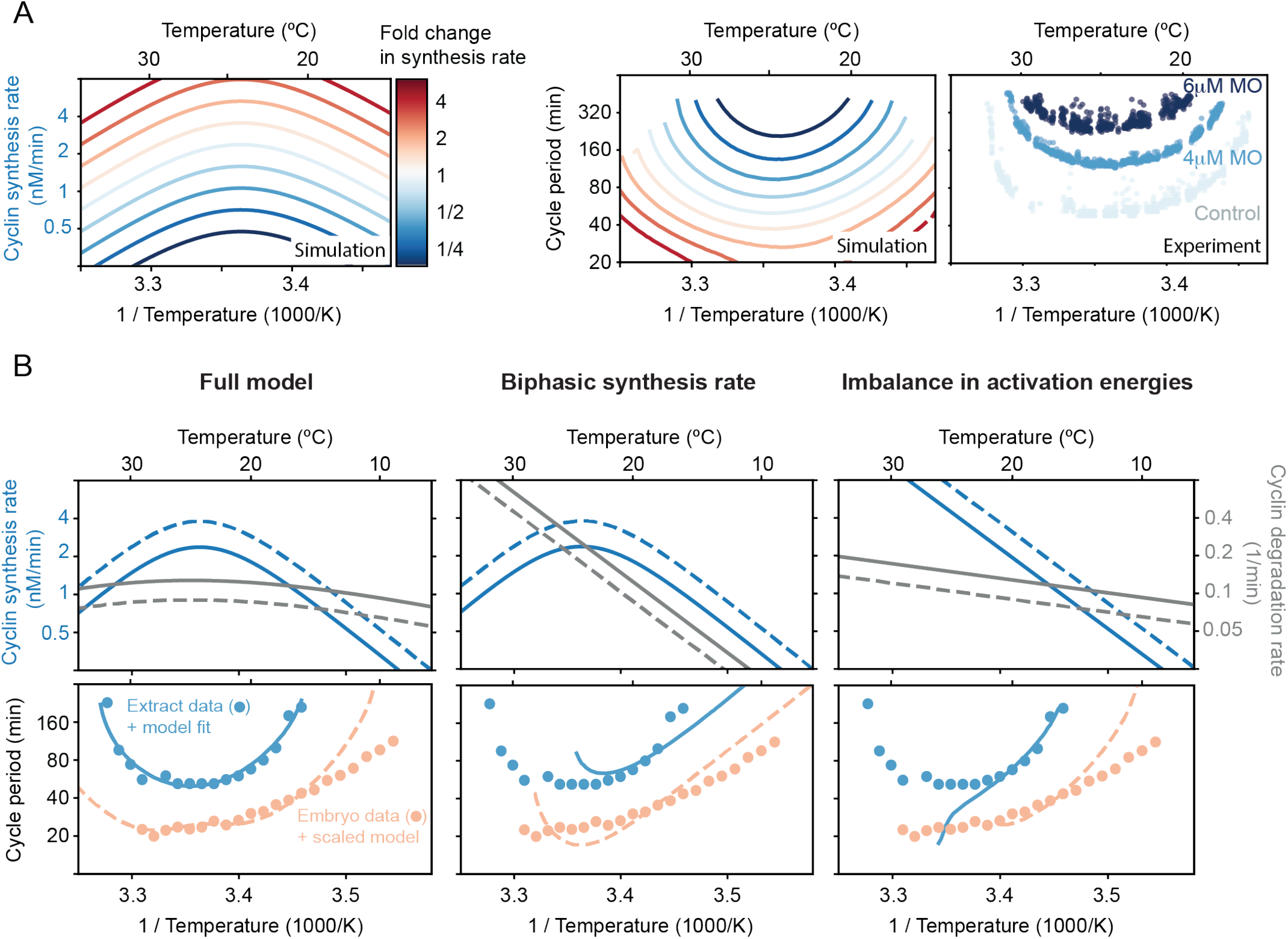
Decreasing the cyclin synthesis rate decreases the viable temperature range. A. Influence of changing the basal cyclin synthesis rate by a factor up to 5 on the shape of the temperature response curves. The degradation rate is scaled up to a factor of 3. The two left panels show simulations of the 2-ODE model of the cell cycle oscillator using a parameter set obtained from the ABC method (one of the gray lines in Fig. 5B), plotting the temperature-dependence of the cyclin synthesis rate and the corresponding cell cycle period. The right panel shows the cell cycle duration as a function of temperature obtained from encapsulated extracts with 0, 4, or 6 µM morpholino (MO) oligonucleotides against isoforms of Xenopus cyclin B1/B2 mRNA species, thus lowering the cyclin synthesis rate. B. Different scenarios in temperature dependence of cyclin synthesis and degradation lead to different non-Arrhenius scaling of cell cycle oscillations. While a biphasic cyclin synthesis rate leads to a double exponential response curve, the imbalance in activation energies introduces a curved non-Arrhenius response at lower temperatures, which is critical for reproducing the experimental data measured in frog egg extract.

We also measured the temperature dependence of two other key cell cycle regulators, cyclin B-Cdk1 and PP2A-B55 (Figs. S16,S17). These opposing enzymes are critical for the phosphorylation and dephosphorylation of many cell cycle proteins, and their activities would be expected to contribute to the dynamics of mitotic entry and mitotic exit. Fig. 6 suggests minor variations in their temperature sensitivity, with apparent Arrhenius energies of 46 and 56 kJ/mol, compatible with robust functioning of the cell cycle oscillator over its nominal temperature range.

### Decreasing the cyclin synthesis rate decreases the viable temperature range

Our analysis suggests that the failure of the *Xenopus* embryonic cell cycle oscillator at temperature extremes is governed by two distinct mechanisms. At high temperatures, oscillations break down due to the biphasic temperature dependence of cyclin B synthesis: the synthesis rate declines with increasing temperature and eventually becomes insufficient to sustain oscillations in the face of baseline degradation. At low temperatures, failure arises from an imbalance in Arrhenius scaling. Specifically, the higher activation energy of synthesis relative to degradation causes the synthesis rate to become too low to counteract degradation. Thus, the temperature range over which the oscillator functions is determined by the temperature-dependent interplay of these two opposing processes, both essential for cell cycle progression.

This reasoning predicts that modulating the overall cyclin synthesis rate should systematically alter the temperature range over which the oscillator can operate. To test this, we used our 2-ODE model of the cell cycle oscillator using a parameter set obtained from the ABC method (one of the gray lines in Fig. 5B). We then systematically varied the synthesis rate at a reference temperature while preserving its temperature dependence. At the same time we allowed the degradation rate to be scaled similarly, but to a lesser extent. In Fig. 7A we show one representative example (see more simulation details in Fig. S18). The resulting temperature-period curves were U-shaped and shifted in a consistent, and perhaps non-intuitive, way: increasing the synthesis rate led to faster oscillations and broader temperature ranges, while decreasing it caused both upper and lower temperature bounds to move inward. At high synthesis rates oscillations failed, starting at intermediate temperatures.

We next tested this prediction experimentally by titrating cyclin B morpholino antisense oligonucleotides (0, 4, or 6 *µ*M) into encapsulated extracts subjected to a temperature gradient (see Materials and Methods). These morpholinos inhibit cyclin B translation by binding to its mRNA. As predicted, increasing morpholino concentrations resulted in longer minimum cycle periods and a narrower viable temperature range (Fig. 7A), consistent with the model. A quantitative discrepancy remained, with experimental oscillations persisting at more extreme temperatures than predicted. This could reflect additional regulatory layers that are not captured in our minimal model, beyond cyclin synthesis/degradation and Cdk1 (in)activation. Moreover, the cell cycle oscillator in extracts is also gradually slowing down over time (see Fig. 4B), which could also explain the discrepancy between the experiments and the idealized, time-invariant simulations.

We then asked whether the same model, tuned to cycling extract data, could recapitulate the cell division timing in early *X. laevis* embryos (Fig. 7B). While the general shape was similar, oscillations in embryos were systematically faster than in extracts. This could be due to cytoplasmic dilution during extract preparation — though such effects are modest at moderate dilutions (87) — or the absence of nuclear and membrane components. The period differences between embryos and extracts were also similar to variations among extracts from different biological replicates (Fig. S7). We therefore rescaled the cyclin synthesis by a factor of 1.6 and the degradation rates by 0.7 to match cycle duration at 25°C. Remarkably, this single-point scaling allowed the model to capture the embryo temperature response curve with reasonable accuracy across a broader temperature range than would be achieved by simply scaling the extract response curve. (Fig. 7B, bottom left). The embryo data did deviate at lower temperatures, suggesting additional embryo-specific dynamics not captured by our simple model.

Finally, we asked whether either mechanism, biphasic synthesis or an imbalance in Arrhenius scaling exponents, could alone account for the observed temperature dependence. Using our calibrated model, we isolated each effect and found that both independently produce non-Arrhenius temperature scaling (Fig. 7B, middle-right). However, neither mechanism alone reproduced the fine structure of the experimental curves, indicating that both are required to explain the wide temperature adaptability of the embryonic cell cycle oscillator. In particular, a biphasic cyclin synthesis rate leads to a double exponential response curve, capturing deviations at high temperatures, but still appears Arrhenius-like at low temperatures. Especially for the extract data, such a model fails to capture the strong low-temperature deviations. In contrast, the imbalance in activation energies introduces a curved non-Arrhenius response across all temperatures, yet it fails to capture the sharp increase in period at high temperatures.

Taken together, these findings support our model, which attributes the non-Arrhenius scaling properties of the early embryonic cell cycle oscillator to two concurrent mechanisms. At low temperatures, the scaling is primarily driven by an imbalance between opposing cyclin synthesis and degradation rates. In contrast, at high temperatures, the biphasic nature of cyclin synthesis plays a critical role in capturing the upward curvature of the oscillation period.

## Discussion

Previous work suggested that the early embryonic cell cycle scales similarly with temperature in several organisms (30–33). Here we have extended these measurements to *Xenopus tropicalis* and *Danio rerio*, and have supplemented previous work on *Xenopus laevis* with additional types of measurements. We found that although the periods of the cell cycle at the organisms’ nominal temperatures vary from about 5 min for *C. elegans* and *C. briggsae* to about 25 min for *X. laevis*, the temperature scaling of the periods is quite similar. The apparent Arrhenius energies averaged 75 ± 7 kJ/mol (mean ± std. dev., n = 6), and the average *Q*_10_ value at 20°C was 2.8 ± 0.2 (mean ± std. dev., n = 6) (Figs. 1-2). In all cases the periods deviated from the Arrhenius relationship at high temperatures, and for *X. laevis*, the Arrhenius plots were non-linear throughout the range of permissible temperatures.

In some ways it is perhaps not surprising that the temperature scaling data could be approximated reasonably well by the Arrhenius equation. Crapse et al. (32) have shown computationally that chaining together a sequence of chemical reactions results in only minor deviations from ideal Arrhenius scaling if one assumes that the individual enzymes’ activation energies do not differ greatly. Experiments have shown that Min protein oscillations, crucial for bacterial cell division, also display Arrhenius-like scaling behaviors (88). The classic chemical oscillator, the Belousov-Zhabotinsky reaction, approximately obeys the Arrhenius equation (89–91), and in general, many biological processes at least approximately conform to the Arrhenius equation or one of the proposed modified versions of the Arrhenius equation (15–22).

These observations notwithstanding, it was not obvious to us why a complex oscillator circuit, with non-linearities and feedback loops, should yield even approximately Arrhenius temperature scaling, and what the origins of the experimentally-observed deviations from Arrhenius scaling might be. Through modeling studies we identified two plausible mechanisms for the observed non-Arrhenius behavior: an emergent mechanism resulting from differences in the Arrhenius energies of opposing enzymes in the network (Fig. 3D, Case 3), and a biphasic temperature dependence for one or more of the critical individual enzymes (Fig. 3D, Case 4). A priori, either or both of these mechanisms could pertain.

Experimental observations combined with model-based inference suggest that a key step in the oscillator circuit — the synthesis of the mitotic cyclin protein — exhibits a strongly biphasic dependence on temperature. While intact *Xenopus* embryos do not survive above 29°C, cell-free cycling extracts can continue oscillating at temperatures exceeding *≈* 30°C. Above 30°C, the rate of cyclin synthesis (as inferred from the Cdk1 activity sensor) and the rate of progression through interphase clearly decrease with increasing temperature, whereas at lower temperatures they increase with increasing temperature (Fig. 4). Our hypothesis is that above some maximum permissible temperature, the imbalance between the cyclin synthesis and degradation rates causes the oscillator to fail and the cell cycle to arrest.

Cyclin synthesis and degradation also scaled differently with temperature at the low end of the permissible temperature range. This was inferred from fitting the parameters of the two-ODE model to the experimental data (Fig. 5), and then was directly shown by *in vitro* assays of cyclin synthesis and degradation (Fig. 6). This means that below a critical temperature, cyclin synthesis and degradation should again be out of balance, causing oscillations to cease.

To further test this hypothesis, extracts were treated with a mixture of four morpholino oligonucleotides, two for cyclin B1 and two for cyclin B2, to inhibit cyclin translation enough to slow but not block the cell cycle at normal temperatures (see Materials and Methods). We asked whether this decreased the maximum permissible temperature, raised the minimum permissible temperature, or both. We found that both temperature limits were similarly affected, and the operating range of the cell cycle oscillator was narrowed, as predicted by our simple two ODE model (Fig. 7). This finding is consistent with the hypothesis that both operating limits are determined by the balance between cyclin synthesis and degradation.

One question then is why did evolution not arrive at a system where cyclin synthesis and degradation did not go out of balance, at high and low temperatures? We suspect that there are trade-offs between competing performance goals for the oscillator and its components. Perhaps the molecular flexibility required to make protein synthesis run as fast as possible at the temperatures typically experienced by an ectotherm render the ribosomes vulnerable to unfolding at slightly higher temperatures. Likewise, cyclin synthesis and degradation might work best at normal temperatures even if their temperature scaling does not perfectly match the overall system’s activation energy, suggesting that the observed *E*_*a*_ reflects a trade-off between fast reaction rates and ideal scaling.

While our study focuses on early embryonic systems that are largely transcriptionally silent, recent work in yeast (92) has shown that temperature-induced changes in gene expression can drive fate decisions, and synthetic bacterial circuits have been engineered to achieve temperature compensation through specific protein modifications (93). These findings highlight complementary mechanisms of thermal adaptation, from network-level emergent scaling, as we demonstrate here, to dedicated molecular adaptations. Furthermore, synthetic gene circuit evolution studies (94) offer promising opportunities to explore how temperature robustness can arise in engineered systems, providing a future experimental platform to test and extend the principles uncovered here.

One final question is how the behaviors seen here compare to those of the same circuit in endotherms, organisms that have at great metabolic cost freed their biochemistry from needing to function reliably over such wide temperature ranges. Although the four enzymatic processes individually assessed here (cyclin synthesis, cyclin degradation, Cdk1 activity, and PP2A-B55 activity) differed in their temperature scaling, they did not differ by that much; their *E*_*a*_ values averaged to 60 kJ/mol with a standard deviation of 16 kJ/mol or 27%. It seems plausible that endothermy might allow enzymes with a wider range of activation energies to be used than would be possible in ectotherms.

## Materials and Methods

### Xenopus egg extract

Cell-free cycling extracts and CSF extracts were made from *Xenopus laevis* eggs following a published protocol from Murray (95). For cycling extracts, this protocol was adapted as in (81). Extracts for Fig. 3B-G and Fig. 4 were then supplemented with 1µM Cdk1-FRET sensor, as described in Maryu and Yang (78), and also with 1X energy mix (7.5 mM Creatine phosphate, 5mM ATP, 1mM EGTA, 10 mM MgCl2). Work from the Yang lab demonstrated that an intermediate range of dilution of the extracts can improve the number of cycles, with the best activity at around 30% dilution (87). As a result, for the data described here, the dilution was kept constant at 30% with extract buffer (100 mM KCl, 0.1 mM CaCl2, 1 mM MgCl2, 10 mM potassium HEPES, 50 nM sucrose, pH 7.8). Extracts for the biochemical assays in Fig. 3A were undiluted.

The extract was encapsulated via a water-in-oil emulsion using a micrufluidic device. The fabrication of the device and droplet generation followed a previously published protocol (96). Briefly, cycling extract (water phase) was mixed with 2% 008-FluoroSurfactant in HFE7500 (Ran Biotechnologies, Inc.) (oil phase) inside a microfluidic device driven by an Elveflow OB1 multi-channel flow controller (Elveflow). Air pressure was 2 psi for both the extract and oil channels. After droplets were generated, they were loaded into VitroCom hollow glass tubes with a height of 100 µm (VitroCom, 5012) pre-coated with trichloro (1H,1H,2H,2H-perfluorooctyl) silane, and then immersed into a glass-bottom dish (WillCo Wells) filled with heavy mineral oil (Macron Fine Chemicals) to prevent evaporation.

### Temperature gradient generation

A custom plastic microscope stage was fabricated to fit two aluminum plates on each side of the imaging dish. Each aluminum plate was attached to a TEC1-12706 40*40MM 12V 60W Heatsink Thermoelectric Cooler Cooling Peltier Plate (HiLetGo) using thermal conductive glue (G109, GENNEL). The plate designated for temperatures above room temperature had an additional heatsink (40mm x 40mm x 20mm, black aluminum, B07ZNX839V, Easycargo) and a cooling fan to improve performance. The plate designated for cold temperatures had an additional liquid cooling system (Hydro Series 120mm, CORSAIR) attached with termal conductive glue.

Peltier devices were controlled via two CN79000 1/32 DIN dual zone temperature controllers (Omega). In all experiments, the target temperature was set to 65°C and 1°C for the hot and cold plates respectively. With both plates on, it was always the case that the hot plate reached its target temperature and stayed constant within 5-10 min and the cold plate stayed stable at 10°C.

The imaging dish was attached to the aluminum plates with Thermal adhesive tape 2-5-8810 (DigiKey) to ensure proper thermal conduction. Temperature was logged via 4 K-Bead-Type thermocouples placed on the imaging dish touching the bottom surface. Data was acquired using a 4 Channel SD Card Logger 88598 AZ EB (AZ Instruments). Room temperature was also captured using the same method via a thermocouple attached to the microscope stage.

### Western blotting

Cycling extracts were prepared according to the method by Murray et al. (95), except that eggs were activated with calcium ionophore A23187 (5 µl of a 10 mg/ml stock of A23187 in 100 ml 0.2x MMR) rather than with electric shock. After preparing the extracts, they were distributed to several eppendorf tubes and brought to a specified temperature between 16 and 26°C within 20 minutes. 2 µl samples were taken every 4 minutes (every 8 minutes at 16°C) and immediately frozen on dry ice. To each 2 µl aliquot, 48 µl of SDS sample buffer supplemented with DTT was added, and the samples were boiled during 10 minutes at 95°C. 12 µl of the cycling extract samples and 4 µl of the reference samples (CSF extract prepared according to the method by Murray et al. (95), were run on 10% Criterion TGX Precast protein gels and transferred to a PVDF membrane using the Bio-Rad Trans-blot Turbo system. After blocking in milk (4% w/v in TBST), the blots were incubated with a 1/500 dilution of anti-cyclin B2 antibody (X121.10, Santa Cruz) overnight at 4°C followed by a 1/10.000 dilution of anti-mouse IgG HRP-linked whole secondary antibody (GE Healthcare NA931), during 1 hour at room temperature. Finally, the blots were developed using Supersignal West Femto chemiluminescent substrate. The images of the blots can be found on the Zenodo repository (doi: 10.5281/zenodo.15591678).

### Time-lapse fluorescence microscopy

For Figs. 4, 5, 7, imaging was carried out on an inverted Olympus IX83 fluorescence microscope with a 4× air objective, a light emitting diode fluorescence light source, a motorized x-y stage, and a digital complementary metal–oxide–semiconductor camera (C13440-20CU, Hamamatsu). The open-source software µManager v1.4.23 was used to control the automated imaging acquisition. Bright-field and multiple fluorescence images of CFP, FRET, and YFP were recorded at a frequency of one cycle every 3 to 7 min for 40 to 50 hours for each sample.

### Image processing and analysis methods

Grids of images were captured and subsequently stitched together using ImageJ’s Grid/Pairwise Stitching plug-in, in conjunction with additional pipeline code written in Fiji/Java. Bright-field images from the first frame were used to generate stitching parameters, which were fed to ImageJ to stitch each channel at each frame consecutively. The FRET ratio was calculated as in Maryu and Yang (78).

For Figs. 4, 5, 7, custom scripts in MATLAB 2020a and ImageJ were written to perform image processing. Briefly, each microscope position was processed by manually selecting the region containing the tube of interest and then algorithmically cropping and resizing that region in all channels. Then, bright-field images were used for individual droplet segmentation and tracking using Trackmate 7.12.1. Only individual droplets whose radius was smaller than 100 µm and track started within the first 60 min of the experiment were selected for further analysis. FRET ratio intensity peaks and troughs were first auto-selected and then manually checked and corrected using custom Python scripts. Rising and falling periods were calculated from this data. All code is available at https://github.com/YangLab-um/temperature and https://github.com/YangLab-um/dropletDataProcessing. The tracking data can be found on the Zenodo repository (doi: 10.5281/zenodo.15591678)

### Morpholino oligonucleotides

A combination of four morpholino antisense oligonucleotides (Gene Tools, LLC) at equal concentrations were designed and their sequences are as follows:

- Morpholino anti-Xenopus-CyclinB1 (ccnb1_a): ACATTTTCCCAAAACCGACAACTGG
- Morpholino anti-Xenopus-CyclinB1 (ccnb1_b): ACATTTTCTCAAGCGCAAACCTGCA
- Morpholino anti-Xenopus-CyclinB2 (ccnb2_l): AATTGCAGCCCGACGAGTAGCCAT
- Morpholino anti-Xenopus-CyclinB2 (ccnb2_s): CGACGAGTAGCCATCTCCGGTAAAA

We applied a total concentration of 0, 4, or 6 µM the morpholino cocktail to the cycling *Xenopus* extracts to suppress the endogenous translation of cyclin B1/B2. These concentrations were chosen within a dynamic range that could inhibit cyclin translation but should not terminate the cell cycle at normal temperatures, based on microfluidic channel tuning experiments (97).

### Fitting of scaling laws

For the fitting of Arrhenius and other functional forms, we always binned the data per integer temperature value, and then took the median per temperature. This results in a dataset with one rate/duration per temperature which is the basis for the fits.

For fitting the Arrhenius equation, we use linear regression on the logarithm of the duration Δ*t* and 1*/T* where *T* is the absolute temperature.

We also fitted a *Double Exponential* (DE) function, which contains two exponential functions and four free parameters:

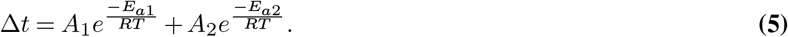

To fit the four parameters, we use a two-step approach. First, on a manually selected Arrhenius interval, we fit the standard Arrhenius law. This yields fitted values of the duration 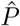. Next, we take the durations for the other temperatures and fit an Arrhenius law on the differences 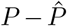, such that the sum of these fits describes the whole curve. Next, we used the resulting parameters as starting values in a full nonlinear fit of Eq. (5) using the `curve_fit` function from `scipy`.

The *Quadratic Exponential* (QE) function, which contains three free parameters, is given by

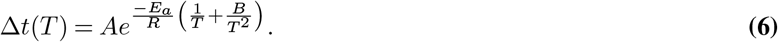

It can be fit using standard least squares on the logarithm of the duration as function of 1*/T*.

The *Power law – Exponential* (PE) function is given by

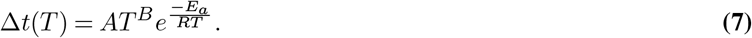

The fit is done using standard least squares. The code for fitting the functions is all included in the Github repository.

Note that for the fitting of the data for *X. tropicalis* embryos, we left out the point at the lowest temperature since this seems to be an outlier.

The bootstrap procedures we used to obtain distributions for the fitted activation energies is described in detail in Supplementary Note 2.

### ODE modeling

The equations and parameter values for both the 2 ODE model and the 5 ODE model are described in Supplementary Note 3. Simulations were performed in Python using `solve_ivp` from the `scipy` package. In general we simulated for a time of 1000 minutes using the `BDF` solver.

To detect the cycle period from a simulation, we detect peaks in the timeseries of the cyclin variable, and use the last two peaks to determine the period. If these peaks are too different in their *y*-values, we don’t consider the system oscillating as this would correspond to a damped rather than a sustained oscillation.

## Data and code availability

Source data and codes are provided with this paper.

- The western blots and droplet tracking data have been deposited in a Zenodo repository (doi: 10.5281/zenodo.15591678), and are publicly available as of the date of publication.
- All datasets necessary to reproduce the figures in the manuscript, as well as the code to produce the figures, can be found at the Gelens Lab Gitlab [https://gitlab.kuleuven.be/gelenslab/publications/temperature_scaling], and is publicly available as of the date of publication.
- All original modeling code has been deposited at the Gelens Lab Gitlab [https://gitlab.kuleuven.be/gelenslab/publications/temperature_scaling], and is publicly available as of the date of publication.
- All codes for image processing and analysis methods can be found in https://github.com/YangLab-um/temperature and https://github.com/YangLab-um/dropletDataProcessing, publicly available as of the date of publication.
- Any additional information required to reanalyze the data reported in this paper is available from the lead contact upon request.

## Acknowledgements

The work is supported by grants from the National Institutes of Health (R01 GM046383 and P50 GM107615, J.E.F., R01GM144584, Q.Y.), the National Science Foundation (MCB#2218083, Q.Y.), Internal funds KU Leuven (C14/23/130, L.G.), a junior research grant from the Research Foundation – Flanders (G074321N, L.G.), a doctoral fellowship from the Research Foundation – Flanders (11D0918N, J.R.) and postdoctoral fellowships from EMBL/EIPOD4 (Marie Skłodowska-Curie Cofund actions 847543, J.R.) and FNRS (Chargé de recherche, 40024839, J.R.). We acknowledge the support of the EMBL HPC resources. We thank Ernesto Flores for his contributions to the design and testing of the temperature chamber during his NSF REU project in the Yang lab in the summer of 2022.

## Author contributions statement

L.G., Q.Y. and J.E.F. conceived the study; F.T., A.V., C.P. and L.G. conducted the experiments; J.R., F.T. and L.G. analyzed the data; J.R. developed and analyzed the models; J.R., F.T., J.E.F. and L.G. prepared the figures; L.G. and J.E.F. wrote the manuscript, with L.G. incorporating feedback from all authors.

## Supplementary Note 1: Determination of local *E*_*a*_ **and** *Q*_10_

In the case of a duration or rate that scales according to the Arrhenius equation

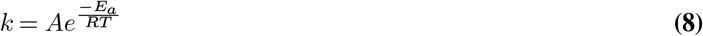

the activation energy is a constant. The formula above is equivalent to

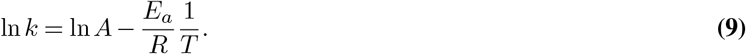

The value of *E*_*a*_ can thus be calculated from the slope of the line obtained when plotting ln *k* vs 1*/T*. Or,

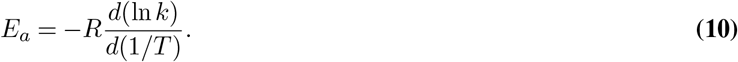

This equation can also be used as the definition of a local activation energy *E*_*a*_(*T*) for any function *k*(*T*).

Similarly, we can define a local *Q*_10_ value. In this section we explain how a *Q*_10_ value can be obtained for a process with temperature-dependent rate. The *Q*_10_ is the fold change in rate when the temperature increases by 10 degrees Celsius. If the value of *Q*_10_ is constant over all temperatures, the rate dependence should have the form

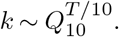

This form is different from the Arrhenius equation. It would therefore be incorrect to say of a process that its activation energy *E*_*a*_ *and* its *Q*_10_ are both constant over a range of temperatures.

The *Q*_10_ can be calculated as

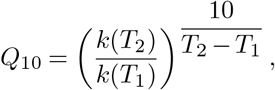

where *k*(*T*_*i*_) is the rate of the process calculated at temperature *T*_*i*_. Since this formula holds for any choice of *T*_1_ and *T*_2_, we can look at the limit *T*_2_ *→ T*_1_ and use this formula to define a *local Q*_10_, *Q*_10_(*T*). The use of ‘local’ for a number that is meant to convey what happens over a temperature change of 10 degrees is a bit contradictory, but we will make abstraction of this and use *Q*_10_(*T*) to indicate a local sensitivity to temperature.

If the rate of a process depends on temperature through any (differentiable) function *k*(*T*), then for any *h* we would have

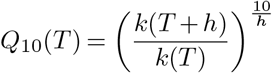

A Taylor expansion for small *h* gives that this is approximately equal to

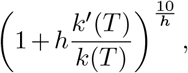

where 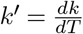. Using the definition of the exponential function, this goes to

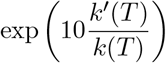

for small *h*. We thus define the local *Q*_10_ as

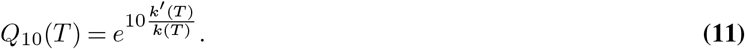

We can also express the formula for the local *E*_*a*_ (Eq. (10)) using the derivative of the rate:

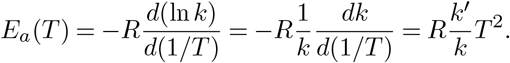

This also gives a link between local *Q*_10_ and *E*_*a*_:

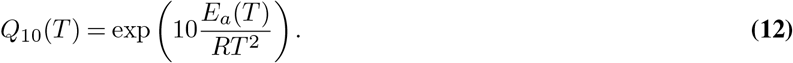

## Supplementary Note 2: Bootstrapping

The histograms that describe the uncertainty on the fitted activation energies on the rates, shown in Figs. 1 and 6, were obtained using a bootstrapping procedure. First, we generated a new dataset by resampling the rate measurements *with replacement* from the original data. In this case, we used a stratified bootstrap: we resampled per temperature. So if in the original dataset, there were three values of *E*_*a*_ for a given temperature, in a bootstrapped dataset there will also be three values for that temperature, and they are resampled with replacement from the original three. For each bootstrapped dataset, we computed the activation energy by fitting a straight line in the Arrhenius plot. We fitted on the median per temperature. We did this for 1000 bootstrapped datasets, and these 1000 values of *E*_*a*_ are represented in the histograms. The Github repository contains the code to reproduce the bootstrapping procedure.

## Supplementary Note 3: Computational cell cycle models

### 3.A Two-ODE cell cycle model

As described in the main text, we made use of a two-ODE cell cycle model based on one originally described in (66):

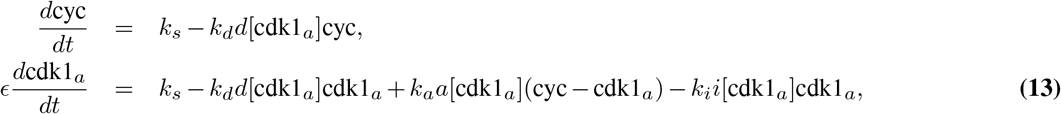

The first equation describes how cyclin B (cyc) is synthesized throughout the cell cycle at a rate *k*_*s*_ (nM/min) and how it is degraded at a rate *k*_*d*_ (1/min) by the proteasome after ubiquitination by active APC/C. APC/C is assumed to be activated instantaneously by active cyclin B – Cdk1 complexes (abbreviated as cdk1_*a*_) in an ultrasensitive way (given by *d*[cdk1_*a*_]). The second ODE describes the time evolution of active cyclin B – Cdk1 complexes, assuming that all synthesized cyclin B quickly binds to Cdk1 to form a complex. Moreover, the positive feedback of Cdk1 via Cdc25, and the double negative feedback of Cdk1 via Wee1 are included as an activating (*a*[cdk1_*a*_]) and inhibiting (*i*[cdk1_*a*_]) ultrasensitive function as well, motivated by direct experimental measurements of those response functions (67, 68). The different ultrasensitive functions have the following form:

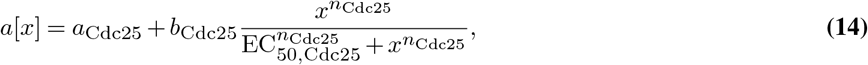

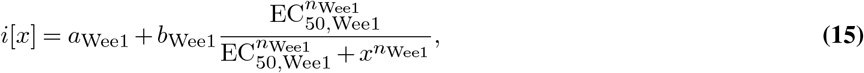

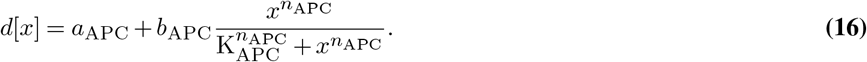

This type of model has previously been used to successfully describe various aspects of cell cycle oscillations (34, 66, 73). In the present work, we simplified this model further by omitting the degradation term from the Cdk1 equation, to have a clearer separation of the first ODE with cyclin synthesis and degradation, and the second ODE just describing Cdk1 activation and inactivation processes. This simplification does not significantly influence the results or our conclusions.

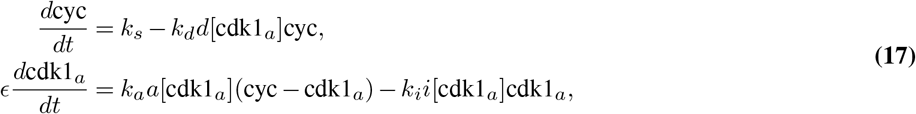

Using experimentally motivated parameters (66), model (17) reproduces cell cycle oscillations with a period of approx. 30 min (Fig. 3C). These oscillations manifest as a closed trajectory, a limit cycle, in the (cyc, cdk1_*a*_) phase plane (Fig. 3B, red). The phase-plane picture helps to better understand the existence of the oscillations via the intersection of nullclines (NCs). NCs are defined by points where 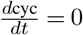 (Cyc NC) or 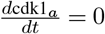 (Cdk1 NC). When *ϵ ≪* 1, oscillations occur at the intersection of the cyclin NC and the middle branch of the S-shaped Cdk1 NC (as depicted in Fig. 3B).

### 3.B. Five-ODE mass action model

#### 3.B.1. The model

We used a five-equation model for relaxation oscillations arising out of the interaction between Cdk1, Greatwall and PP2A (Fig. S6A). This pathway is a different part of the mitotic control system, which underlies the second mitotic switch. In the two-ODE model used in the main text, Cdk1 is involved in two feedback loops, through Wee1 and Cdc25. These feedbacks lead to bistability of Cdk1 activity as function of cyclin B levels. In the five-equation model, these feedback loops are not present, as is appropriate for cycles 2–12 in embryos (73). Here, any cyclin B-Cdk1 complex is directly activated assuming Cdk1 is present in excess over cyclin B: the response curve of Cdk1 activity as function of cyclin B levels would be linear. The production rate of cyclin B therefore directly corresponds to the production rate of active cyclin B-Cdk1 complexes. The equations we use are based on the paper by Hopkins et al. (72), who show that the equations for Greatwall, ENSA and PP2A lead to bistability. We complement their system with equations for Cdk1 and APC/C to turn the bistable system into a relaxation oscillator (77) (Fig. S6B).

We use the following equations:

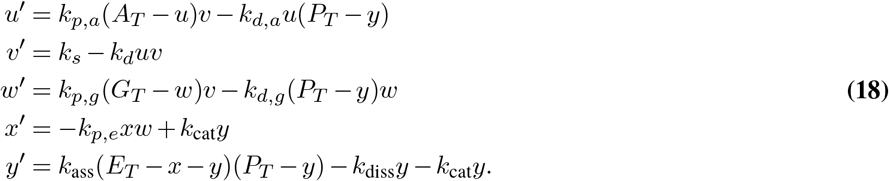

Here, *u* denotes active APC/C, *v* are the active cyclin B-Cdk1 complexes. The variable *w* corresponds to phosphorylated Greatwall, *x* is free, unphosphorylated ENSA and *y* is the complex ENSA-PP2A. The parameters of the model are the biochemical rates and the total amounts of APC/C (*A*_*T*_), Greatwall (*G*_*T*_), ENSA (*E*_*T*_), and PP2A (*P*_*T*_).

The first equation describes the activity of APC/C. We assume that APC/C can be converted from its inactive (unphosphorylated) form to its active (phosphorylated) form by Cdk1 through mass-action kinetics. The dephosphorylation is performed by PP2A. Note that *y* is the concentration of the ENSA-PP2A complex and *P*_*T*_ is total PP2A, such that the available PP2A is *P*_*T*_ − *y*. The second equation describes cyclin B-Cdk1 levels. These are governed by production and degradation of cyclin B, the latter of which is modeled through mass action. The equation for active Greatwall (variable *w*) describes the conversion between active and inactive Greatwall by Cdk1 and PP2A respectively — analogous to the APC/C equation. The fourth equation describes free unphosphorylated ENSA. This concentration decreases through phosphorylation of ENSA by Greatwall, and it increases through the dephosphorylation, which is mediated by PP2A. The final equation describes the concentration of the ENSA-PP2A complex.

The parameters, their meaning and their standard values can be found in Table 2. The parameter set is not based on experimental values, but was chosen to obtain a relaxation oscillation of amplitude (in the Cdk1 variable) and period that correspond to observations. All variables except for cyclin B-Cdk (*v*) are in arbitrary units, this is why the units of the rate constants look a bit awkward. As for the 2-ODE model, *k*_*s*_ and *k*_*d*_ are multiplied by 1.3 if simulations are to be compared with data from *X. tropicalis*.

#### 3.B.2. Interpretation in the phase plane

The model with only Greatwall, ENSA and PP2A (*w, x, y*) has been shown to produce a bistable response as function of the amount of active Cdk1 (*v*) (72). In our version, we added production of cyclin B and its degradation through ubiquitination by APC/C to turn the bistable system into a relaxation oscillator. Even though the system is five-dimensional, we can understand it in the phase plane. To do this, we perform a reduction to a two-variable system. We assume a quasi-steady-state condition on *w, x* and *u*, and set their derivatives to zero. When we do this, we assume that these variables evolve on a faster timescale than the others. We do expect that the levels of cyclin B-Cdk1 evolve on a slower timescale than the other variables: production and degradation are slower than the phosphorylation and dephosphorylation reactions. The reasons for keeping *y* as the additional variable and not one of the others is more practical: taking *w*^*′*^ = *x*^*′*^ = *u*^*′*^ = 0 leads to explicit expressions for these variables as function of *v* and *y*, which does not work if we take, say, *u* as remaining variable. We find

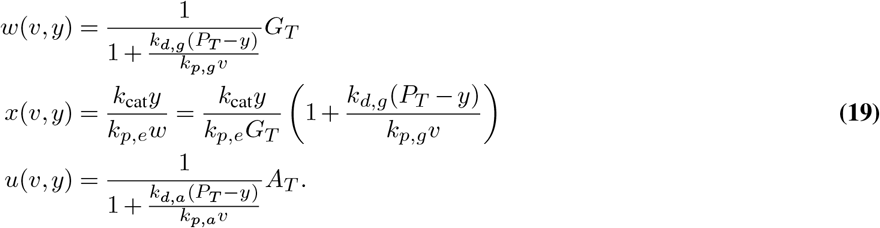

Using these, we can reduce the system to two equations:

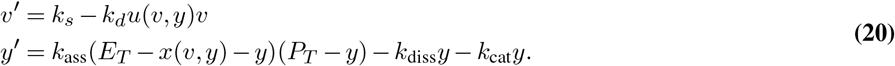

Fig. S6C shows the phase plane of this system with the associated limit cycle. The projection of the solution of the five-ODE system for the same parameter values is also shown. The two limit cycles are close in the phase plane, but their period is significantly different (Fig. S6D). This reduction shows that we can qualitatively understand the oscillations of this system in the phase plane. In particular, we confirm that the limit cycle is of relaxation type and goes around the underlying bistable switch. The cycle is, as in the 2-ODE model, driven by cyclin B accumulation and degradation. Once cyclin B-Cdk1 levels cross a threshold, the activity of the phosphatase is quickly suppressed. This allows activation of APC/C, which leads to cyclin degradation and brings the system back to a state of low Cdk1 activity. In particular, we can see from Eq. (20) that the production and degradation rates affect the non-S-shaped nullcline only.

For this model, we studied how the period of the oscillation changes if each of the parameters has an Arrhenius-like dependence on temperature. We analyzed how different activation energies for the different parameters can lead to different scaling as well as thermal ranges. As for the Yang-Ferrell model, this analysis can be interpreted in the phase plane, by examinining how parameter changes affect the location of the nullclines. From the phase-plane picture we can see that, when the oscillations disappear, the system becomes stuck in a state with either high or low phosphatase activity. As before, which one it is will depend on the relative magnitude of *E*_*a*_(*k*_*s*_) and *E*_*a*_(*k*_*d*_).

The steady state of the system only depends on the ratios of the following parameters:

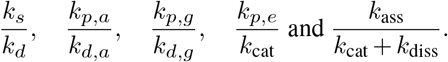

It follows that, for any set of activation energies such that

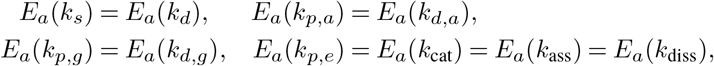

the steady state of the system is independent of temperature. Under the assumption that the relative magnitude of the timescales stays the same, this means that we would expect oscillations over a large range of temperatures if these rates scale in a similar way. These ratios usually have the rates for two counteracting reactions in numerator and denominator.

### 3.C. Details on the parameter sweep for fitting the 2-ODE model to embryo data

In Fig. 3E, we show fits of the 2-ODE model to the duration of the embryonic cycle in different cases. These were obtained by performing computational parameter sweeps. All nonscaling parameters were as in Table 1, and for *X. tropicalis* we multiplied *k*_*s*_ and *k*_*d*_ by 1.3. We kept *k*_*i*_ and *k*_*a*_ and only scale *k*_*s*_ and *k*_*d*_.

**Table 1:**
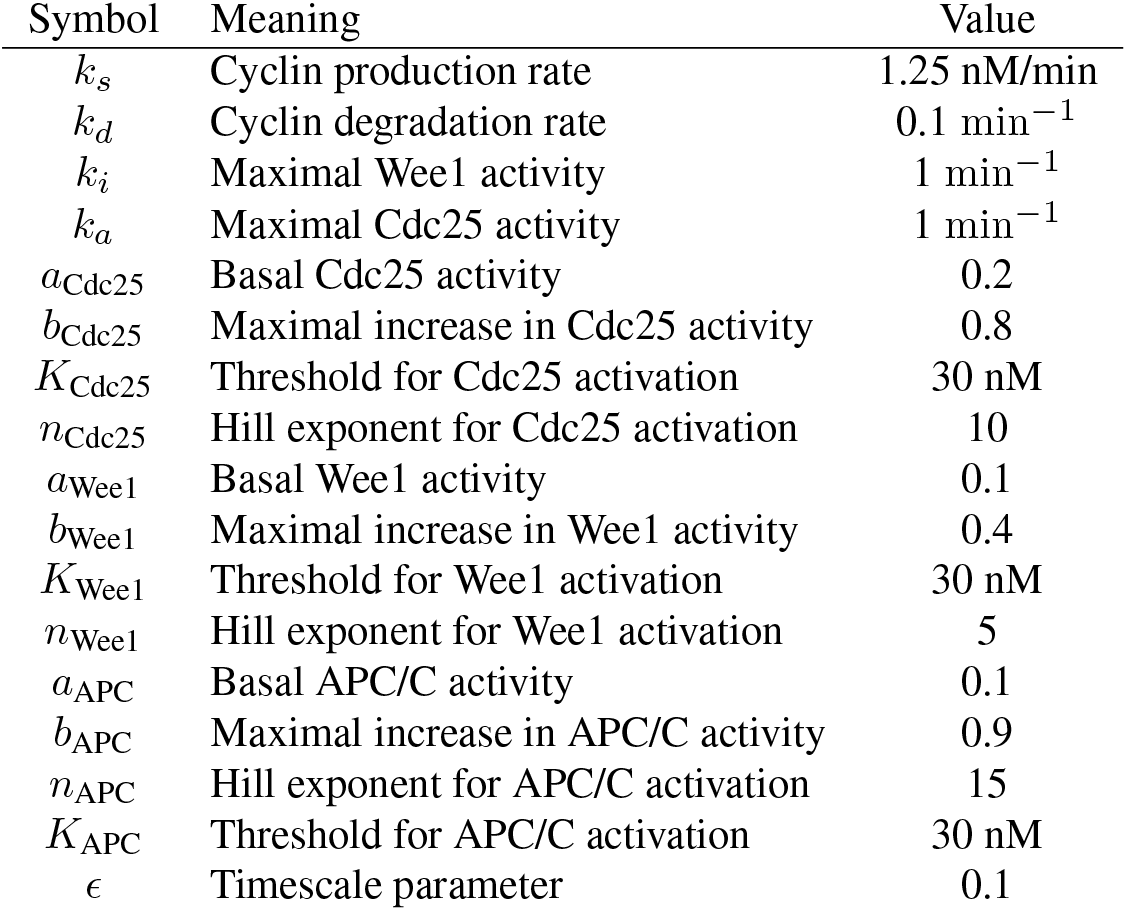
Parameter values used in the 2-ODE cell cycle model. These are the basal values that correspond roughly to the period of the *X. laevis* and *D. rerio* cell cycle. To describe the *X. tropicalis cycle*, which is faster, we multiply *k*_*s*_ and *k*_*d*_ by 1.3. The basal values of *k*_*s*_, *k*_*d*_ and *ϵ* are also different for extracts: these values are obtained as part of the fitting to the time series.

**Table 2:**
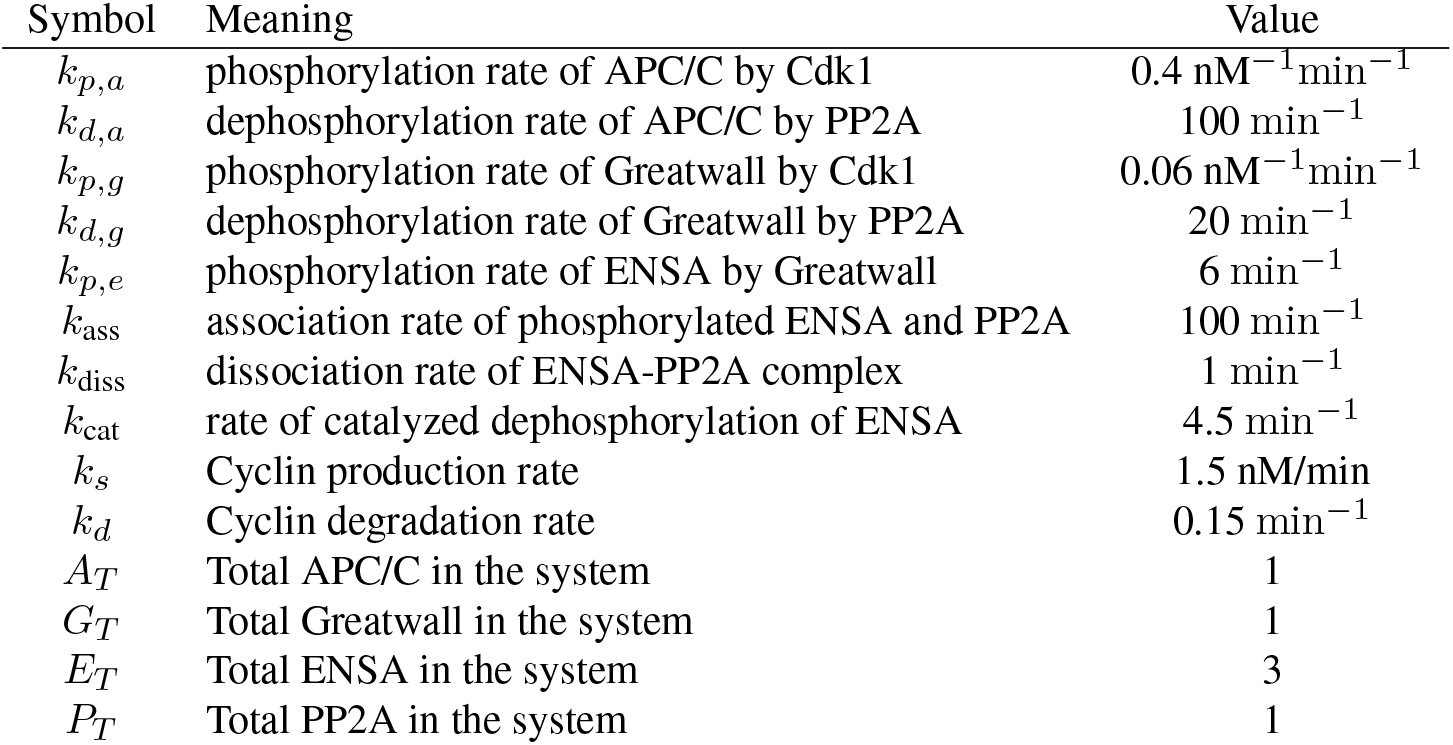
Parameter values used in the mass action model.

In particular, we performed two sweeps:

1. Both *k*_*s*_ and *k*_*d*_ scale Arrhenius, but their activation energies differ. We varied *E*_*a*_(*k*_*s*_) and *E*_*a*_(*k*_*d*_) from 0 to 150 kJ/mol in steps of 5 kJ/mol. For each combination of *E*_*a*_, we perform simulations for temperatures from 0 to 50°C and save the period to file. Next, we calculate the distance between simulated periods and the duration from the data as follows. First, if the simulation did not yield oscillations for the full range of temperatures in the dataset, we consider the error infinite. In the other case, we calculate the Mean Squared error on the logarithms of the durations. The curves shown in Fig. 3E, Case 1 and 3, are obtained by finding the *E*_*a*_(*k*_*s*_) and *E*_*a*_(*k*_*d*_) that minimize the MSE under the constraint that *E*_*a*_(*k*_*s*_) = *E*_*a*_(*k*_*d*_) (Case 1, green line), *E*_*a*_(*k*_*s*_) *> E*_*a*_(*k*_*d*_) (Case 3, orange line) and *E*_*a*_(*k*_*s*_) *< E*_*a*_(*k*_*d*_) (Case 3, red line).
2. For the biphasic response, we assumed that the temperature scaling of *k*_*s*_ is given by a double-exponential curve

In this parameter sweep, *k*_*d*_ scales Arrhenius and has the same activation energy as *k*_*s*_ for the lower-temperature regime. The difference now is the second exponential term which leads to a bending of the curve for *k*_*s*_ at high *T*.

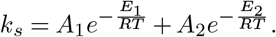

We performed a sweep over *E*_1_ and *E*_2_, with *E*_1_ ranging between 0 and 150kJ/mol and *E*_2_ between –150 and 0 kJ/mol. We determined the prefactors by fixing the rate of *k*_*s*_ at a reference temperature at 18°C, and by fixing the optimal temperature *T*_*m*_. We also scanned over different values of *T*_*m*_ around the optimal temperature from the data, in particular we simulated for *T*_*m*_ for integer values of the temperature between *T*_*m*,data_ − 7 and *T*_*m*,data_ + 7, where *T*_*m*,data_ is 28°C for *X. laevis*, 30°C for *X. tropicalis* and 32°C for *D. rerio*.

Distance between model simulaton and data was determined as above.

### 3.D. Fitting temperature-dependent computational models to data using the ABC algorithm

#### 3.D.1. Fitting cycling extract data with the two-ODE model

In Fig. 5 we show the results of fitting the parameters to the time scaling of extract data. These fits were obtained using Approximate Bayesian Computation – Sequential Monte Carlo (ABC-SMC) (74). This algorithm sequentially samples parameter sets that provide better and better fits to the data. The output of the algorithm is *N* parameter sets Θ_*i*_ with associated weights *w*_*i*_. Each of these parameter sets provides a fit closer than a prescribed distance *ε* to the data. The *N* weighted parameter sets constitute a sample from the posterior distribution *P* (Θ | *d*(*x*^*\**^, *x*_0_) *< ε*), where *x*_0_ is the data, *x*^*\**^ is the data resulting from a simulation with parameters Θ and *d* is a distance function. If *ε* is small, this distribution approximates the posterior distribution *P* (Θ | *x*_0_): the probability that a parameter set Θ is the true one, given the observed data. In ABC-SMC, the value of *ε* is lowered over the course of different generations as a way of getting better and better approximations of the posterior. We use the implementation of this algorithm given in pyABC (75).

For the extract fits, we describe the temperature scaling of each of the rates *k*_*s*_, *k*_*d*_ and *ϵ* using a double-exponential formula. For *k*_*s*_ and *k*_*d*_, we parametrize the rate as

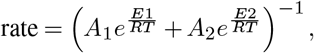

and for *ϵ*, which has units of duration and not rate, we use

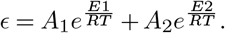

Instead of the four parameters *A*_1_, *E*_1_, *A*_2_, *B*_2_, we decide to use more interpretable parameters: the basal value of the parameter at 18 degrees Celsius, the temperature *T*_*m*_ at which is maximal (minimal for *ϵ*) value is obtained, and the two activation energies *E*_1_ and *E*_2_. Note that we can map *k*_0_, *T*_*m*_, *E*_1_, *E*_2_ to *A*_1_, *E*_1_, *A*_2_, *E*_2_ directly.

A parameter vector Θ contains 12 values [*E*_1_(*k*_*s*_), *E*_2_(*k*_*s*_), *T*_*m*_(*k*_*s*_), *k*_*s*,0_, *E*_1_(*k*_*d*_), *E*_2_(*k*_*d*_), *T*_*m*_(*k*_*d*_), *k*_*d*,0_, *E*_1_(*ϵ*), *E*_2_(*ϵ*), *T*_*m*_(*ϵ*), *ϵ*_0_]. Each parameter set thus defines three functions *k*_*s*_(*T*), *k*_*d*_(*T*) and *ϵ*(*T*). We simulate the 2-ODE model over the temperatures from the dataset, using the rates defined by these scaling functions. All the other model parameters are kept to their basal value (Table 1). For each temperature, we detect whether the system is oscillating using the peaks of the time series, as described in Methods. For oscillating systems, we then use the *cdk* timeseries to determine the rising (min to max) and falling (max to min) durations.

The output of the simulation is thus *X*_simulation_ = {(*R*_*i*_, *F*_*i*_), *i* = 1,… *N*}: the duration of rising and falling part of the cycle, for each temperature *T*_*i*_ in the dataset (*N* being the total number of temperature points). These data from the simulation are then compared to the same data obtained from the extract time series *X*_data_. The distance function we use is

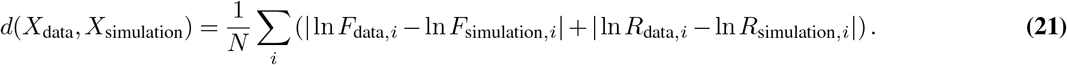

We thus consider the differences of the logarithms of the rising and falling times, for each temperature, and take the average of their absolute values. The durations can vary quite a bit in absolute value, and we use the logarithms to prevent the algorithm being skewed to approximating the large durations (at extreme temperatures). We take *d*(*X*_data_, *X*_simulation_) = *∞* if there is a temperature in the dataset for which the simulation did not produce an oscillation. The consequence is that we only search for parameter values for which the model produces oscillations over *at least* the range we see in the experiment.

For the extract simulations, we use 1000 particles per generation of the ABC algorithm. The prior distributions were uniform distributions for all parameters, for the *E*_1_ between 0 and 200 kJ/mol, for the *E*_2_ between –200 and 0 kJ/mol, for the *T*_*m*_ between 10 and 45 degrees Celsius, for *k*_*s*,0_ between 0 and 5, for *k*_*d*,0_ between 0 and 2 and for *ϵ*_0_ between 0 and 100.

We let the algorithm run for 40 generations and inspected visually the resulting fits described. We did not retain the last generation for the figures shown, because these fit the data too closely. Since the data itself has variability we did not want to overfit. We picked the 25th generation for the results in the main text. Fig. S10 shows the marginal parameter distributions in this generation. All the parameters for *k*_*s*_, as well as *k*_*d*,0_ and *ϵ*_0_, are clearly centered on one value. Moreover, 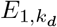 and 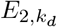 are peaked close to zero. For the other parameters, the distribution is much wider. This is also clear from the plots in Fig. S11: These show the marginal distribution of the different parameters over the different generations of the ABC SMC run. For the parameters peaked around one value, we see clear convergence of the posteriors. For the others, this is less clear.

#### 3.D.2 Fitting embryo data with the two-ODE model

Here we describe the setup of the ABC algorithm for fitting the embryo data using purely Arrhenius scaling on the rates (Fig. S5). We use the two-ODE cell cycle model to capture the period scaling observed in the data for *Xenopus laevis, Xenopus tropicalis* and *Danio rerio*.

For this model we use four different parameters: the activation energies of *k*_*s*_, *k*_*d*_ and of the ‘activation’ and ‘inactivation’ reactions, *k*_*a*_ and *k*_*i*_ in Eq. (4). The activation can be thought of as scaling the dephosphorylation rate of Cdk1 by Cdc25. The inactivation rate scales phosphorylation by Wee1. Each simulation produces the period *P*_*i*_ of the oscillation for each of the temperatures *T*_*i*_ in the embryo dataset (or zero if for a temperature there is no oscillation).

We compare the distance between the simulated dataset *X*_simulation_ = {*P*_*i*_, *i* = 1 … *N*} (the durations of the embryonic cycles for all temperatures *T*_*i*_) and the observed data *X*_data_with the distance function

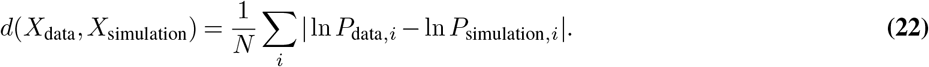

The distance is set to infinity if there is at least one temperature *T*_*i*_ for which the simulation does not produce an oscillation.

The basal rates at 18°C are as in Table 1, but for *X. tropicalis* we multiply *k*_*s*_ and *k*_*d*_ by 1.3 to account for its faster cycle. Since the temperature scaling of a rate is defined by

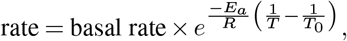

only varying the *E*_*a*_ will lead to always the same value of the rate at *T*_0_, 18 degrees Celsius in our case. This means that all period curves will go through the same point at 18 degrees. To circumvent this, we include an additional parameter which adds an overall scaling of the period, by scaling the basal values of *k*_*s*_ and *k*_*d*_. In all the samples, this parameter is very close to one.

We use 200 particles in the ABC run and let the algorithm run for 15 generations. The prior distributions for all activation energies were uniform on [0, 150] kJ/mol. Fig. S5 shows the resulting fits from the ABC algorithm.

The two-dimensional probability densities in Fig. S5B are projections of the points (*E*_*a*_(*k*_*s*_), *E*_*a*_(*k*_*d*_),*E*_*a*_(*k*_*a*_), *E*_*a*_(*k*_*i*_)) into two different planes. Lighter color means higher probability. The bottom row in this figure shows a white cloud along the diagonal. This means that the best fits to the data are obtained when *E*_*a*_(*k*_*a*_) *≈ E*_*a*_(*k*_*i*_). The upper heatmaps show that the activation energies of *k*_*s*_ and *k*_*d*_ need to be different (off-diagonal clouds) for good fits. Whereas for *X. tropicalis* and *D. rerio*, good fits are obtained when *E*_*a*_(*k*_*s*_) is larger than *E*_*a*_(*k*_*d*_), for *X. laevis* there are two off-diagonal clouds. These density plots were obtained from the 200 weighted points that are the output of the ABC algorithm, smoothed using a Gaussian kernel. In Fig. S5A we show the actual fits corresponding to each of these sampled points in gray, with more transparent lines corresponding to parameter sets with lower weight. In orange, we show the best fit (lowest distance). The fits are quite good overall, but the upper thermal limit and the bend upwards are not so easy to capture.

The results from the ABC algorithm give us similar insights to what we obtained from a full parameter scan (the results in Fig. 3E). However, it is less computationally intensive. We conclude from the results here that good fits to the data can be obtained using the two-ODE model using Arrhenius scaling for the four main parameters. The parameter sets that best capture the scaling in the data and the upper thermal limit have activation energies for Cdc25 and Wee1 (activation/inactivation) that are close and different values for the production and degradation activation energies.

#### 3.D.3. Fitting embryo data with the five-ODE mass action model

The five-ODE mass action model has ten different activation energies. A full parameter scan is unfeasible here, but the ABC algorithm can still be used. We also used 200 particles and let the algorithm run for 15 generations. Prior distributions were uniform on [0, 150] kJ/mol for all activation energies. We also included an overall scaling factor to avoid every temperature-period curve going through the same value at 18 degrees, as explained above. This value was close to one for all sampled parameter sets.

Good fits can generally be obtained using this model, although there are not so many parameter sets that can capture the bend upwards for high temperatures Fig. S6C. We are not entirely sure whether this is unexpected or not: in general more parameters to vary increases the possibility of an accurate fit, and ten parameters is already quite a lot. On the other hand, the oscillations in this system are quite sensitive to changes in the rate constants, making a good fit more difficult. We did observe that the sets of activation energies that provide a good fit are more localized in parameter space, whereas for the two-ODE model there were quite broad areas that gave a good fit.

The output of the algorithm is now a set of parameters which represents a probability distribution in ten-dimensional space. This is not so easy to visualize. We can however look at some summary statistics of this distribution, as in Fig. S6D. This figure shows the marginal distribution of the different activation energies and the pairwise correlations between them. Some of the pairs with high correlations correspond to antagonistic rates, such as *k*_*p,g*_ and *k*_*d,g*_. This suggests that the parameter sets that provide a good fit have more or less equal activation energies for the faster reactions, forward and backward, and that the thermal limits are generated by an imbalance between the *E*_*a*_ of production and degradation rates. This is a tentative conclusion, however, since the complete distribution of the distribution of all 10 rates is not completely captured by looking only at pairwise correlations and marginal distributions.

## Supplementary Note 4: Direct estimation of the cyclin production rate scaling from extract time series

In Fig. 5B, we plot estimates of the scaling of *k*_*s*_ with temperature obtained directly from the time series. These measurements are the slopes of the increasing part of the Cdk1 activity. We use a heuristic algorithm to determine these slopes automatically.

The time series mostly have a slow linear increase and then a fast jump. We want to capture the slope of the linear increase only. To detect the interval of this linear increase, we do the following. The idea behind this is shown in Fig. S12, and the code is included in the Github repository.

1. We start from one cycle of Cdk1 activity with values (*t*_*i*_, *u*_*i*_) with *i*_1_ *≤ i < i*_2_ (as in Supplementary Note 5).
2. Let *i*_*m*_ = argmin_*i*_ *u*_*i*_, the index of the minimal value of the *u*-values.
3. For *j* in *i*_*m*_ + 2 … *i*_2_, we determine the slope of a linear fit on the points (*t*_*i*_, *u*_*i*_), *i* = *i*_*m*_ … *j*. In other words, we start from the first three points starting at *i*_*m*_, and then always add one more point to the right until we hit the last point of the cycle. For each of these intervals, we obtain a slope *a*.
4. This yields a set of slopes *a*_*j*_. We then look at this array starting from its last element, i.e. we have *a*_0_ be the slope obtained fitting from *i*_*m*_ to the last point of the cycle, *a*_1_ from *i*_*m*_ to the one-to-last point, etc.
5. We determine *r*_*j*_ = (*a*_*j*_ −*a*_*j*−1_)*/a*_*j*_, the relative errors in this array of slope (*j* starting from 1).
6. Reasoning that first, *r*_*j*_ will be large because at the end of the cycle there are large variations, and that for points on the linearly increasing part of the cycle the relative errors do not change much, we fix the final slope as the one corresponding to the first relative minimum of the *r*_*j*_ (we exclude the first two *r*_*j*_ to avoid detecting the very first point as a minimum).

## Supplementary Note 5: Determination of the average cycle of Cdk1 activity

In Fig. 5, we show an average cycle shape obtained from extracts. Here we explain how we obtained this. This is also illustrated in Fig. S13, and the code is included in the Github repository. For each temperature, and for each droplet, we have a set of time series measurements (*t*_*i*_, *u*_*i*_), where *u*_*i*_ is the measurement of Cdk1 activity (FRET ratio). We then process these data as follows:

1. We determine all the indices *i* that belong to one cycle, which is defined as all the datapoints between two peaks: we select(*t*_*i*_, *u*_*i*_) for *i*_1_ *≤ i ≤ i*_2_. Here *i*_1_ and *i*_2_ are the indices corresponding to peaks in the Cdk1 signal.
2. We rescale time to the interval [0, 1]: set 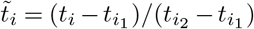 and we shift the values of *u* vertically by subtracting the mean: 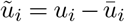.
3. We interpolate the values of 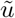 at 100 evenly spaced time points in the interval [0,1], obtaining a new time series 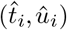 with *t*_1_ = 0 and *t*_100_ = 1.
4. We do this for all the droplets that have a given temperature. This yields (*t*_*k,i*_, *u*_*k,i*_) where *k* indexes the different time series (droplets). The average shape of the time series is then obtained by taking, for each *i*, the median of the values of *u*. The resulting time series is (*t*_*i*_, *U*_*i*_) with *U*_*i*_ = median_over *k*_ *u*_*k,i*_

## Supplementary Note 6: Fitting rate measurements of individual regulatory processes

We describe how the rates for cyclin B synthesis, Cdk1 activity, PP2A activity and APC/C activity were fitted from time series to obtain Fig. 6. For the linear fit we used the numpy function `polyfit`, and for the nonlinear fits we used the function `curve_fit` from the `scipy.optimize` package.

### 6.A Cyclin B synthesis rates

Fig. S14A-B shows a representative Western blot. By calculating the integrated density for each band and subtracting the background using FIJI, we obtain the intensity for each time point. This value is then divided by the average intensity for a CSF extract, to obtain the normalized intensities as shown in Fig. S14C. To obtain the cyclin accumulation rates, we next fit the slopes for each of the cycles, e.g. for the points indicated in S14C. The obtained results for the cyclin accumulation rates *k*_*s*_ from all Western blots, obtained from three independent extracts, are summarized in Fig. S8A (left).

### 6.B. Cdk1 rates

The time series from which we derive the rates are shown in Fig. S16. We fit a function of the form

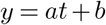

to the time series, and save *a* as the resulting rate. We do this for every replicate, giving us multiple rate measurements per temperature. We used the non-P32 decay adjusted rates.

### 6.C. PP2A rates

The time series from which we derive the rates are shown in Fig. S17. We fit a function of the form

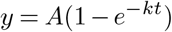

to the time series. The rate is determined as the derivative of this function at *t* = 0, i.e. *kA*.

### 6.D. APC rates

The time series from which we derive the rates are shown in Fig. S15. We fit a function of the form

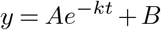

to the time series where we impose that *A, B, k >* 0. The rate we save is *k*. For the fit, we only use datapoints for times larger than or equal to 10 minutes, because the exponential decay does not start in the beginning.

## Main figures

## Supplemental Figures

**Fig. S1:**
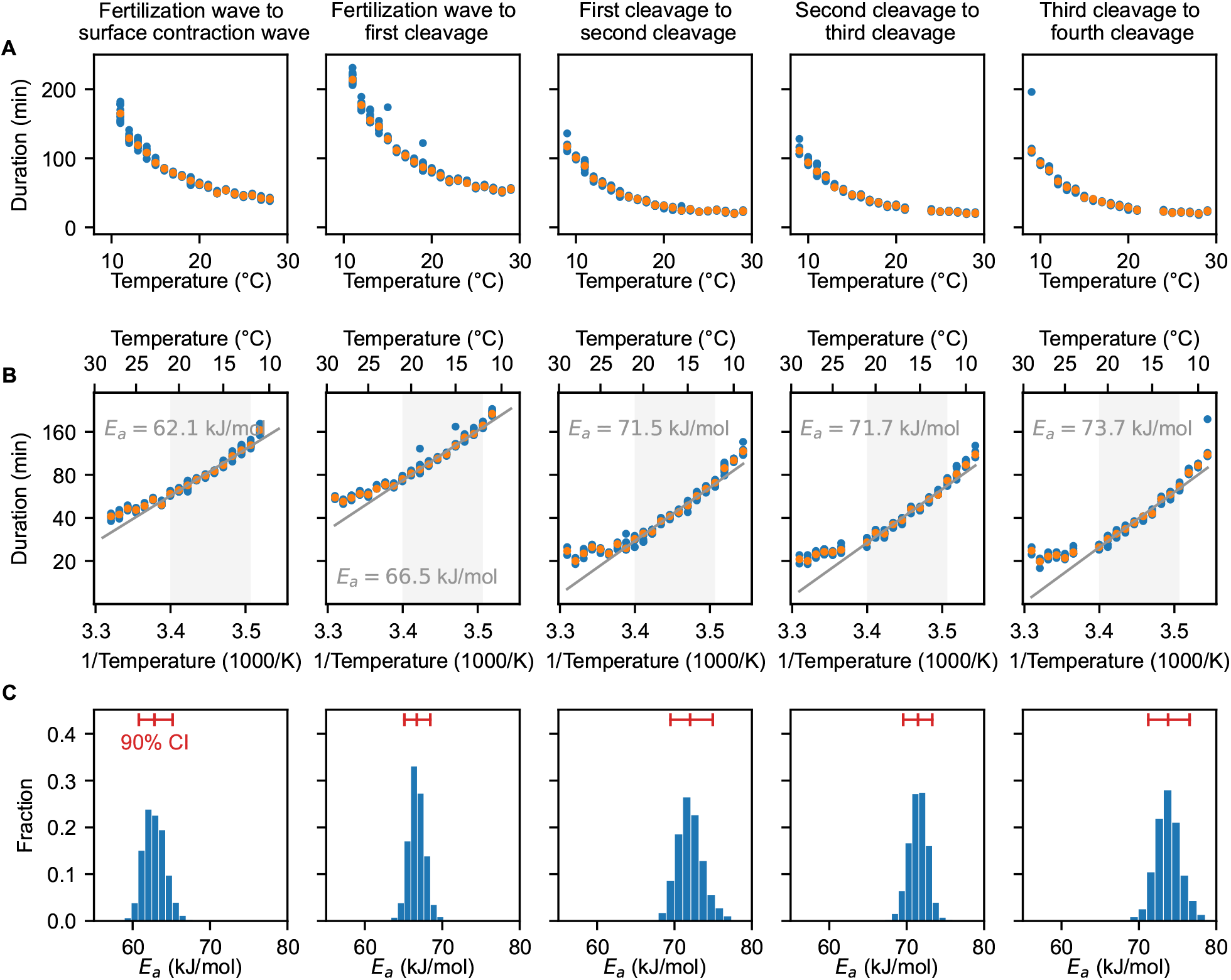
Cell division timing in early Xenopus laevis embryos scales approximately Arrhenius over a wide range of temperatures. A. Duration of several early developmental periods in function of temperature in the range [*T*_min_ = 9°C, *T*_max_ = 29°C]. B. An Arrhenius fit is shown for the values between 12°C and 21°C, with the apparent activation energy indicated. C. Bootstrapping provides a probability distribution for the apparent activation energies. The mean and 90% confidence interval (CI) are also indicated.

**Fig. S2:**
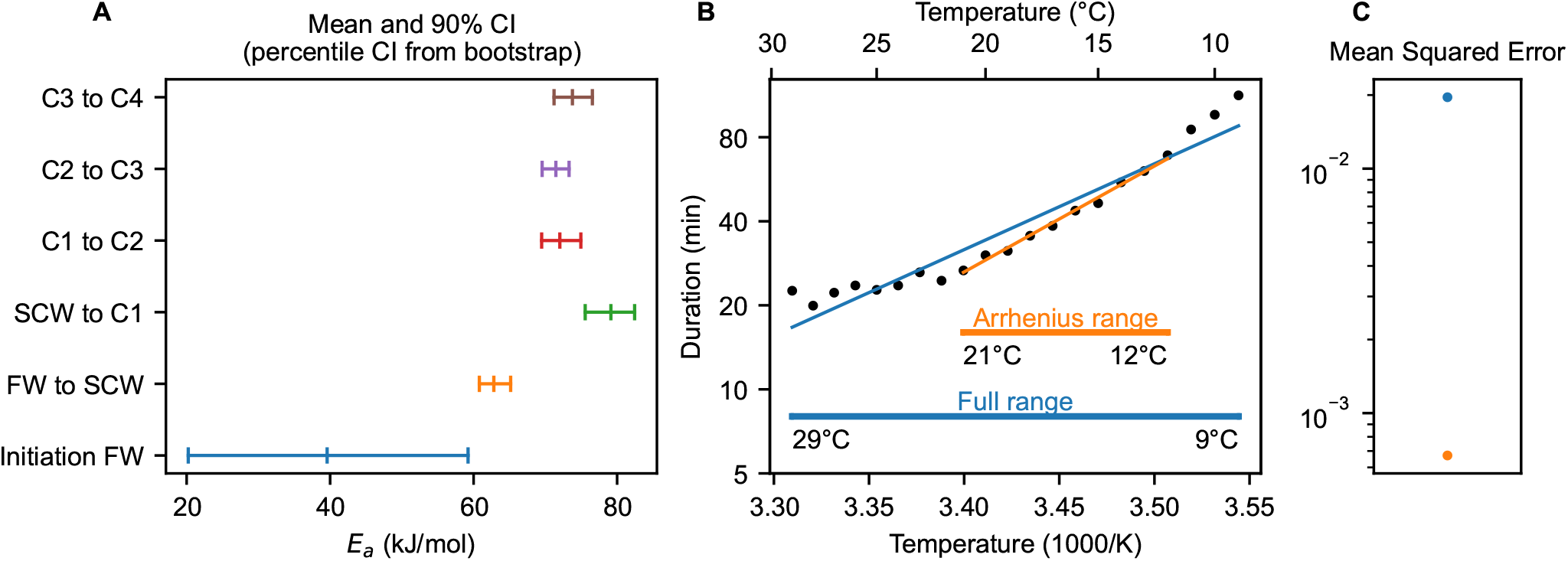
Cell division timing in early Xenopus laevis embryos does not scale Arrhenius over the whole temperature range. A. In Fig. 1C, we show the duration of several early developmental periods in function of temperature in the range [*T*_min_ = 9°C,*T*_max_ = 29°C], with the apparent activation energy as obtained by an Arrhenius fit between 12°C and 21°C in Fig. 1D. Bootstrapping provides a probability distribution for the apparent activation energies (Fig. 1E). Here, we show the mean and 90% confidence interval (CI) for comparison. FW is fertilization wave, SCW is surface contraction wave, C means cleavage. B. Cleavage cycle duration in function of temperature for the second to fourth cell cycle in the range [*T*_min_ = 9°C,*T*_max_ = 29°C] for *Xenopus laevis*. Optimal fits using single exponential Arrhenius (SE) are shown in two different temperature ranges: from 12°C and 21°C (orange), and the whole temperature range (blue). The mean square error (MSE) is much higher over the whole temperature range than within the selected range (panel C), indicating that the Arrhenius equation does not fit the data well over the whole measured data range.

**Fig. S3:**
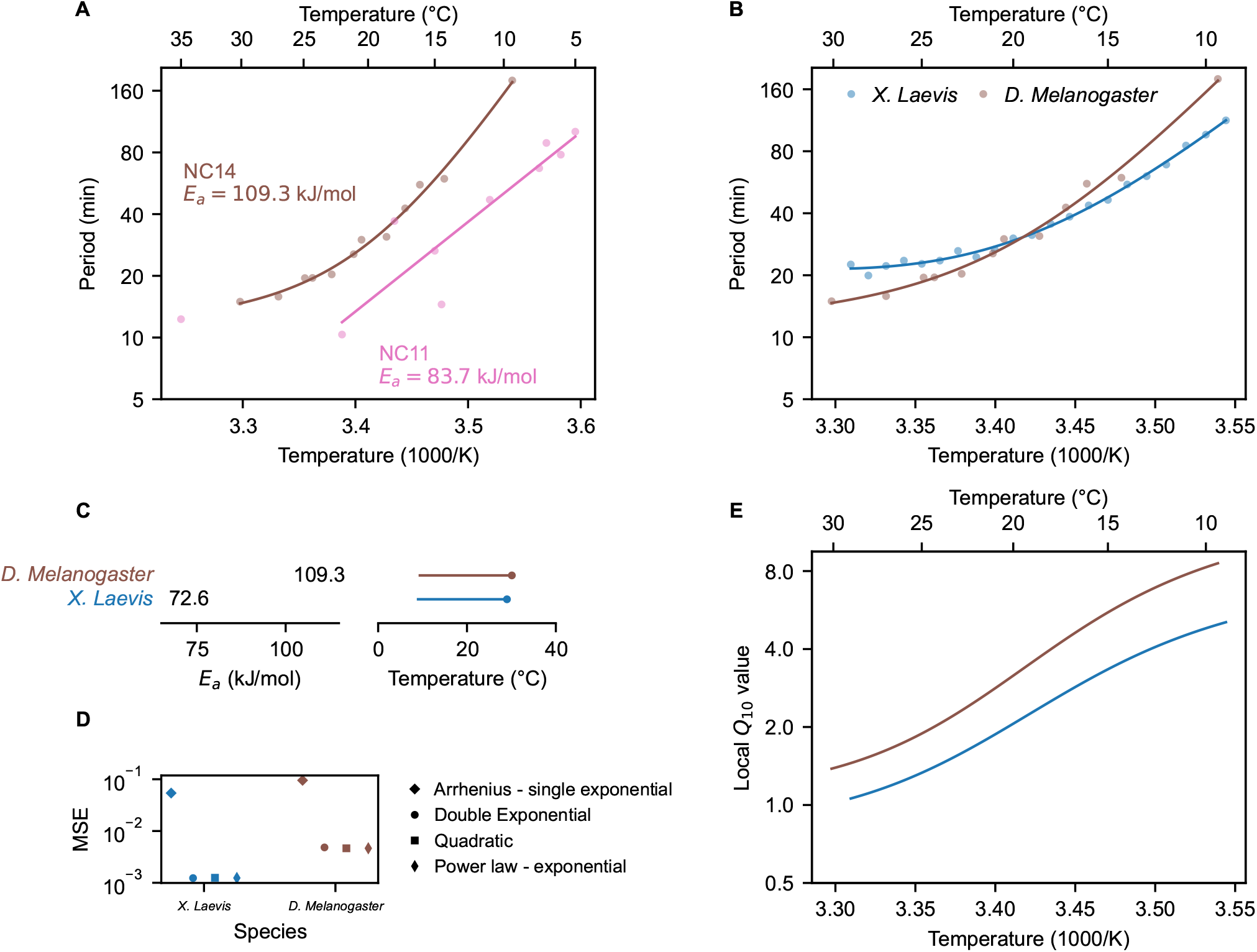
Temperature scaling of embryonic processes in *Drosophila melanogaster*. A. Median cleavage period in function of temperature for the eleventh and thirteenth cell cycle *D. melanogaster*. Data for NC11 from Falahati et al. (33), for NC14 from Crapse et al. (32). Optimal fits using a double exponential (DE) function are overlayed. B. Median cleavage period in function of temperature for the second to fourth cell cycle (all pooled) in *X. laevis* (this work), and for the eleventh and fourteenth cell cycle *D. melanogaster* (32), together with double exponential (DE) best fits. C. Activation energy, minimal and maximal temperature for *X. laevis* and *D. melanogaster*, corresponding to the curves in panel B. The dot shows the optimal temperature, which in this case is also the maximal temperature. D. Comparison of the mean squared error (MSE) for fits with different functional forms, as in Fig. 2G. E. Using the best DE fit, the local *Q*_10_ value is plotted as function of temperature.

**Fig. S4:**
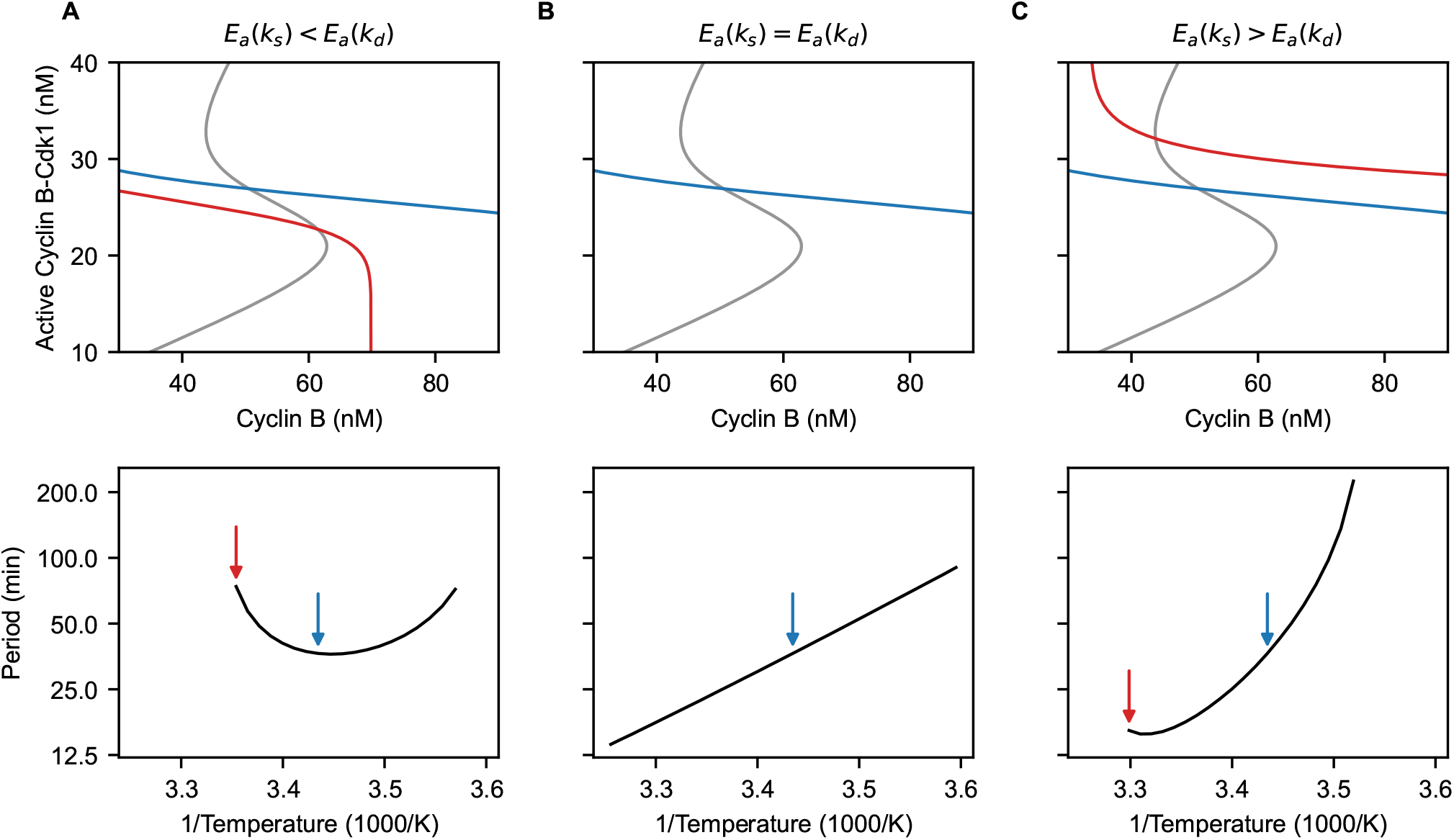
Temperature dependence of nullclines in the phase plane. In the two-ODE cell cycle model, thermal limits are determined by intersection of nullclines. Depending on the relative size of *E*_*a*_(*k*_*s*_) and *E*_*a*_(*k*_*d*_), the non-S-shaped nullcline (red/blue) shifts upward or downward with rising temperatures. When the intersection of the nullclines lies on the upper or lower branch of the S-shaped nullcline (gray), oscillations cease to exist. A. The system ends up in a low-activity state at high temperatures. B. The second nullcline is temperature independent. There is no thermal limit. C. The system is stuck in a high-activity state at high temperatures.

**Fig. S5:**
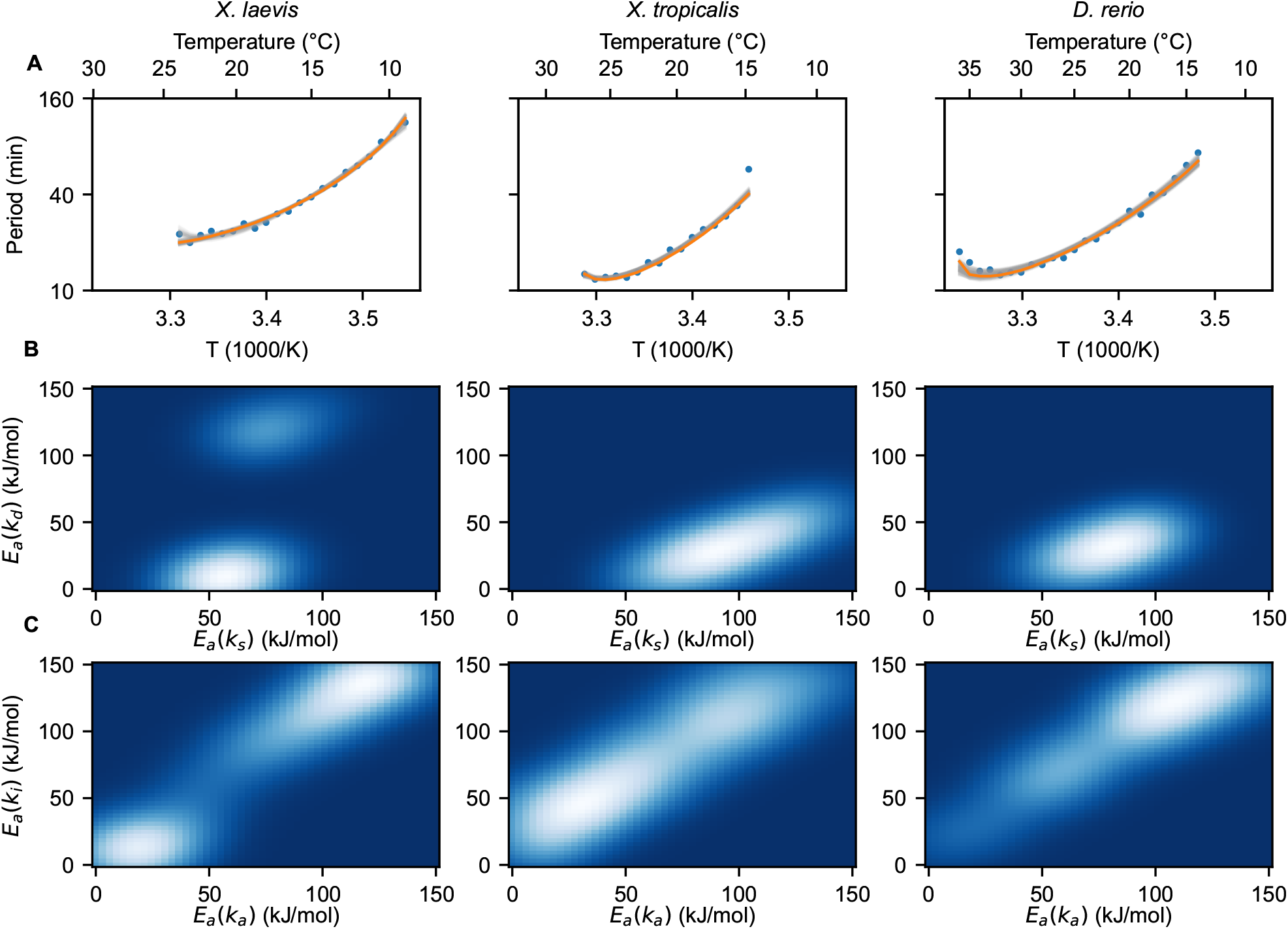
Optimal fits of the two-ODE model to the measured data using the ABC method. A. Resulting fits from the ABC algorithm for the two-ODE model. Gray lines show the 200 resulting parameter sets, with more transparent lines corresponding to points with lower weight. The orange line is the best fit (smallest distance). B. Projection of the four-dimensional probability density onto the (*E*_*a*_(*k*_*s*_), *E*_*a*_(*k*_*d*_))-plane. C. Projection onto the (*E*_*a*_(*k*_*a*_), *E*_*a*_(*k*_*i*_))-plane. The heatmaps were constructed from 200 weighted samples and smoothed with a Gaussian kernel.

**Fig. S6:**
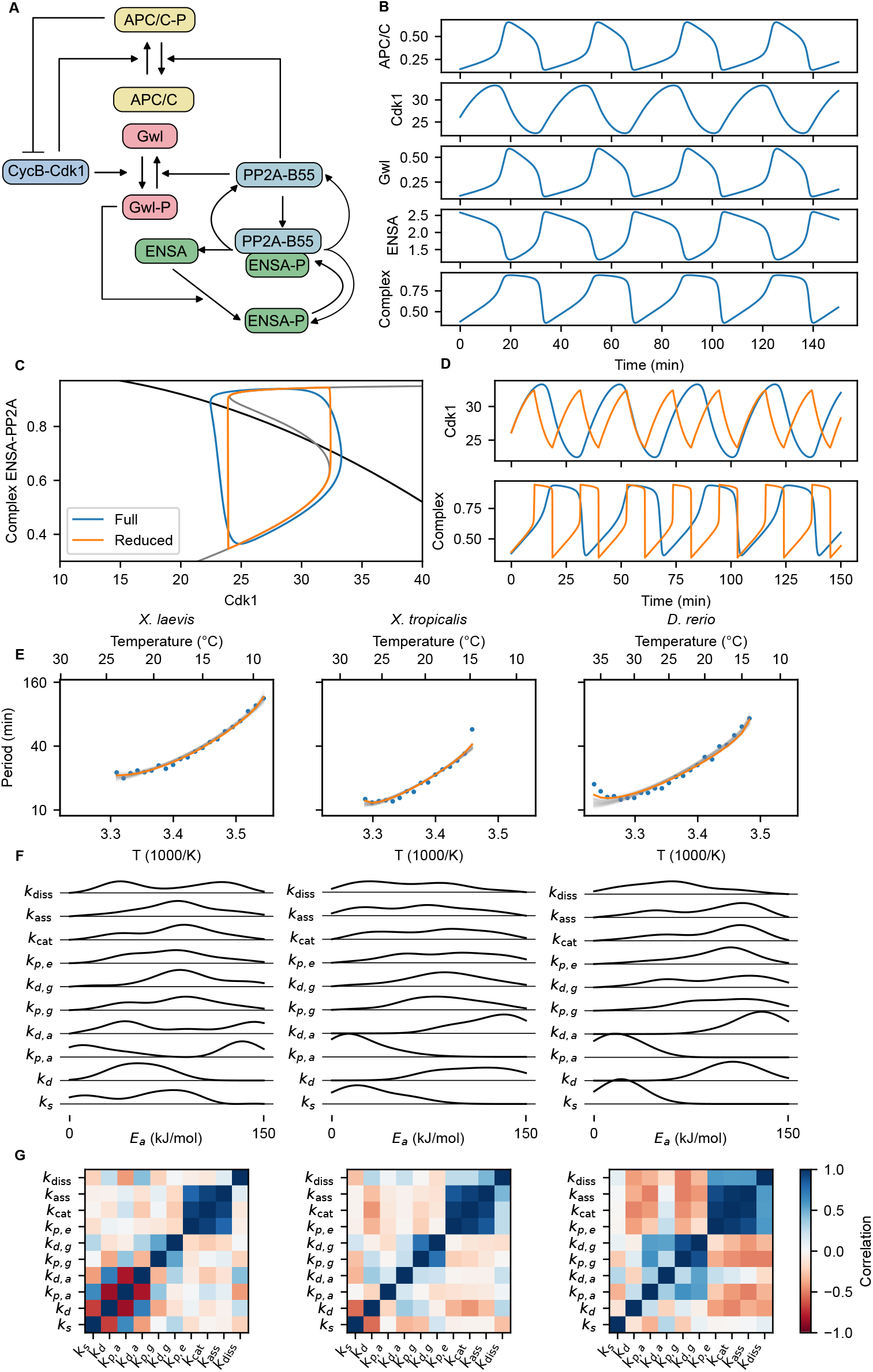
The five-ODE mass action model and fits to the measured data using ABC method. A. The interaction diagram for the five-ODE mass action model. B. Representative time series of a simulation of the 5-equation model. C. Phase plane of the reduced two-ODE model and projection of the five-ODE model onto this plane. Nullclines are shown in black and gray, the blue limit cycle is the projection of the oscillation of the five-ODE system and the orange limit cycle is the one in the two-ODE system. D. Time series of corresponding variables in the full (blue) and reduced (orange) model. E. Resulting fits from the ABC algorithm for the mass action model. Gray lines show the 200 resulting parameter sets, with more transparent lines corresponding to lower-weighted points. The orange line is the best fit (smallest distance). F. Results of the ABC algorithm for the mass-action model. Marginal distributions of the different activation energies (smooth distribution obtained using Gaussian kernel density). G. Pairwise correlation of the activation energies of the different rates computed from the result of the ABC algorithm.

**Fig. S7:**
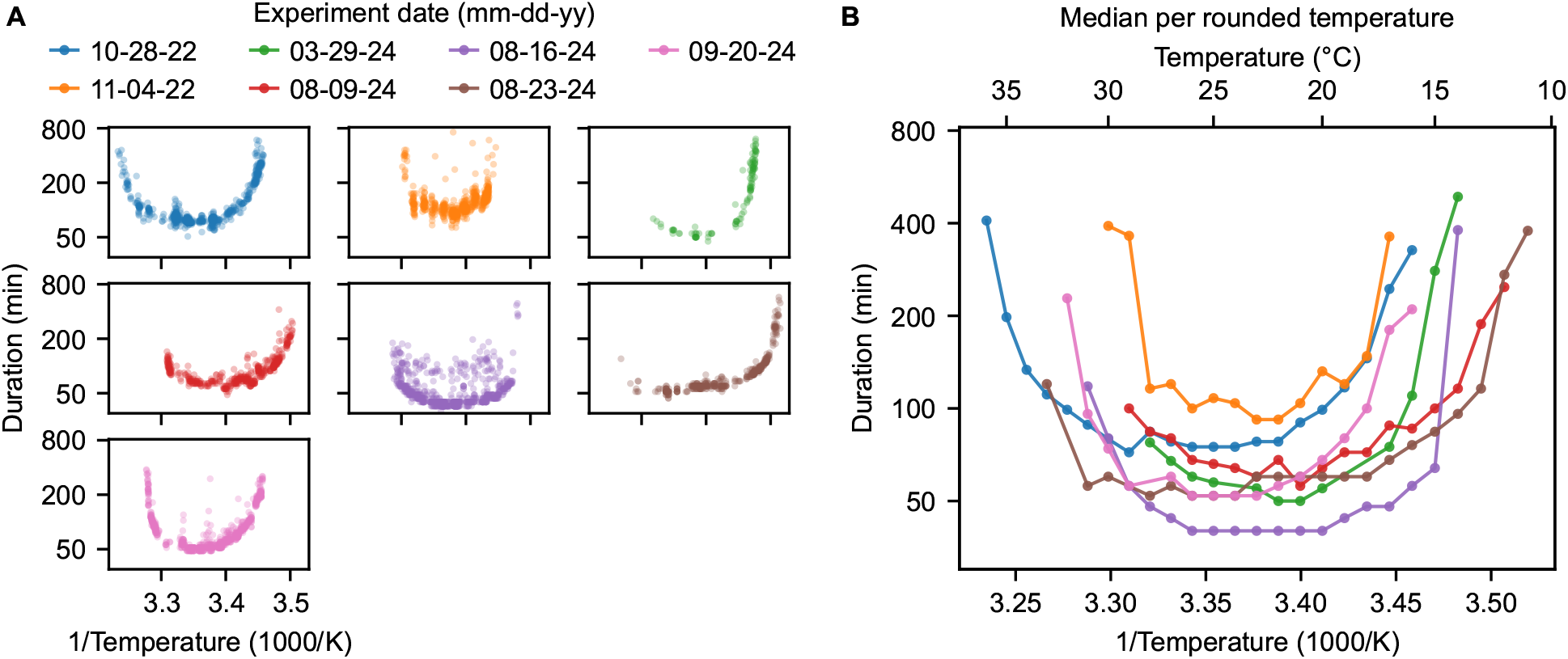
Comparison of temperature response across different replicates. A. Duration of the second cycle as a function of temperature for different biological replicates (different frogs and different experimental days). Dots represent individual droplet cycles. B. The median per rounded temperature of the datasets in panel A.

**Fig. S8:**
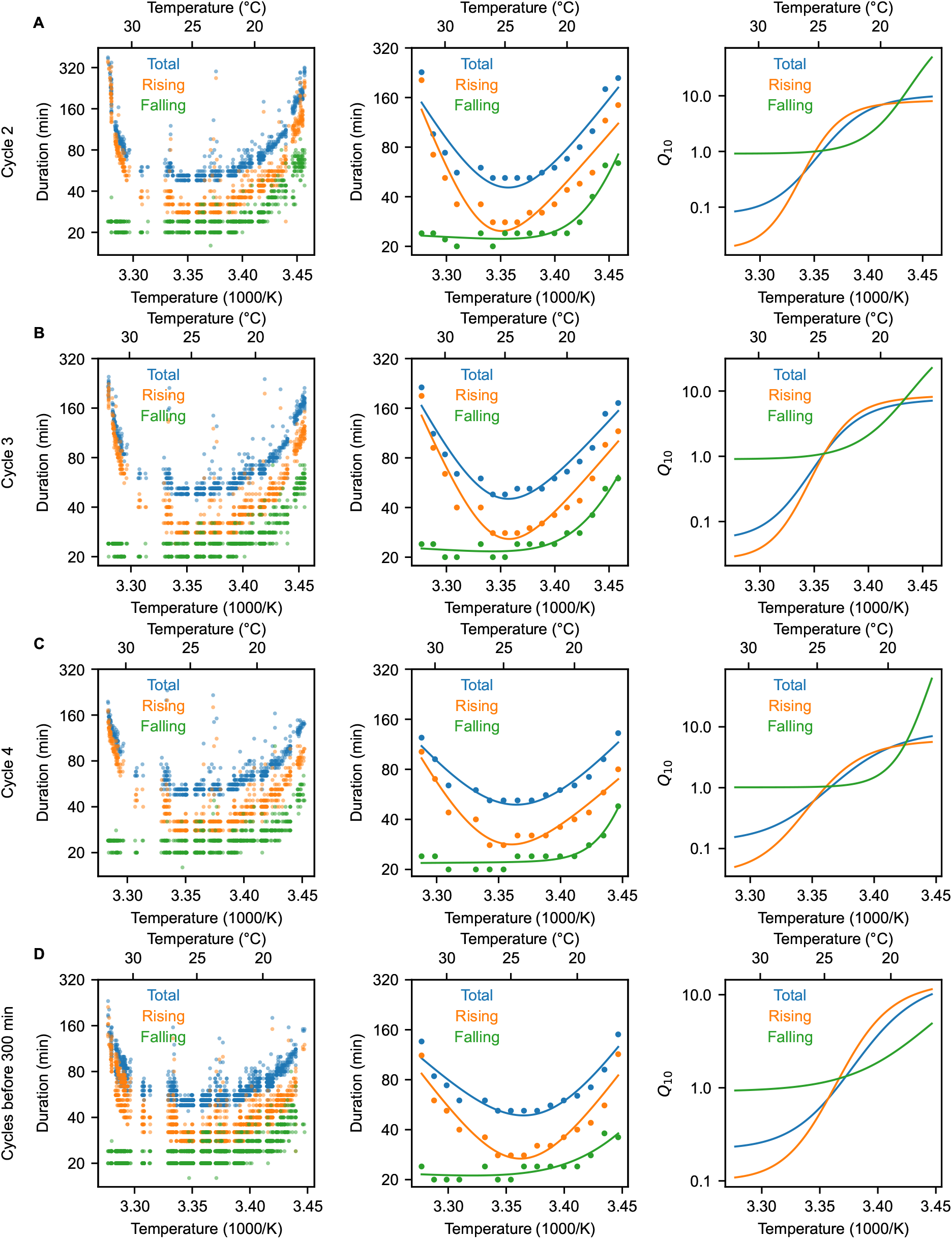
Scaling of the duration of the total cell cycle, rising phase, and falling phase for different cycles. Analogous to Fig. 4C-E: left the raw data, middle the median per rounded temperature with double exponential fit, right the local *Q*_10_ computed from the double exponential fit. A-C. The data for cycles 2, 3, 4 separately. D. Data from all the cycles that occur in the first 300 minutes of the experiment.

**Fig. S9:**
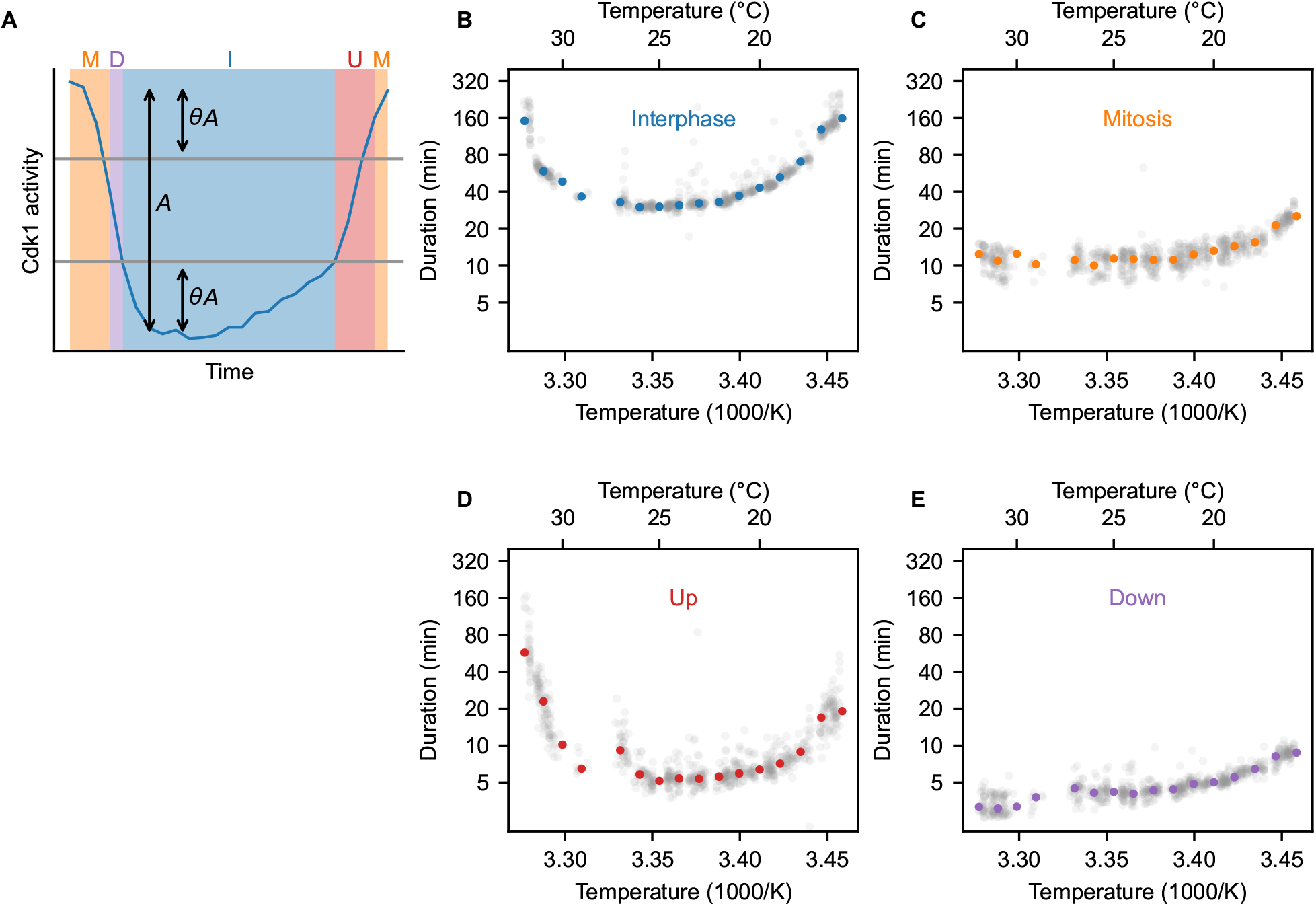
Scaling of four parts of the cycle in extracts. A. Diagram of the method for determining the duration of four parts of the cycle. After determining the minimal and maximal value of the cycle, we determine the amplitude *A* and two threshold values determined by *θ*, which we take to be 0.3. Duration of interphase is the time the cycle is below the lower threshold, mitosis is the time the cycle spends above the higher one, and up and down times are the times it spends in between. B-E Duration of these four parts as function of temperature. Light gray dots are datapoints from all droplets, colored dots show the median duration per temperature.

**Fig. S10:**
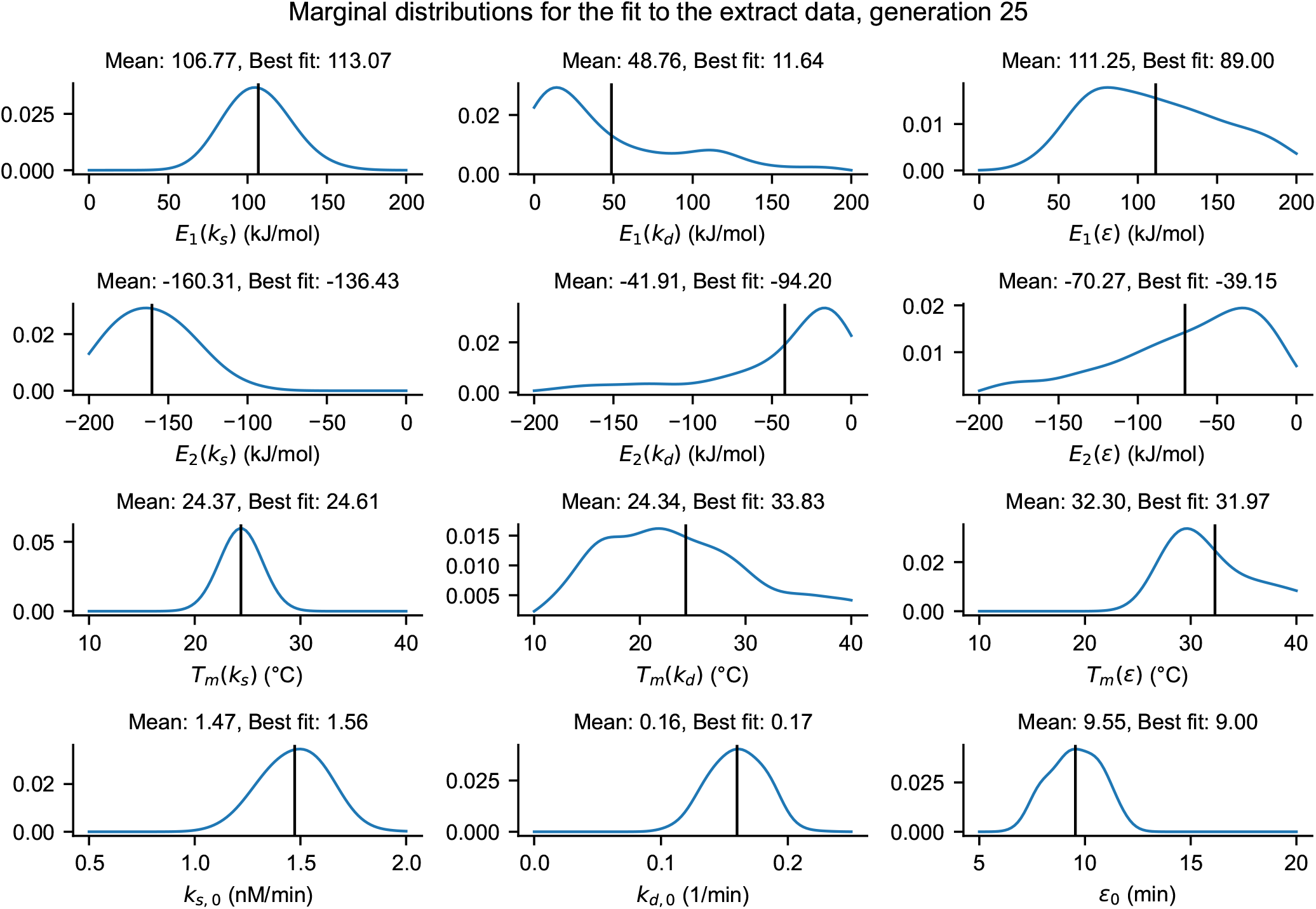
Marginal distributions of the different parameters for the optimal fits to extract data. These parameters determine the scaling of *k*_*s*_, *k*_*d*_ and *ϵ* that is shown in Fig. 4. The marginal distribution over 1000 weighted samples, that are the result of the ABC algorithm, is shown. Smooth distribution obtained by Gaussian kernel density. Black line indicates the mean. The ‘Best fit’ quoted corresponds to the value of the parameter for the sample with least distance to the data.

**Fig. S11:**
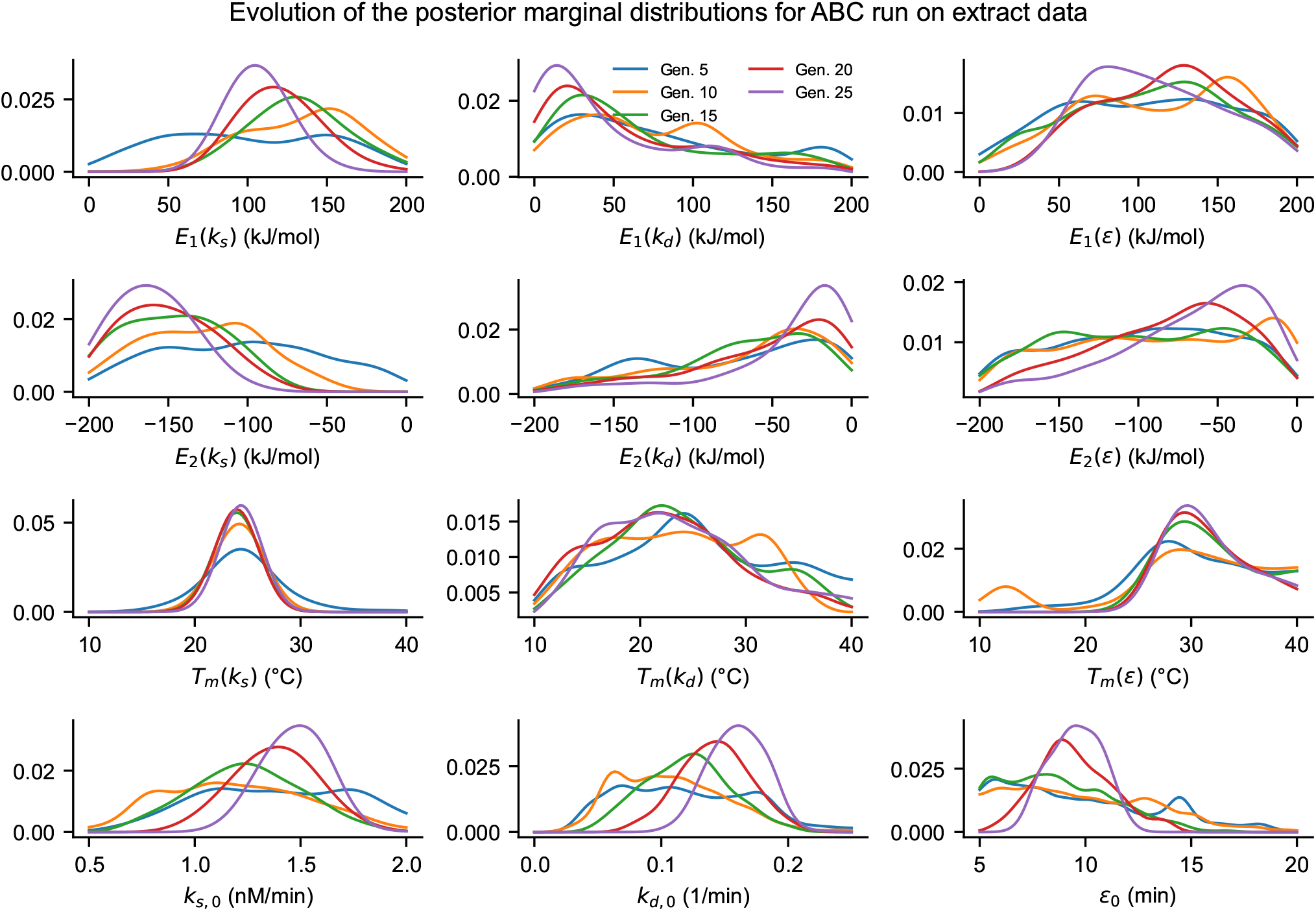
Evolution of the marginal distributions of the parameter sets over the course of the ABC algorithm. Similar to Fig. S10, but the distributions at different generations of the ABC algorithm are shown.

**Fig. S12:**
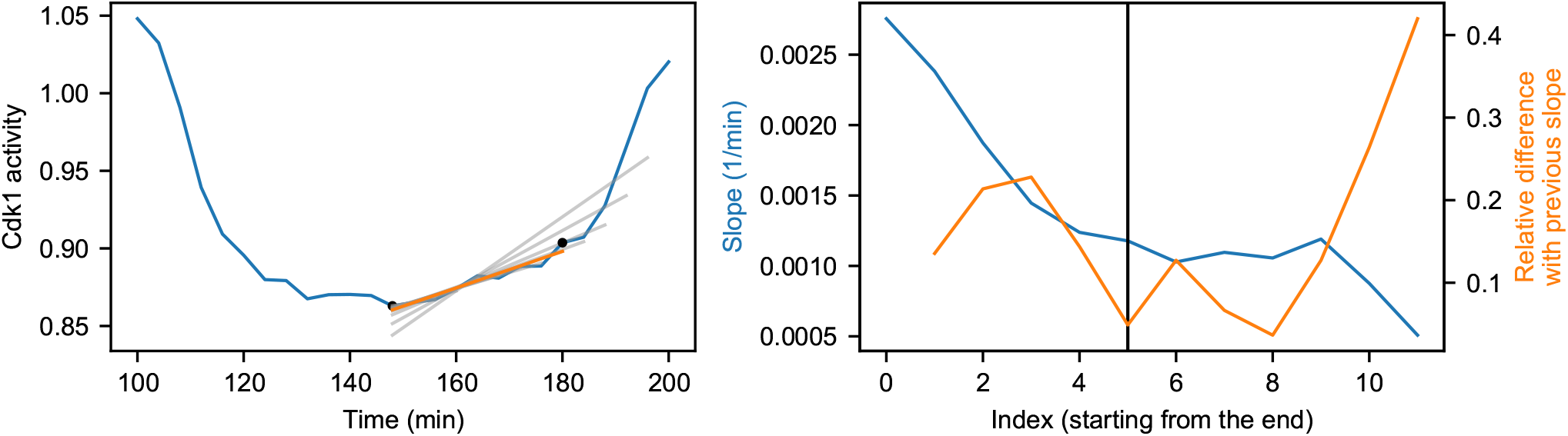
Determination of the cyclin synthesis rate. *k*_*s*_ **from the time series**. Shows what is explained in Supplementary Note 4.

**Fig. S13:**
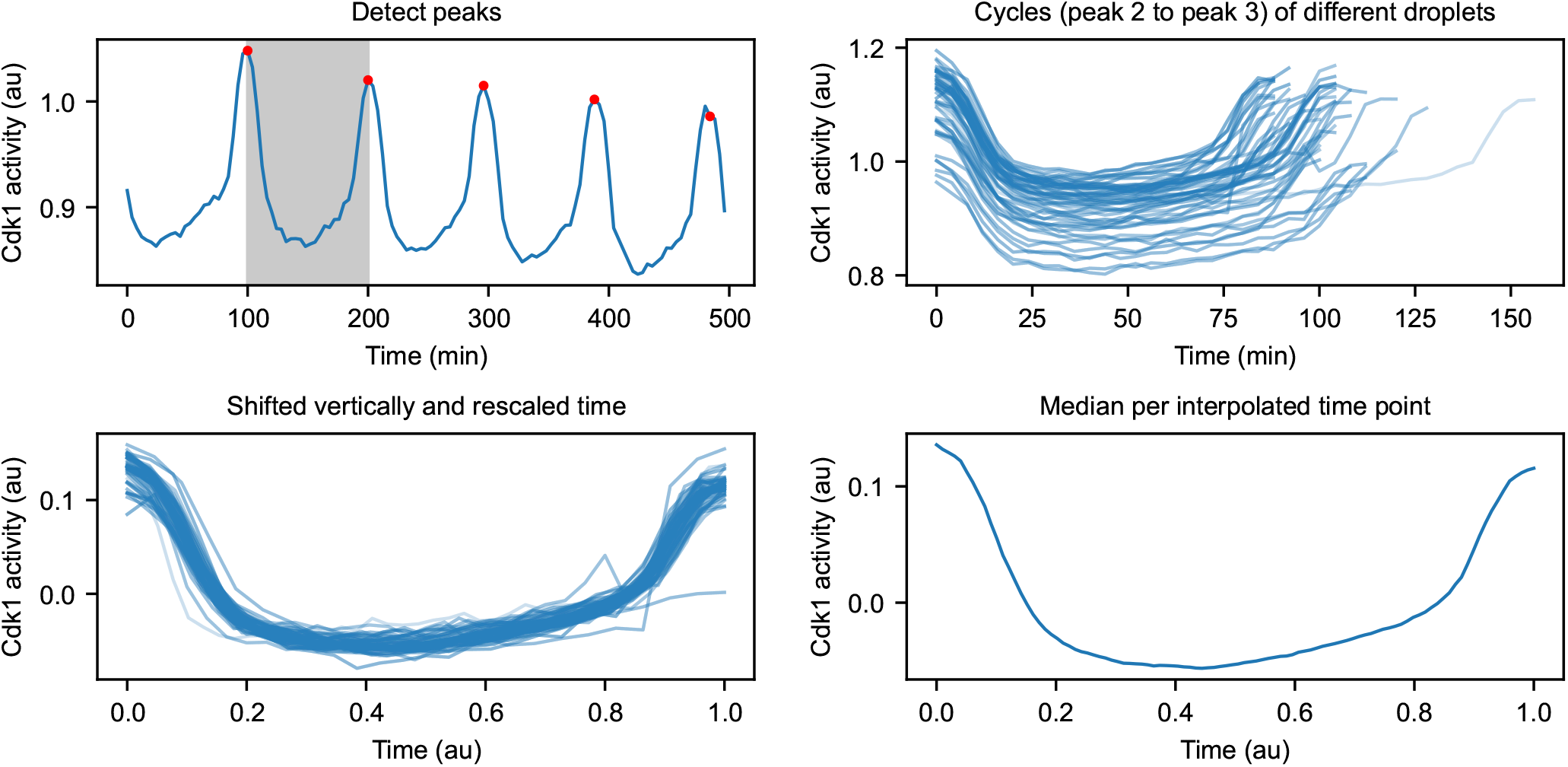
Determination of the average cycle shape. Shows what is explained in Supplementary Note 5. In this example, *T* = 22°C is shown. the bottom right panel shows the final average cycle.

**Fig. S14:**
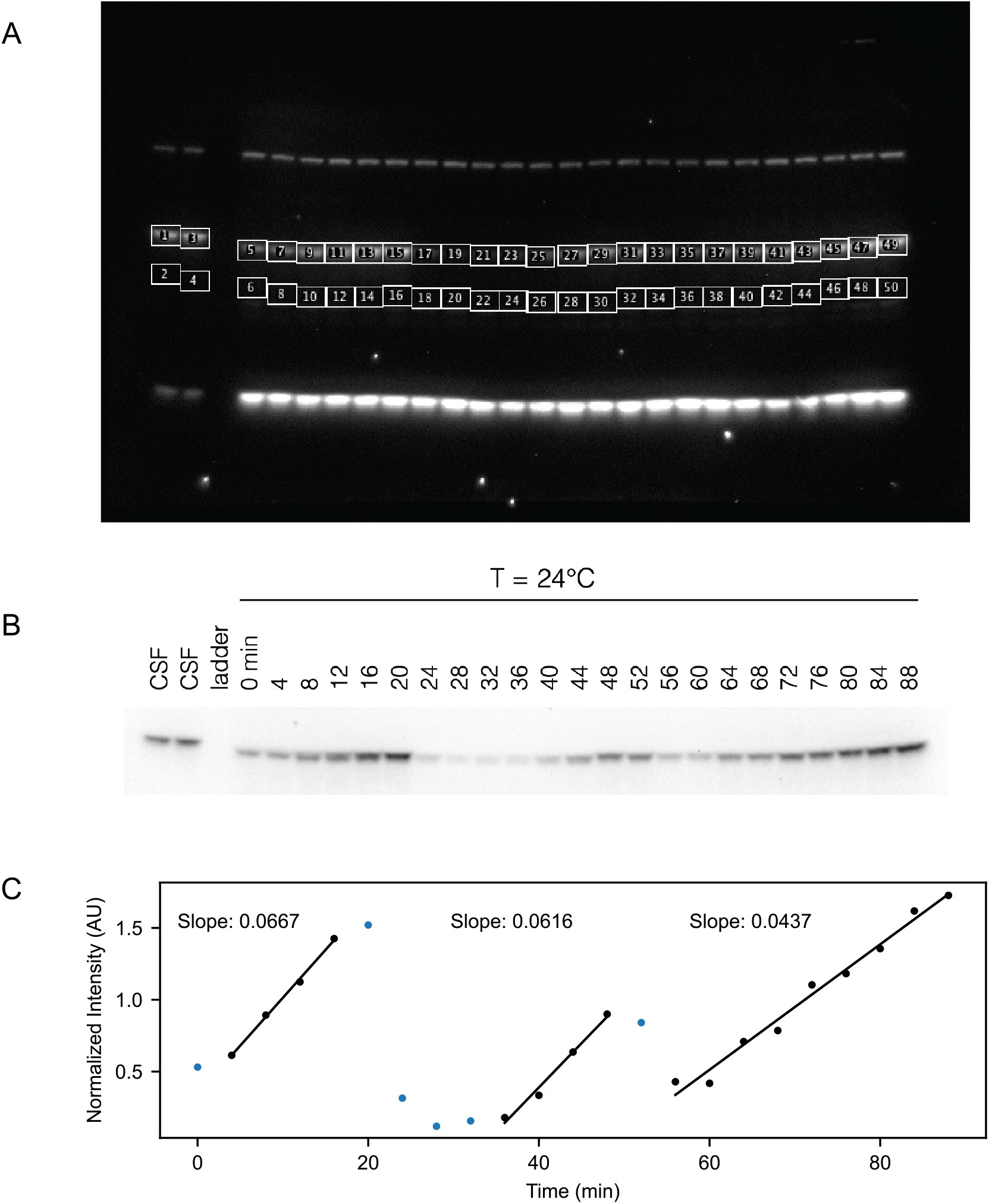
Measuring cyclin synthesis rates using quantitative Western blotting. A. Time course of a representative Western blot using anti-cyclin B2 antibody on a cycling frog egg extract. B. Selected region of same Western blot as in A. C. Quantification of the Western blot by calculating the integrated density for each band and subtracting the background using FIJI, we obtain the intensity for each time point. This value is then divided by the average intensity for a CSF extract, to obtain the normalized intensities. To obtain the cyclin synthesis rates, we next fit the slopes for each of the cycles for the points indicated in black.

**Fig. S15:**
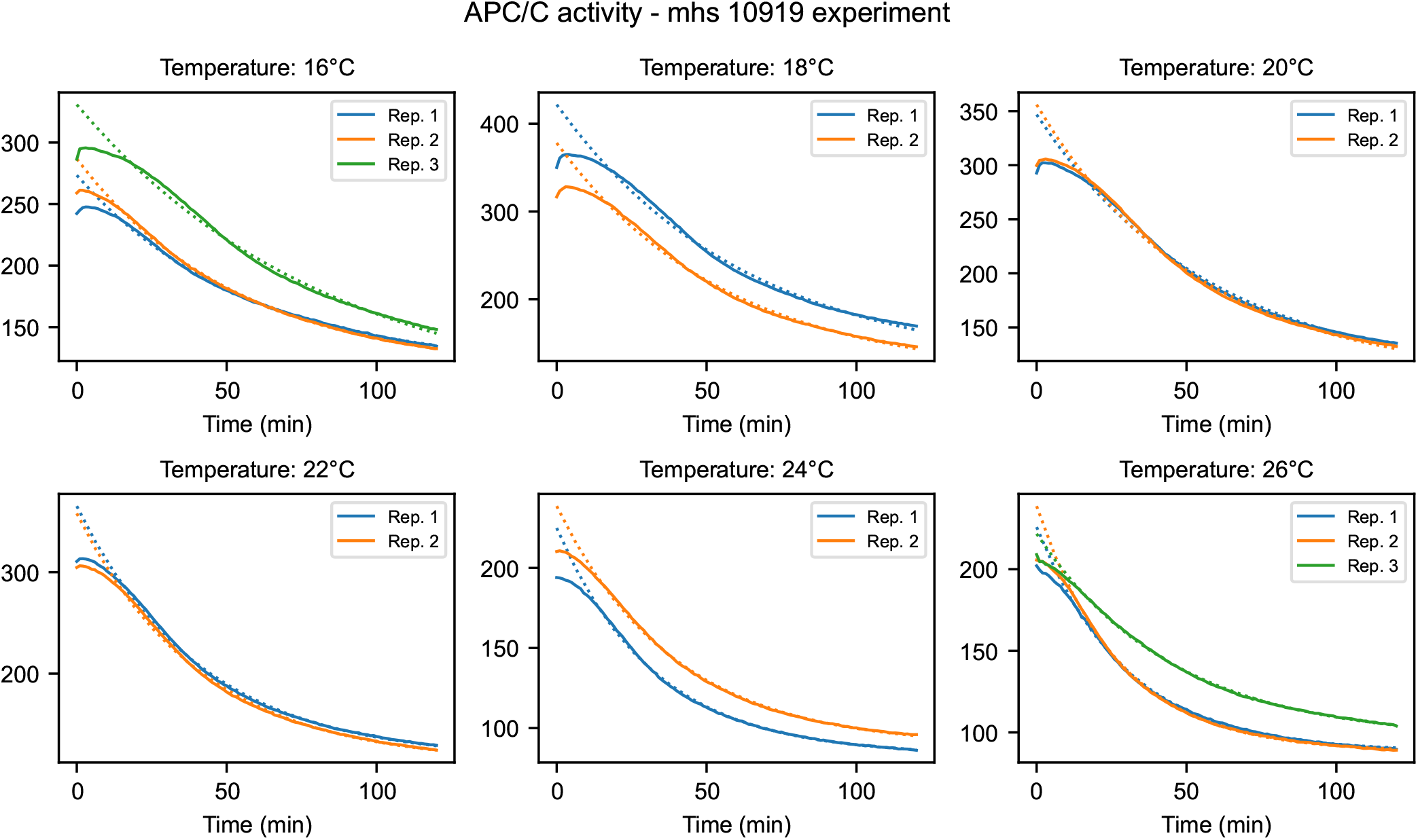
Time series for measuring APC activity. Time series and fits, from which the rates in Fig. 6 are obtained. Dotted lines are fits of the form *y* = *Ae*^−*kt*^ + *B*. Details in Supplementary Note 6.

**Fig. S16:**
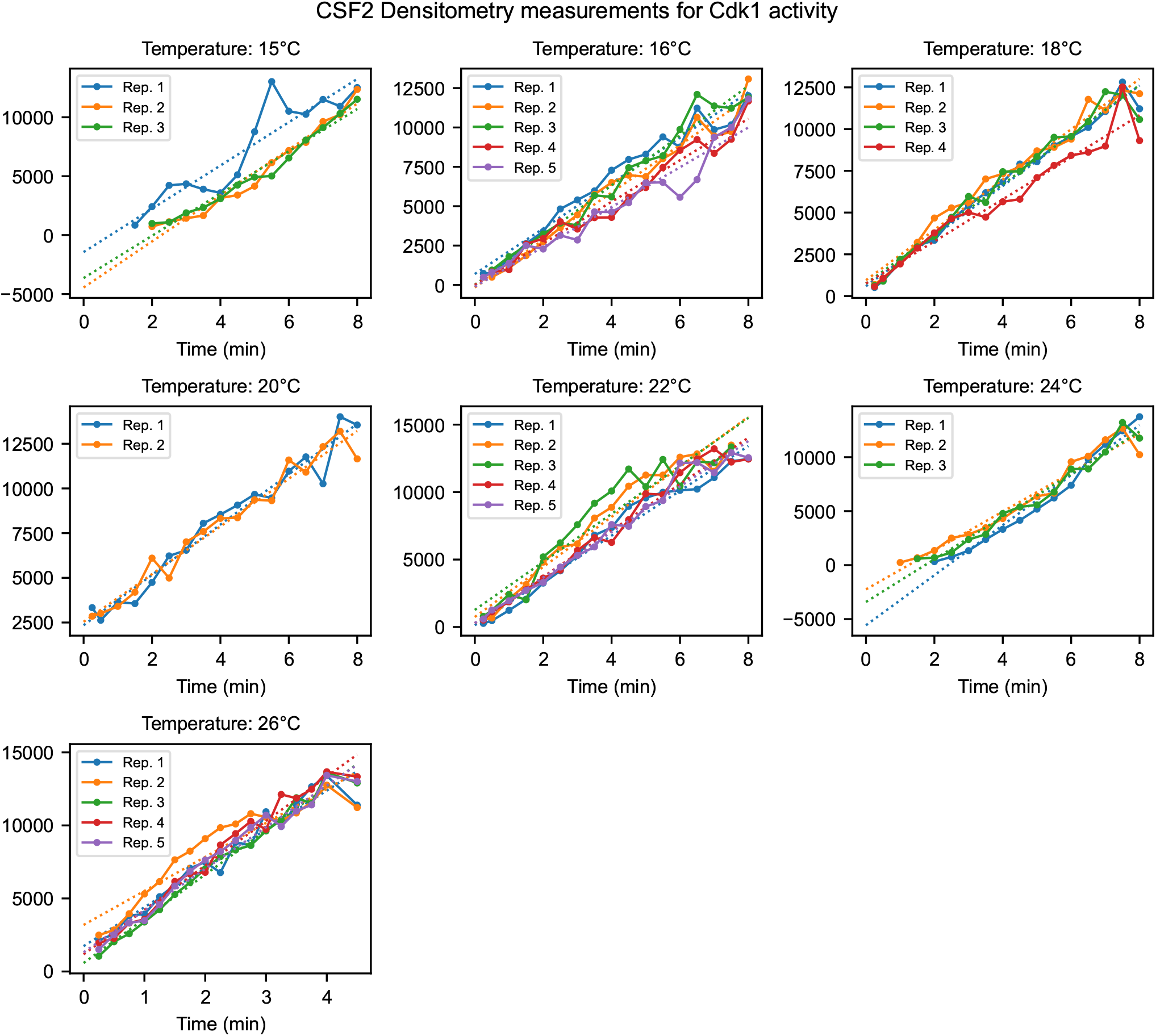
Time series for measuring Cdk1 activity. Time series and fits, from which the rates in Fig. 6 are obtained. Dotted lines are fits of the form *y* = *at* + *b*. Details in Supplementary Note 6.

**Fig. S17:**
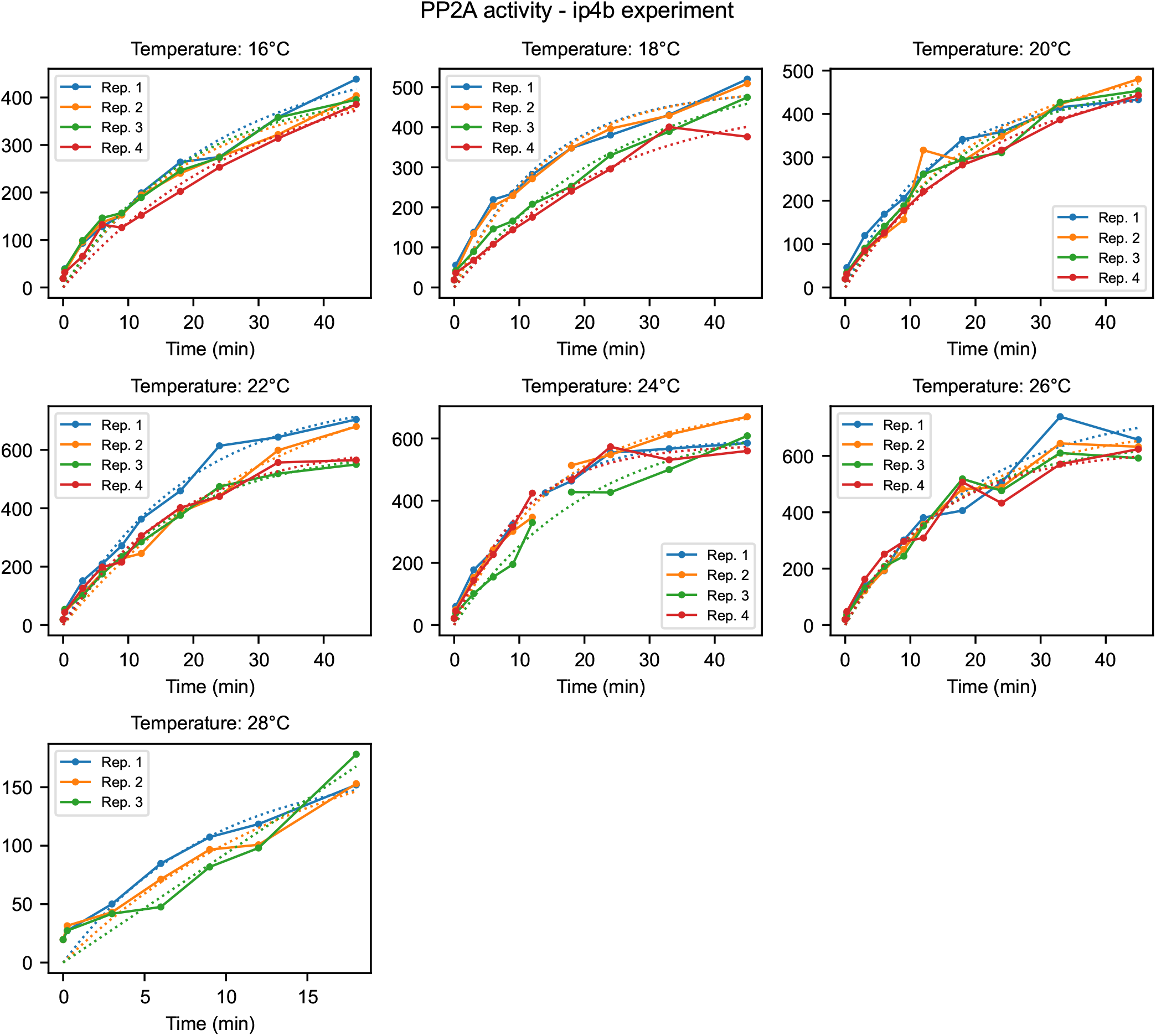
Time series for measuring PP2A activity. Time series and fits, from which the rates in Fig. 6 are obtained. Dotted lines are fits of the form *y* = *A*(1 −*e*^−*kt*^). Details in Supplementary Note 6.

**Fig. S18:**
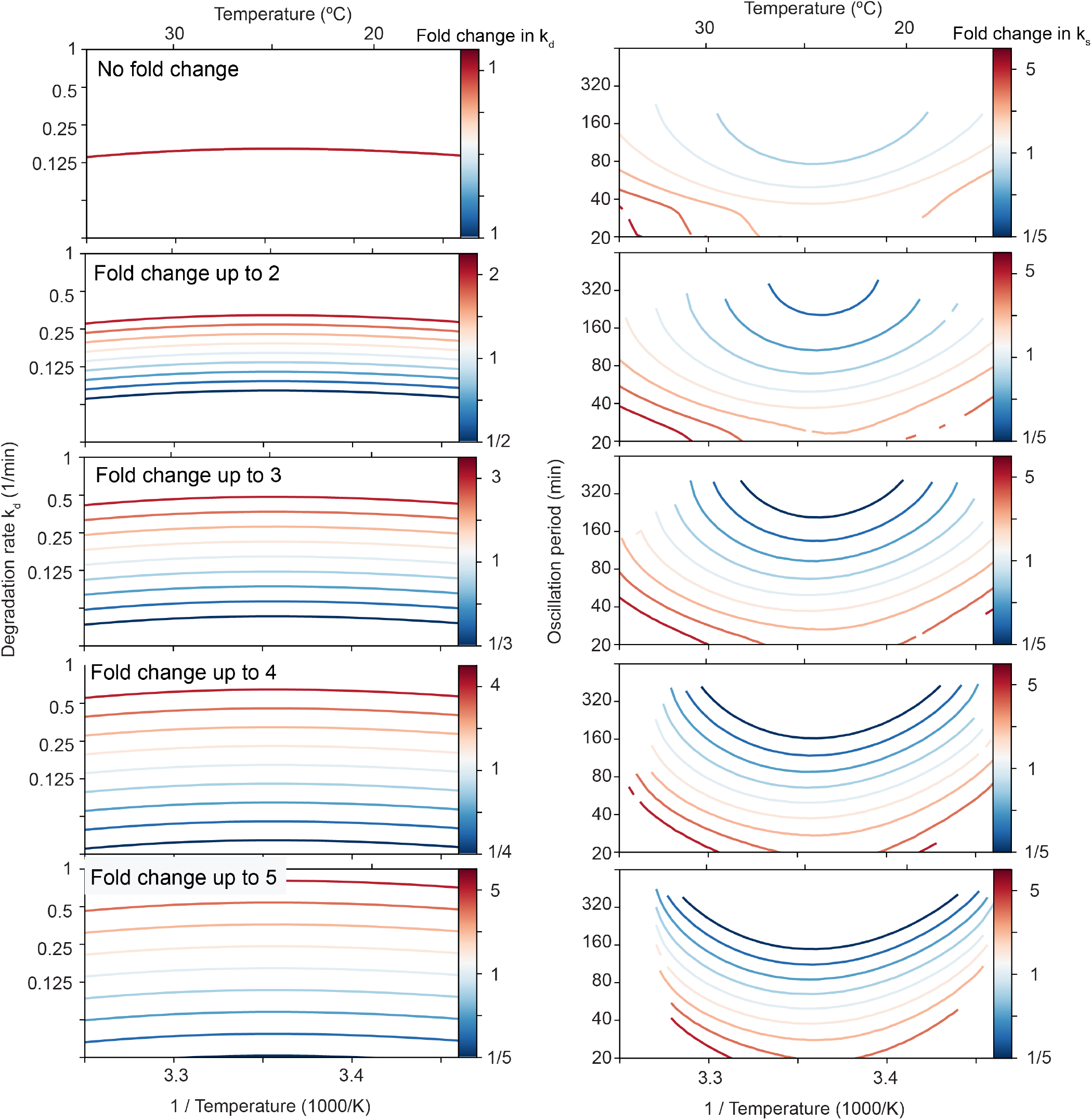
Decreasing the cyclin synthesis rate decreases the viable temperature range. Influence of changing the basal cyclin synthesis and the basal cyclin degradation rate by varying factors. In all simulations the basal cyclin synthesis rate is increased and decreased by a factor up to 5, similarly as shown in Fig. 7A. Additionally, from top to bottom we allow for increasing changes of the degradation rate as well. In the top panels, the basal degradation rate is kept constant as the basal cyclin synthesis rate is scaled. The lower panels show increasing fold changesin the basal degradation rate up to a scaling factor of 5, similar as for cyclin synthesis. Larger differences in cyclin synthesis and degradation rates (larger differences in scaling) lead to stronger reductions of the viable temperature range upon decreasing cyclin synthesis rate.

