## Supplemental notes and figures for "Mechanistic origins of temperature scaling in the early embryonic cell cycle"

#### Supplementary Note 1: Determination of local $E_a$ and $Q_{10}$

In the case of a duration or rate that scales according to the Arrhenius equation

$$k = Ae^{\frac{-E_a}{RT}} \quad \textbf{(8)}$$

the activation energy is a constant. The formula above is equivalent to

$$\ln k = \ln A - \frac{E_a}{R}\frac{1}{T}. \quad \textbf{(9)}$$

The value of $E_a$ can thus be calculated from the slope of the line obtained when plotting $\ln k$ vs $1/T$. Or,

$$E_a = -R\frac{d(\ln k)}{d(1/T)}. \quad \textbf{(10)}$$

This equation can also be used as the definition of a local activation energy $E_a(T)$ for any function $k(T)$.

Similarly, we can define a local $Q_{10}$ value. In this section we explain how a $Q_{10}$ value can be obtained for a process with temperature-dependent rate. The $Q_{10}$ is the fold change in rate when the temperature increases by 10 degrees Celsius. If the value of $Q_{10}$ is constant over all temperatures, the rate dependence should have the form

$$k \sim Q_{10}^{T/10}.$$

This form is different from the Arrhenius equation. It would therefore be incorrect to say of a process that its activation energy $E_a$ *and* its $Q_{10}$ are both constant over a range of temperatures.

The $Q_{10}$ can be calculated as

$$Q_{10} = \left(\frac{k(T_2)}{k(T_1)}\right)^{\frac{10}{T_2 - T_1}},$$

where $k(T_i)$ is the rate of the process calculated at temperature $T_i$. Since this formula holds for any choice of $T_1$ and $T_2$, we can look at the limit $T_2 \to T_1$ and use this formula to define a *local* $Q_{10}$, $Q_{10}(T)$. The use of 'local' for a number that is meant to convey what happens over a temperature change of 10 degrees is a bit contradictory, but we will make abstraction of this and use $Q_{10}(T)$ to indicate a local sensitivity to temperature.

If the rate of a process depends on temperature through any (differentiable) function $k(T)$, then for any $h$ we would have

$$Q_{10}(T) = \left(\frac{k(T+h)}{k(T)}\right)^{\frac{10}{h}}$$

A Taylor expansion for small $h$ gives that this is approximately equal to

$$\left(1 + h\frac{k'(T)}{k(T)}\right)^{\frac{10}{h}},$$

where $k' = \frac{dk}{dT}$. Using the definition of the exponential function, this goes to

$$\exp\left(10\frac{k'(T)}{k(T)}\right)$$

for small $h$. We thus define the local $Q_{10}$ as

$$Q_{10}(T) = e^{10\frac{k'(T)}{k(T)}}. \quad \textbf{(11)}$$

We can also express the formula for the local $E_a$ (Eq. (10)) using the derivative of the rate:

$$E_a(T) = -R\frac{d(\ln k)}{d(1/T)} = -R\frac{1}{k}\frac{dk}{d(1/T)} = R\frac{k'}{k}T^2.$$

This also gives a link between local $Q_{10}$ and $E_a$:

$$Q_{10}(T) = \exp\left(10\frac{E_a(T)}{RT^2}\right). \quad \textbf{(12)}$$

#### Supplementary Note 2: Bootstrapping

The histograms that describe the uncertainty on the fitted activation energies on the rates, shown in Figs. 1 and 6, were obtained using a bootstrapping procedure. First, we generated a new dataset by resampling the rate measurements *with replacement* from the original data. In this case, we used a stratified bootstrap: we resampled per temperature. So if in the original dataset, there were three values of $E_a$ for a given temperature, in a bootstrapped dataset there will also be three values for that temperature, and they are resampled with replacement from the original three. For each bootstrapped dataset, we computed the activation energy by fitting a straight line in the Arrhenius plot. We fitted on the median per temperature. We did this for 1000 bootstrapped datasets, and these 1000 values of $E_a$ are represented in the histograms. The Github repository contains the code to reproduce the bootstrapping procedure.

#### Supplementary Note 3: Computational cell cycle models

**3.A. Two-ODE cell cycle model.** As described in the main text, we made use of a two-ODE cell cycle model based on one originally described in (66):

$$\begin{aligned} \frac{d\text{cyc}}{dt} &= k_s - k_d d[\text{cdk1}_a]\text{cyc}, \\ \epsilon\frac{d\text{cdk1}_a}{dt} &= k_s - k_d d[\text{cdk1}_a]\text{cdk1}_a + k_a a[\text{cdk1}_a](\text{cyc} - \text{cdk1}_a) - k_i i[\text{cdk1}_a]\text{cdk1}_a, \end{aligned} \tag{13}$$

The first equation describes how cyclin B (cyc) is synthesized throughout the cell cycle at a rate $k_s$ (nM/min) and how it is degraded at a rate $k_d$ (1/min) by the proteasome after ubiquitination by active APC/C. APC/C is assumed to be activated instantaneously by active cyclin B - Cdk1 complexes (abbreviated as $\text{cdk1}_a$) in an ultrasensitive way (given by $d[\text{cdk1}_a]$). The second ODE describes the time evolution of active cyclin B - Cdk1 complexes, assuming that all synthesized cyclin B quickly binds to Cdk1 to form a complex. Moreover, the positive feedback of Cdk1 via Cdc25, and the double negative feedback of Cdk1 via Wee1 are included as an activating ($a[\text{cdk1}_a]$) and inhibiting ($i[\text{cdk1}_a]$) ultrasensitive function as well, motivated by direct experimental measurements of those response functions (67, 68). The different ultrasensitive functions have the following form:

$$a[x] = a_{\text{Cdc25}} + b_{\text{Cdc25}} \frac{x^{n_{\text{Cdc25}}}}{\text{EC}_{50,\text{Cdc25}}^{n_{\text{Cdc25}}} + x^{n_{\text{Cdc25}}}}, \tag{14}$$

$$i[x] = a_{\text{Wee1}} + b_{\text{Wee1}} \frac{\text{EC}_{50,\text{Wee1}}^{n_{\text{Wee1}}}}{\text{EC}_{50,\text{Wee1}}^{n_{\text{Wee1}}} + x^{n_{\text{Wee1}}}}, \tag{15}$$

$$d[x] = a_{\text{APC}} + b_{\text{APC}} \frac{x^{n_{\text{APC}}}}{\text{K}_{\text{APC}}^{n_{\text{APC}}} + x^{n_{\text{APC}}}}. \tag{16}$$

This type of model has previously been used to successfully describe various aspects of cell cycle oscillations (34, 66, 73). In the present work, we simplified this model further by omitting the degradation term from the Cdk1 equation, to have a clearer separation of the first ODE with cyclin synthesis and degradation, and the second ODE just describing Cdk1 activation and inactivation processes. This simplification does not significantly influence the results or our conclusions.

$$\begin{aligned} \frac{d\text{cyc}}{dt} &= k_s - k_d d[\text{cdk1}_a]\text{cyc}, \\ \epsilon\frac{d\text{cdk1}_a}{dt} &= k_a a[\text{cdk1}_a](\text{cyc} - \text{cdk1}_a) - k_i i[\text{cdk1}_a]\text{cdk1}_a, \end{aligned} \tag{17}$$

Using experimentally motivated parameters (66), model (17) reproduces cell cycle oscillations with a period of approx. 30 min (Fig. 3C). These oscillations manifest as a closed trajectory, a limit cycle, in the (cyc, $\text{cdk1}_a$) phase plane (Fig. 3B, red). The phase-plane picture helps to better understand the existence of the oscillations via the intersection of nullclines (NCs). NCs are defined by points where $\frac{d\text{cyc}}{dt} = 0$ (Cyc NC) or $\frac{d\text{cdk1}_a}{dt} = 0$ (Cdk1 NC). When $\epsilon \ll 1$, oscillations occur at the intersection of the cyclin NC and the middle branch of the S-shaped Cdk1 NC (as depicted in Fig. 3B).

| Symbol | Meaning | Value |
|---|---|---|
| $k_s$ | Cyclin production rate | 1.25 nM/min |
| $k_d$ | Cyclin degradation rate | 0.1 $\text{min}^{-1}$ |
| $k_i$ | Maximal Wee1 activity | 1 $\text{min}^{-1}$ |
| $k_a$ | Maximal Cdc25 activity | 1 $\text{min}^{-1}$ |
| $a_{\text{Cdc25}}$ | Basal Cdc25 activity | 0.2 |
| $b_{\text{Cdc25}}$ | Maximal increase in Cdc25 activity | 0.8 |
| $K_{\text{Cdc25}}$ | Threshold for Cdc25 activation | 30 nM |
| $n_{\text{Cdc25}}$ | Hill exponent for Cdc25 activation | 10 |
| $a_{\text{Wee1}}$ | Basal Wee1 activity | 0.1 |
| $b_{\text{Wee1}}$ | Maximal increase in Wee1 activity | 0.4 |
| $K_{\text{Wee1}}$ | Threshold for Wee1 activation | 30 nM |
| $n_{\text{Wee1}}$ | Hill exponent for Wee1 activation | 5 |
| $a_{\text{APC}}$ | Basal APC/C activity | 0.1 |
| $b_{\text{APC}}$ | Maximal increase in APC/C activity | 0.9 |
| $n_{\text{APC}}$ | Hill exponent for APC/C activation | 15 |
| $K_{\text{APC}}$ | Threshold for APC/C activation | 30 nM |
| $\epsilon$ | Timescale parameter | 0.1 |

Table 1: Parameter values used in the 2-ODE cell cycle model. These are the basal values that correspond roughly to the period of the *X. laevis* and *D. rerio* cell cycle. To describe the *X. tropicalis cycle*, which is faster, we multiply $k_s$ and $k_d$ by 1.3. The basal values of $k_s$, $k_d$ and $\epsilon$ are also different for extracts: these values are obtained as part of the fitting to the time series.

##### 3.B. Five-ODE mass action model.

***3.B.1. The model.*** We used a five-equation model for relaxation oscillations arising out of the interaction between Cdk1, Greatwall and PP2A (Fig. S6A). This pathway is a different part of the mitotic control system, which underlies the second mitotic switch. In the two-ODE model used in the main text, Cdk1 is involved in two feedback loops, through Wee1 and Cdc25. These feedbacks lead to bistability of Cdk1 activity as function of cyclin B levels. In the five-equation model, these feedback loops are not present, as is appropriate for cycles 2–12 in embryos (73). Here, any cyclin B-Cdk1 complex is directly activated assuming Cdk1 is present in excess over cyclin B: the response curve of Cdk1 activity as function of cyclin B levels would be linear. The production rate of cyclin B therefore directly corresponds to the production rate of active cyclin B-Cdk1 complexes. The equations we use are based on the paper by Hopkins et al. (72), who show that the equations for Greatwall, ENSA and PP2A lead to bistability. We complement their system with equations for Cdk1 and APC/C to turn the bistable system into a relaxation oscillator (77) (Fig. S6B).

We use the following equations:

$$\begin{aligned}
u' &= k_{p,a}(A_T - u)v - k_{d,a}u(P_T - y)\\
v' &= k_s - k_d uv\\
w' &= k_{p,g}(G_T - w)v - k_{d,g}(P_T - y)w\\
x' &= -k_{p,e}xw + k_{\text{cat}}y\\
y' &= k_{\text{ass}}(E_T - x - y)(P_T - y) - k_{\text{diss}}y - k_{\text{cat}}y.
\end{aligned} \tag{18}$$

Here, $u$ denotes active APC/C, $v$ are the active cyclin B-Cdk1 complexes. The variable $w$ corresponds to phosphorylated Greatwall, $x$ is free, unphosphorylated ENSA and $y$ is the complex ENSA-PP2A. The parameters of the model are the biochemical rates and the total amounts of APC/C ($A_T$), Greatwall ($G_T$), ENSA ($E_T$), and PP2A ($P_T$).

The first equation describes the activity of APC/C. We assume that APC/C can be converted from its inactive (unphosphorylated) form to its active (phosphorylated) form by Cdk1 through mass-action kinetics. The dephosphorylation is performed by PP2A. Note that $y$ is the concentration of the ENSA-PP2A complex and $P_T$ is total PP2A, such that the available PP2A is $P_T - y$. The second equation describes cyclin B-Cdk1 levels. These are governed by production and degradation of cyclin B, the latter of which is modeled through mass action. The equation for active Greatwall (variable $w$) describes the conversion between active and inactive Greatwall by Cdk1 and PP2A respectively — analogous to the APC/C equation. The fourth equation describes free unphosphorylated ENSA. This concentration decreases through phosphorylation of ENSA by Greatwall, and it increases through the dephosphorylation, which is mediated by PP2A. The final equation describes the concentration of the ENSA-PP2A complex.

The parameters, their meaning and their standard values can be found in Table 2. The parameter set is not based on experimental values, but was chosen to obtain a relaxation oscillation of amplitude (in the Cdk1 variable) and period that correspond to observations. All variables except for cyclin B-Cdk ($v$) are in arbitrary units, this is why the units of the rate constants look a bit awkward. As for the 2-ODE model, $k_s$ and $k_d$ are multiplied by 1.3 if simulations are to be compared with data from *X. tropicalis*.

| Symbol | Meaning | Value |
|---|---|---|
| $k_{p,a}$ | phosphorylation rate of APC/C by Cdk1 | 0.4 nM$^{-1}$min$^{-1}$ |
| $k_{d,a}$ | dephosphorylation rate of APC/C by PP2A | 100 min$^{-1}$ |
| $k_{p,g}$ | phosphorylation rate of Greatwall by Cdk1 | 0.06 nM$^{-1}$min$^{-1}$ |
| $k_{d,g}$ | dephosphorylation rate of Greatwall by PP2A | 20 min$^{-1}$ |
| $k_{p,e}$ | phosphorylation rate of ENSA by Greatwall | 6 min$^{-1}$ |
| $k_{\text{ass}}$ | association rate of phosphorylated ENSA and PP2A | 100 min$^{-1}$ |
| $k_{\text{diss}}$ | dissociation rate of ENSA-PP2A complex | 1 min$^{-1}$ |
| $k_{\text{cat}}$ | rate of catalyzed dephosphorylation of ENSA | 4.5 min$^{-1}$ |
| $k_s$ | Cyclin production rate | 1.5 nM/min |
| $k_d$ | Cyclin degradation rate | 0.15 min$^{-1}$ |
| $A_T$ | Total APC/C in the system | 1 |
| $G_T$ | Total Greatwall in the system | 1 |
| $E_T$ | Total ENSA in the system | 3 |
| $P_T$ | Total PP2A in the system | 1 |

Table 2: Parameter values used in the mass action model.

***3.B.2. Interpretation in the phase plane.*** The model with only Greatwall, ENSA and PP2A ($w, x, y$) has been shown to produce a bistable response as function of the amount of active Cdk1 ($v$) (72). In our version, we added production of cyclin B and its degradation through ubiquitination by APC/C to turn the bistable system into a relaxation oscillator. Even though the system is five-dimensional, we can understand it in the phase plane. To do this, we perform a reduction to a two-variable system. We assume a quasi-steady-state condition on $w, x$ and $u$, and set their derivatives to zero. When we do this, we assume that these variables evolve on a faster timescale than the others. We do expect that the levels of cyclin B-Cdk1 evolve on a slower timescale than the other variables: production and degradation are slower than the phosphorylation and dephosphorylation reactions. The reasons for keeping $y$ as the additional variable and not one of the others is more practical: taking $w' = x' = u' = 0$ leads to explicit expressions for these variables as function of $v$ and $y$, which does not work if we take, say, $u$ as remaining variable. We find

$$\begin{aligned} w(v,y) &= \frac{1}{1+\frac{k_{d,g}(P_T-y)}{k_{p,g}v}}G_T \\ x(v,y) &= \frac{k_{\text{cat}}y}{k_{p,e}w} = \frac{k_{\text{cat}}y}{k_{p,e}G_T}\left(1+\frac{k_{d,g}(P_T-y)}{k_{p,g}v}\right) \\ u(v,y) &= \frac{1}{1+\frac{k_{d,a}(P_T-y)}{k_{p,a}v}}A_T. \end{aligned} \quad \textbf{(19)}$$

Using these, we can reduce the system to two equations:

$$\begin{aligned} v' &= k_s - k_d u(v,y)v \\ y' &= k_{\text{ass}}(E_T - x(v,y) - y)(P_T - y) - k_{\text{diss}}y - k_{\text{cat}}y. \end{aligned} \quad \textbf{(20)}$$

Fig. S6C shows the phase plane of this system with the associated limit cycle. The projection of the solution of the five-ODE system for the same parameter values is also shown. The two limit cycles are close in the phase plane, but their period is significantly different (Fig. S6D). This reduction shows that we can qualitatively understand the oscillations of this system in the phase plane. In particular, we confirm that the limit cycle is of relaxation type and goes around the underlying bistable switch. The cycle is, as in the 2-ODE model, driven by cyclin B accumulation and degradation. Once cyclin B-Cdk1 levels cross a threshold, the activity of the phosphatase is quickly suppressed. This allows activation of APC/C, which leads to cyclin degradation and brings the system back to a state of low Cdk1 activity. In particular, we can see from Eq. (20) that the production and degradation rates affect the non-S-shaped nullcline only.

For this model, we studied how the period of the oscillation changes if each of the parameters has an Arrhenius-like dependence on temperature. We analyzed how different activation energies for the different parameters can lead to different scaling as well as thermal ranges. As for the Yang-Ferrell model, this analysis can be interpreted in the phase plane, by examinining how parameter changes affect the location of the nullclines. From the phase-plane picture we can see that, when the oscillations disappear, the system becomes stuck in a state with either high or low phosphatase activity. As before, which one it is will depend on the relative magnitude of $E_a(k_s)$ and $E_a(k_d)$.

The steady state of the system only depends on the ratios of the following parameters:

$$\frac{k_s}{k_d}, \quad \frac{k_{p,a}}{k_{d,a}}, \quad \frac{k_{p,g}}{k_{d,g}}, \quad \frac{k_{p,e}}{k_{\text{cat}}} \text{ and } \frac{k_{\text{ass}}}{k_{\text{cat}} + k_{\text{diss}}}.$$

It follows that, for any set of activation energies such that

$$E_a(k_s) = E_a(k_d), \qquad E_a(k_{p,a}) = E_a(k_{d,a}),$$
$$E_a(k_{p,g}) = E_a(k_{d,g}), \qquad E_a(k_{p,e}) = E_a(k_{\text{cat}}) = E_a(k_{\text{ass}}) = E_a(k_{\text{diss}}),$$

the steady state of the system is independent of temperature. Under the assumption that the relative magnitude of the timescales stays the same, this means that we would expect oscillations over a large range of temperatures if these rates scale in a similar way. These ratios usually have the rates for two counteracting reactions in numerator and denominator.

In Fig. 5 we show the results of fitting the parameters to the time scaling of extract data. These fits were obtained using Approximate Bayesian Computation - Sequential Monte Carlo (ABC-SMC) (74). This algorithm sequentially samples parameter sets that provide better and better fits to the data. The output of the algorithm is $N$ parameter sets $\Theta_i$ with associated weights $w_i$. Each of these parameter sets provides a fit closer than a prescribed distance $\varepsilon$ to the data. The $N$ weighted parameter sets constitute a sample from the posterior distribution $P(\Theta \mid d(x^*, x_0) < \varepsilon)$, where $x_0$ is the data, $x^*$ is the data resulting from a simulation with parameters $\Theta$ and $d$ is a distance function. If $\varepsilon$ is small, this distribution approximates the posterior distribution $P(\Theta \mid x_0)$: the probability that a parameter set $\Theta$ is the true one, given the observed data. In ABC-SMC, the value of $\varepsilon$ is lowered over the course of different generations as a way of getting better and better approximations of the posterior. We use the implementation of this algorithm given in pyABC (75).

For the extract fits, we describe the temperature scaling of each of the rates $k_s$, $k_d$ and $\epsilon$ using a double-exponential formula. For $k_s$ and $k_d$, we parametrize the rate as

$$\text{rate} = \left(A_1 e^{\frac{E1}{RT}} + A_2 e^{\frac{E2}{RT}}\right)^{-1},$$

and for $\epsilon$, which has units of duration and not rate, we use

$$\epsilon = A_1 e^{\frac{E1}{RT}} + A_2 e^{\frac{E2}{RT}}.$$

Instead of the four parameters $A_1, E_1, A_2, B_2$, we decide to use more interpretable parameters: the basal value of the parameter at 18 degrees Celsius, the temperature $T_m$ at which is maximal (minimal for $\epsilon$) value is obtained, and the two activation energies $E_1$ and $E_2$. Note that we can map $k_0, T_m, E_1, E_2$ to $A_1, E_1, A_2, E_2$ directly.

A parameter vector $\Theta$ contains 12 values $[E_1(k_s), E_2(k_s), T_m(k_s), k_{s,0}, E_1(k_d), E_2(k_d), T_m(k_d), k_{d,0}, E_1(\epsilon), E_2(\epsilon), T_m(\epsilon), \epsilon_0]$. Each parameter set thus defines three functions $k_s(T)$, $k_d(T)$ and $\epsilon(T)$. We simulate the 2-ODE model over the temperatures from the dataset, using the rates defined by these scaling functions. All the other model parameters are kept to their basal value (Table 1). For each temperature, we detect whether the system is oscillating using the peaks of the time series, as described in Methods. For oscillating systems, we then use the *cdk* timeseries to determine the rising (min to max) and falling (max to min) durations.

The output of the simulation is thus $X_{\text{simulation}} = \{(R_i, F_i), i = 1, \ldots N\}$: the duration of rising and falling part of the cycle, for each temperature $T_i$ in the dataset ($N$ being the total number of temperature points). These data from the simulation are then compared to the same data obtained from the extract time series $X_{\text{data}}$. The distance function we use is

$$d(X_{\text{data}}, X_{\text{simulation}}) = \frac{1}{N} \sum_i \left(|\ln F_{\text{data},i} - \ln F_{\text{simulation},i}| + |\ln R_{\text{data},i} - \ln R_{\text{simulation},i}|\right). \quad \textbf{(21)}$$

We thus consider the differences of the logarithms of the rising and falling times, for each temperature, and take the average of their absolute values. The durations can vary quite a bit in absolute value, and we use the logarithms to prevent the algorithm being skewed to approximating the large durations (at extreme temperatures). We take $d(X_{\text{data}}, X_{\text{simulation}}) = \infty$ if there is a temperature in the dataset for which the simulation did not produce an oscillation. The consequence is that we only search for parameter values for which the model produces oscillations over *at least* the range we see in the experiment.

We compare the distance between the simulated dataset $X_{\text{simulation}} = \{P_i, i = 1 \ldots N\}$ (the durations of the embryonic cycles for all temperatures $T_i$) and the observed data $X_{\text{data}}$with the distance function

$$d(X_{\text{data}}, X_{\text{simulation}}) = \frac{1}{N} \sum_i |\ln P_{\text{data},i} - \ln P_{\text{simulation},i}|. \tag{22}$$

The distance is set to infinity if there is at least one temperature $T_i$ for which the simulation does not produce an oscillation.

The basal rates at 18°C are as in Table 1, but for *X. tropicalis* we multiply $k_s$ and $k_d$ by 1.3 to account for its faster cycle. Since the temperature scaling of a rate is defined by

$$\text{rate} = \text{basal rate} \times e^{\frac{-E_a}{R}\left(\frac{1}{T} - \frac{1}{T_0}\right)},$$

1. We determine all the indices $i$ that belong to one cycle, which is defined as all the datapoints between two peaks: we select$(t_i, u_i)$ for $i_1 \leq i \leq i_2$. Here $i_1$ and $i_2$ are the indices corresponding to peaks in the Cdk1 signal.
2. We rescale time to the interval $[0,1]$: set $\tilde{t}_i = (t_i - t_{i_1})/(t_{i_2} - t_{i_1})$ and we shift the values of $u$ vertically by subtracting the mean: $\tilde{u}_i = u_i - \bar{u}_i$.
3. We interpolate the values of $\tilde{u}$ at 100 evenly spaced time points in the interval [0,1], obtaining a new time series $(\hat{t}_i, \hat{u}_i)$, $i = 1 \ldots 100$ with $t_1 = 0$ and $t_{100} = 1$.
4. We do this for all the droplets that have a given temperature. This yields $(t_{k,i}, u_{k,i})$ where $k$ indexes the different time series (droplets). The average shape of the time series is then obtained by taking, for each $i$, the median of the values of $u$. The resulting time series is $(t_i, U_i)$ with $U_i = \text{median}_{\text{over } k}\, u_{k,i}$

#### Supplementary Note 6: Fitting rate measurements of individual regulatory processes

We describe how the rates for cyclin B synthesis, Cdk1 activity, PP2A activity and APC/C activity were fitted from time series to obtain Fig. 6. For the linear fit we used the numpy function `polyfit`, and for the nonlinear fits we used the function `curve_fit` from the `scipy.optimize` package.

##### 6.A. Cyclin B synthesis rates.

Fig. S14A-B shows a representative Western blot. By calculating the integrated density for each band and subtracting the background using FIJI, we obtain the intensity for each time point. This value is then divided by the average intensity for a CSF extract, to obtain the normalized intensities as shown in Fig. S14C. To obtain the cyclin accumulation rates, we next fit the slopes for each of the cycles, e.g. for the points indicated in S14C. The obtained results for the cyclin accumulation rates $k_s$ from all Western blots, obtained from three independent extracts, are summarized in Fig. S8A (left).

##### 6.B. Cdk1 rates.

The time series from which we derive the rates are shown in Fig. S16. We fit a function of the form

$$y = at + b$$

to the time series, and save $a$ as the resulting rate. We do this for every replicate, giving us multiple rate measurements per temperature. We used the non-P32 decay adjusted rates.

##### 6.C. PP2A rates.

The time series from which we derive the rates are shown in Fig. S17. We fit a function of the form

$$y = A(1 - e^{-kt})$$

to the time series. The rate is determined as the derivative of this function at $t = 0$, i.e. $kA$.

##### 6.D. APC rates.

The time series from which we derive the rates are shown in Fig. S15. We fit a function of the form

$$y = Ae^{-kt} + B$$

to the time series where we impose that $A, B, k > 0$. The rate we save is $k$. For the fit, we only use datapoints for times larger than or equal to 10 minutes, because the exponential decay does not start in the beginning.

### Supplemental Figures

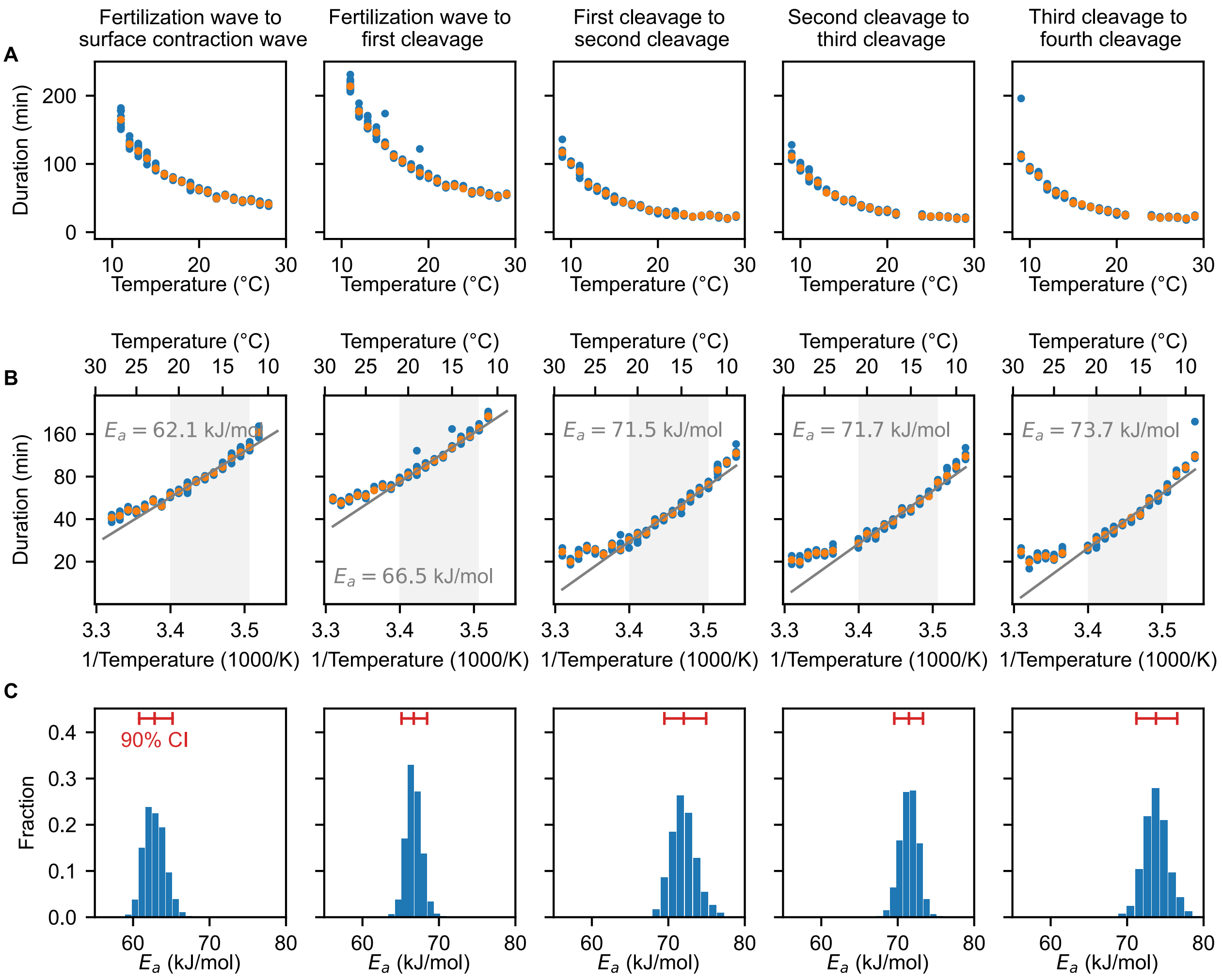


Fig. S1: **Cell division timing in early Xenopus laevis embryos scales approximately Arrhenius over a wide range of temperatures.** A. Duration of several early developmental periods in function of temperature in the range [$T_{\min} = 9°C$, $T_{\max} = 29°C$]. B. An Arrhenius fit is shown for the values between 12°C and 21°C, with the apparent activation energy indicated. C. Bootstrapping provides a probability distribution for the apparent activation energies. The mean and 90% confidence interval (CI) are also indicated.

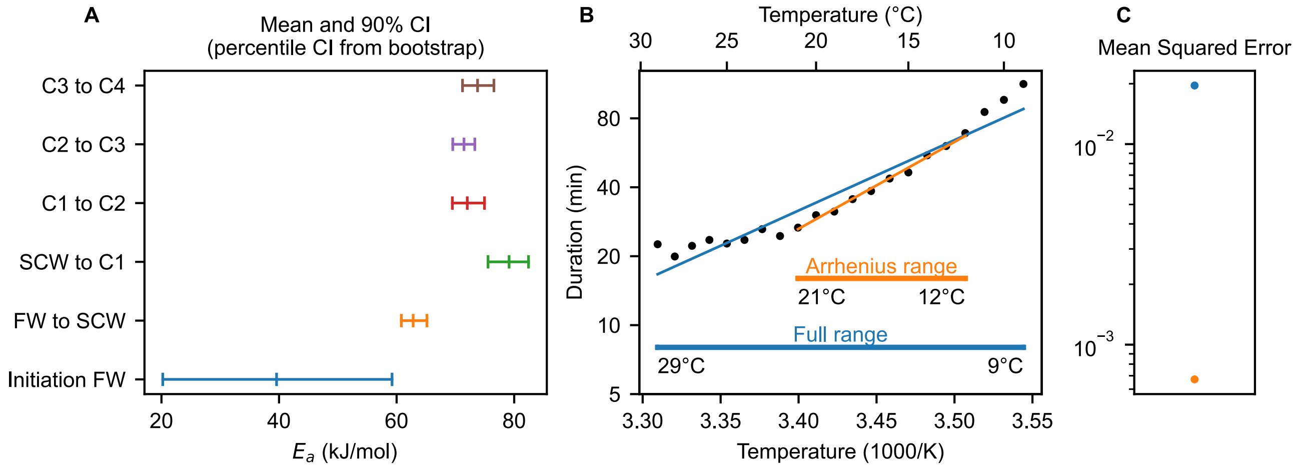


Fig. S2: **Cell division timing in early Xenopus laevis embryos does not scale Arrhenius over the whole temperature range** A. In Fig. 1C, we show the duration of several early developmental periods in function of temperature in the range $[T_{\min} = 9°C, T_{\max} = 29°C]$, with the apparent activation energy as obtained by an Arrhenius fit between 12°C and 21°C in Fig. 1D. Bootstrapping provides a probability distribution for the apparent activation energies (Fig. 1E). Here, we show the mean and 90% confidence interval (CI) for comparison. FW is fertilization wave, SCW is surface contraction wave, C means cleavage. B. Cleavage cycle duration in function of temperature for the second to fourth cell cycle in the range $[T_{\min} = 9°C, T_{\max} = 29°C]$ for *Xenopus laevis*. Optimal fits using single exponential Arrhenius (SE) are shown in two different temperature ranges: from 12°C and 21°C (orange), and the whole temperature range (blue). The mean square error (MSE) is much higher over the whole temperature range than within the selected range (panel C), indicating that the Arrhenius equation does not fit the data well over the whole measured data range.

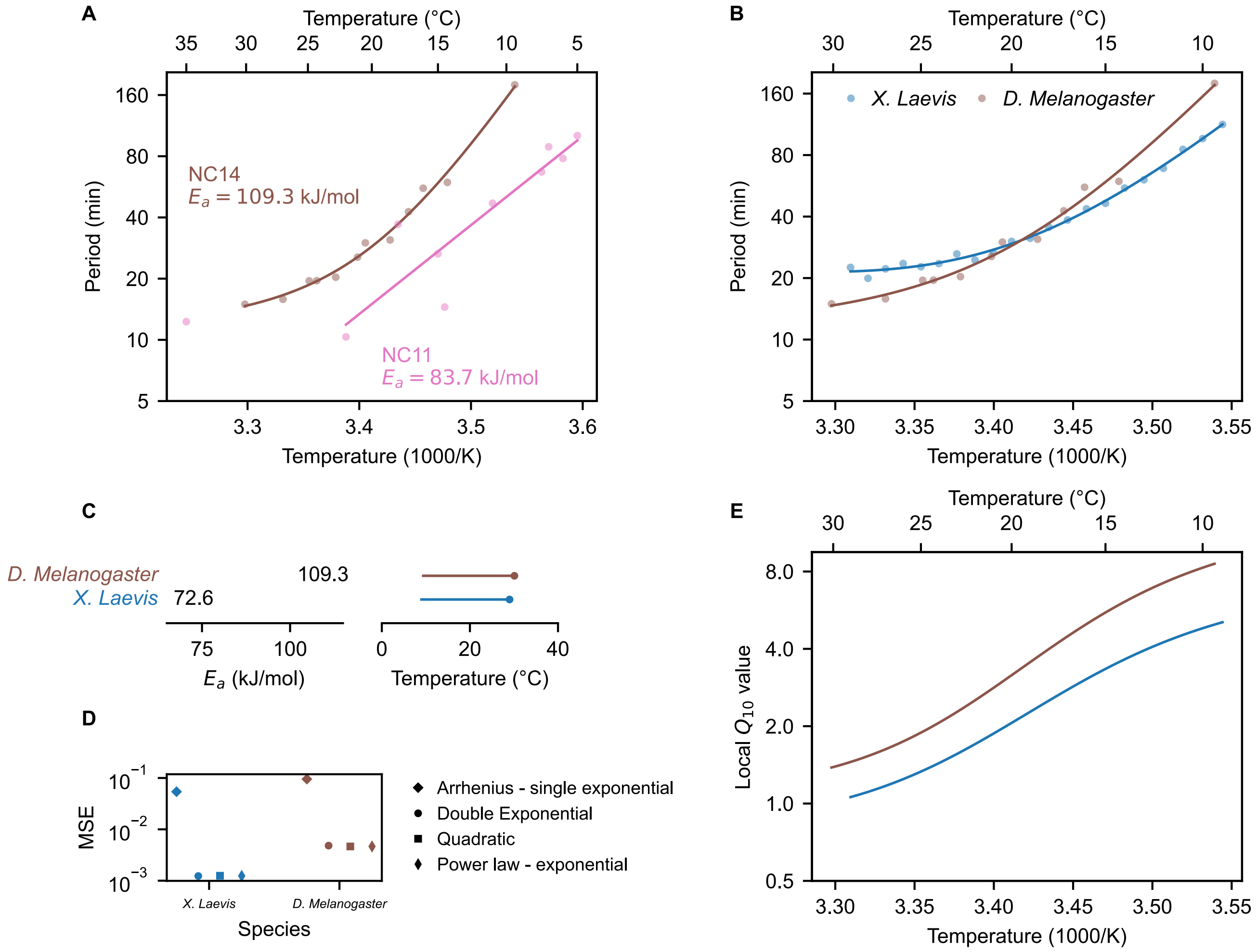


Fig. S3: **Temperature scaling of embryonic processes in *Drosophila melanogaster*** A. Median cleavage period in function of temperature for the eleventh and thirteenth cell cycle *D. melanogaster*. Data for NC11 from Falahati et al. (33), for NC14 from Crapse et al. (32). Optimal fits using a double exponential (DE) function are overlayed. B. Median cleavage period in function of temperature for the second to fourth cell cycle (all pooled) in *X. laevis* (this work), and for the eleventh and fourteenth cell cycle *D. melanogaster* (32), together with double exponential (DE) best fits. C. Activation energy, minimal and maximal temperature for *X. laevis* and *D. melanogaster*, corresponding to the curves in panel B. The dot shows the optimal temperature, which in this case is also the maximal temperature. D. Comparison of the mean squared error (MSE) for fits with different functional forms, as in Fig. 2G. E. Using the best DE fit, the local $Q_{10}$ value is plotted as function of temperature.

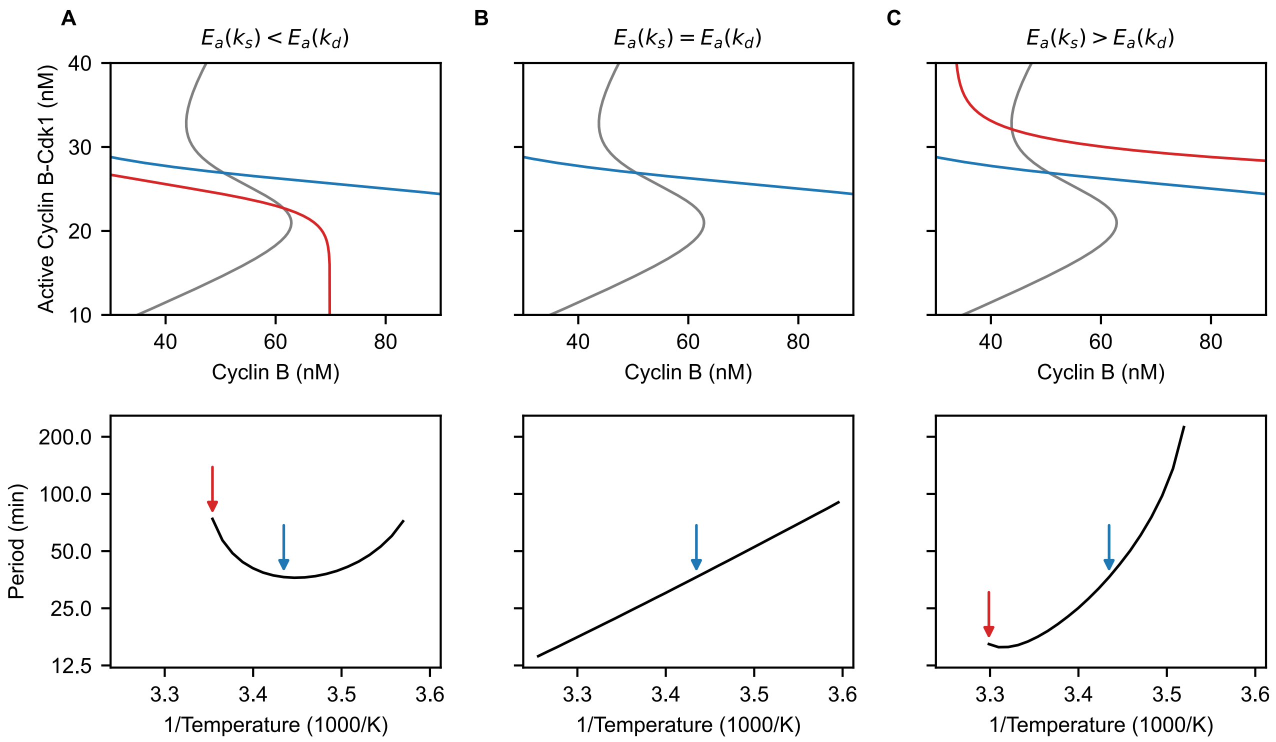


Fig. S4: **Temperature dependence of nullclines in the phase plane.** In the two-ODE cell cycle model, thermal limits are determined by intersection of nullclines. Depending on the relative size of $E_a(k_s)$ and $E_a(k_d)$, the non-S-shaped nullcline (red/blue) shifts upward or downward with rising temperatures. When the intersection of the nullclines lies on the upper or lower branch of the S-shaped nullcline (gray), oscillations cease to exist. A. The system ends up in a low-activity state at high temperatures. B. The second nullcline is temperature independent. There is no thermal limit. C. The system is stuck in a high-activity state at high temperatures.

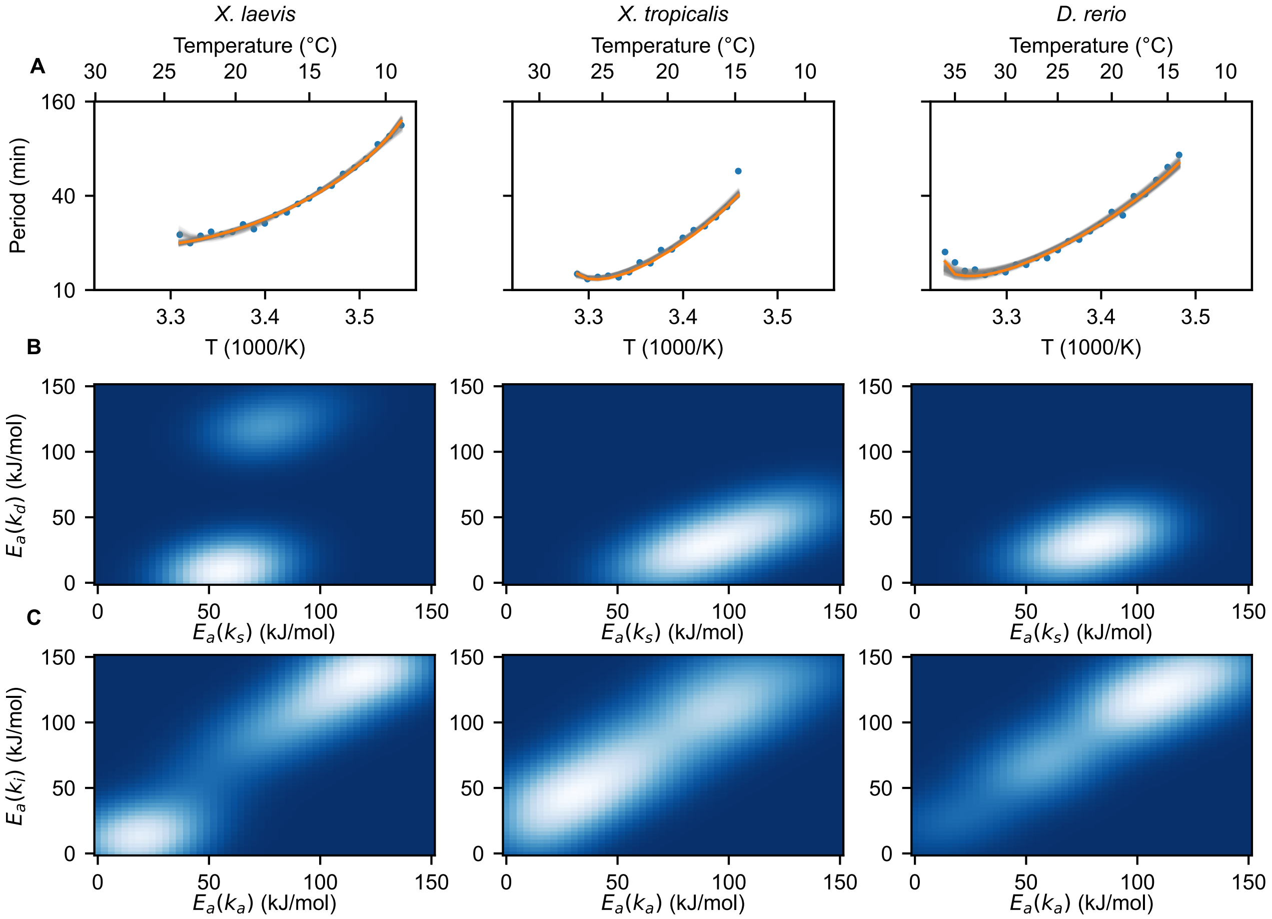


Fig. S5: **Optimal fits of the two-ODE model to the measured data using the ABC method.** A. Resulting fits from the ABC algorithm for the two-ODE model. Gray lines show the 200 resulting parameter sets, with more transparent lines corresponding to points with lower weight. The orange line is the best fit (smallest distance). B. Projection of the four-dimensional probability density onto the ($E_a(k_s)$ , $E_a(k_d)$)-plane. C. Projection onto the ($E_a(k_a)$, $E_a(k_i)$)-plane. The heatmaps were constructed from 200 weighted samples and smoothed with a Gaussian kernel.

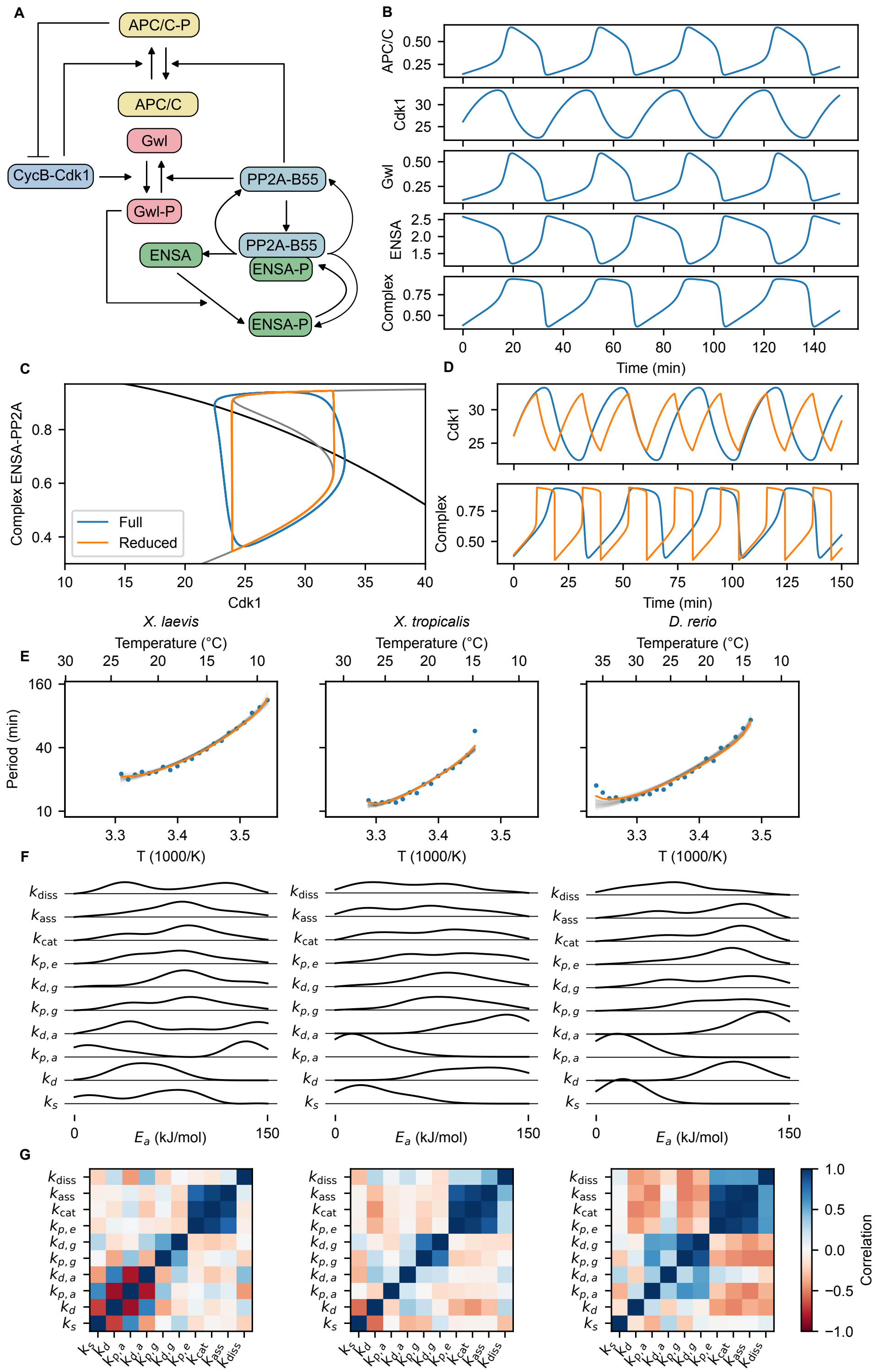


Fig. S6: **The five-ODE mass action model and fits to the measured data using ABC method.** A. The interaction diagram for the five-ODE mass action model. B. Representative time series of a simulation of the 5-equation model. C. Phase plane of the reduced two-ODE model and projection of the five-ODE model onto this plane. Nullclines are shown in black and gray, the blue limit cycle is the projection of the oscillation of the five-ODE system and the orange limit cycle is the one in the two-ODE system. D. Time series of corresponding variables in the full (blue) and reduced (orange) model. E. Resulting fits from the ABC algorithm for the mass action model. Gray lines show the 200 resulting parameter sets, with more transparent lines corresponding to lower-weighted points. The orange line is the best fit (smallest distance). F. Results of the ABC algorithm for the mass-action model. Marginal distributions of the different activation energies (smooth distribution obtained using Gaussian kernel density). G. Pairwise correlation of the activation energies of the different rates computed from the result of the ABC algorithm.

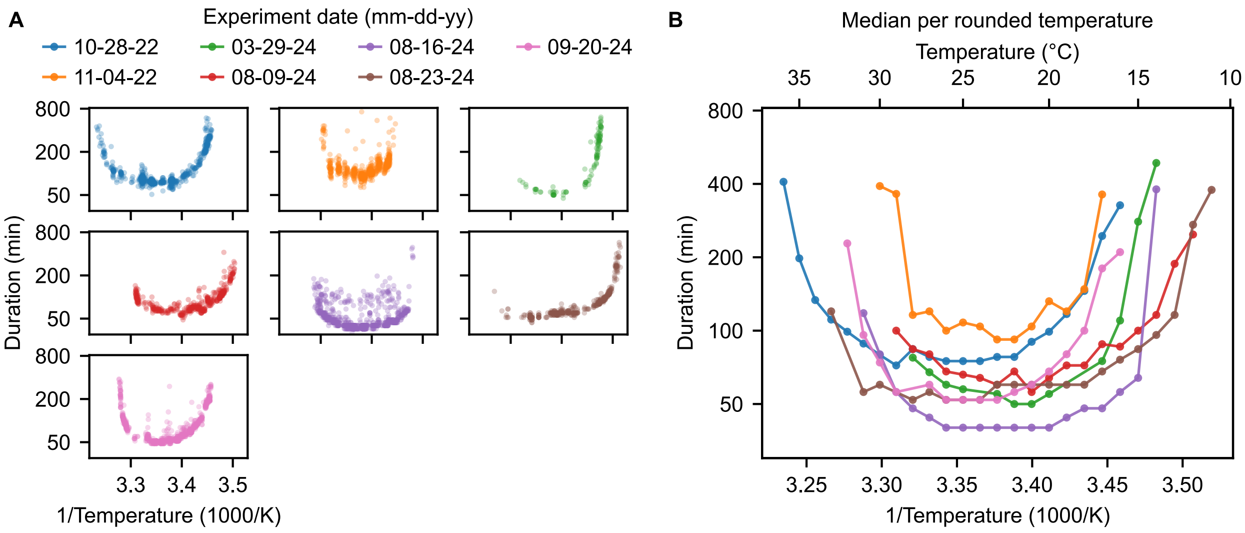


Fig. S7: **Comparison of temperature response across different replicates.** A. Duration of the second cycle as a function of temperature for different biological replicates (different frogs and different experimental days). Dots represent individual droplet cycles. B. The median per rounded temperature of the datasets in panel A.

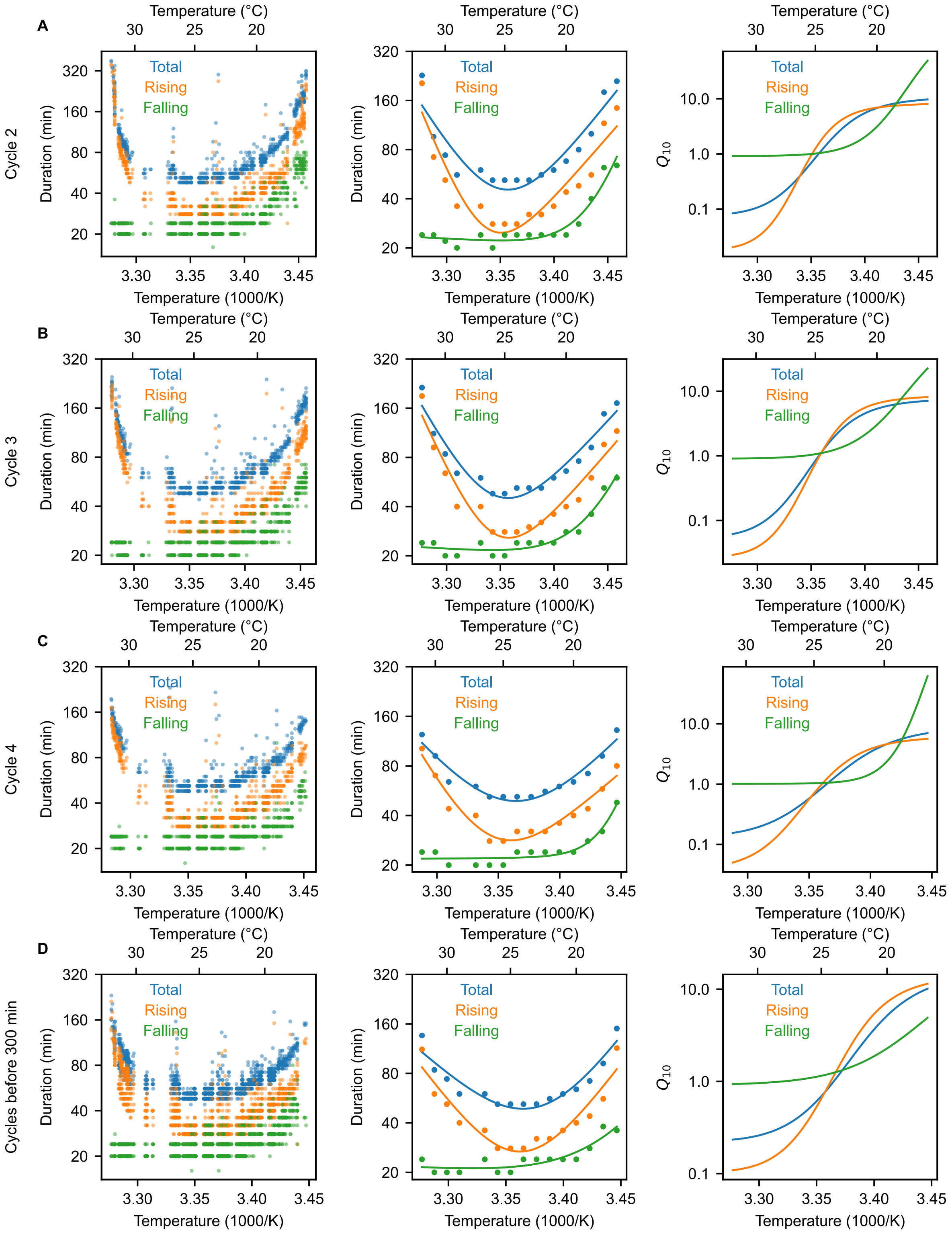


Fig. S8: **Scaling of the duration of the total cell cycle, rising phase, and falling phase for different cycles.** Analogous to Fig. 4C-E: left the raw data, middle the median per rounded temperature with double exponential fit, right the local $Q_{10}$ computed from the double exponential fit. A-C. The data for cycles 2, 3, 4 separately. D. Data from all the cycles that occur in the first 300 minutes of the experiment.

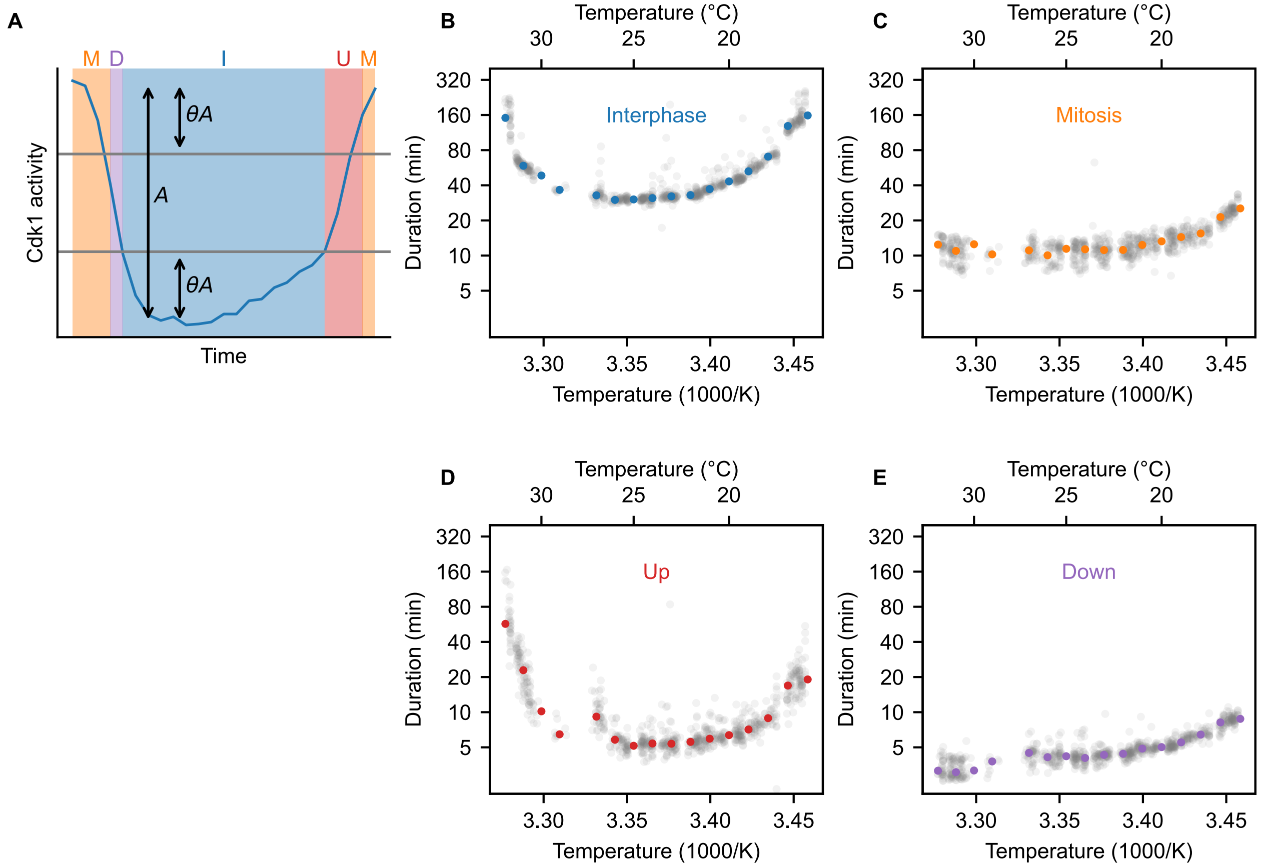


Fig. S9: **Scaling of four parts of the cycle in extracts.** A. Diagram of the method for determining the duration of four parts of the cycle. After determining the minimal and maximal value of the cycle, we determine the amplitude $A$ and two threshold values determined by $\theta$, which we take to be 0.3. Duration of interphase is the time the cycle is below the lower threshold, mitosis is the time the cycle spends above the higher one, and up and down times are the times it spends in between. B-E Duration of these four parts as function of temperature. Light gray dots are datapoints from all droplets, colored dots show the median duration per temperature.

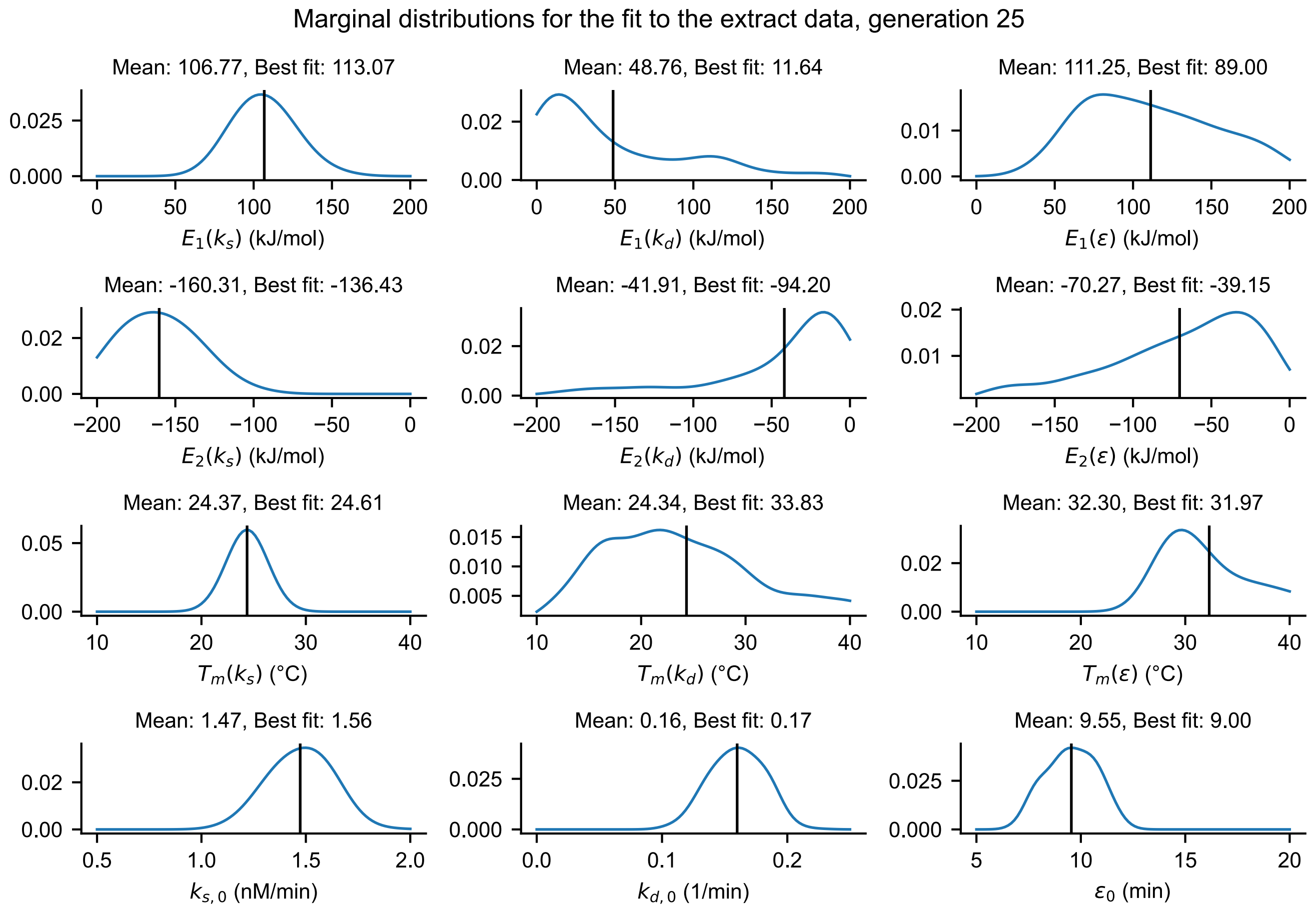


Fig. S10: **Marginal distributions of the different parameters for the optimal fits to extract data.** These parameters determine the scaling of $k_s$, $k_d$ and $\epsilon$ that is shown in Fig. 4. The marginal distribution over 1000 weighted samples, that are the result of the ABC algorithm, is shown. Smooth distribution obtained by Gaussian kernel density. Black line indicates the mean. The 'Best fit' quoted corresponds to the value of the parameter for the sample with least distance to the data.

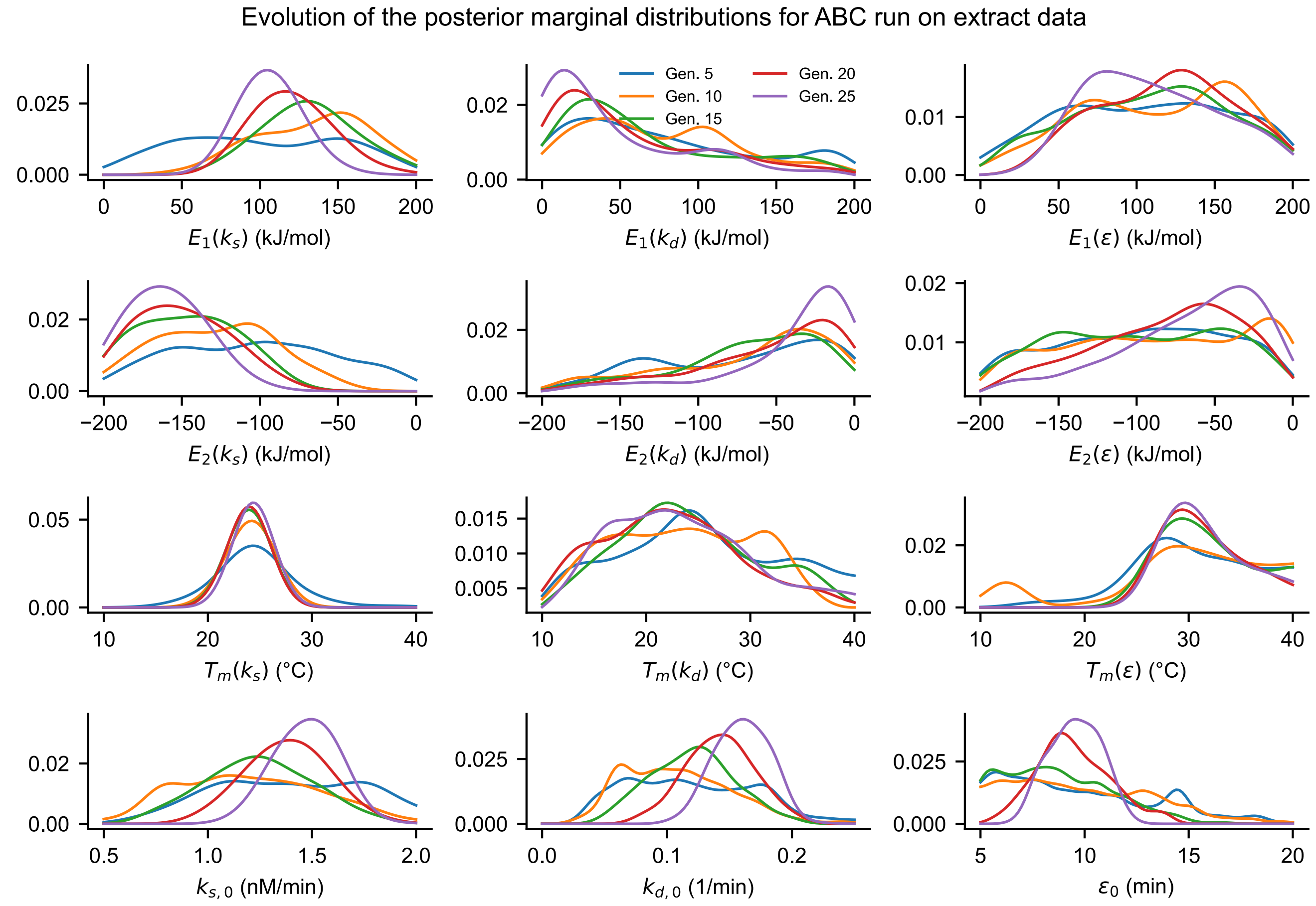


Fig. S11: **Evolution of the marginal distributions of the parameter sets over the course of the ABC algorithm.** Similar to Fig. S10, but the distributions at different generations of the ABC algorithm are shown.

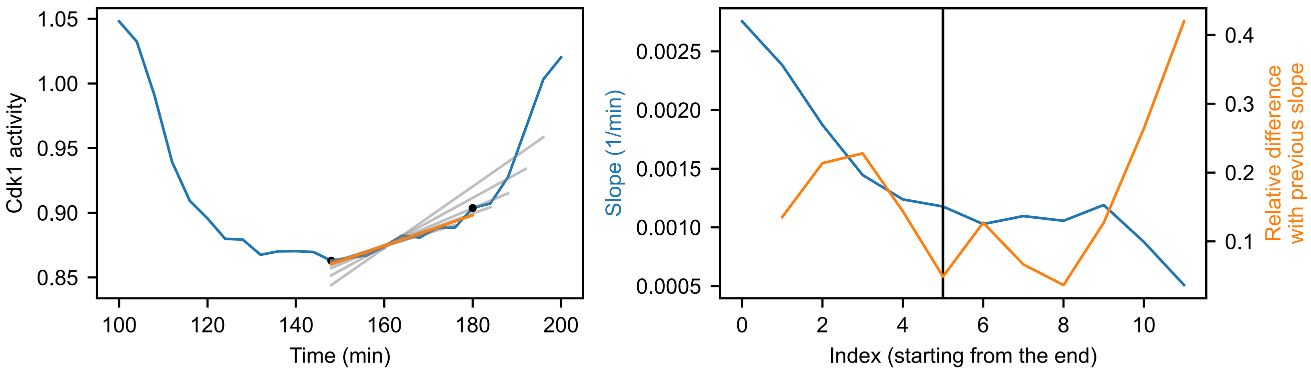


Fig. S12: **Determination of the cyclin synthesis rate $k_s$ from the time series.** Shows what is explained in Supplementary Note 4.

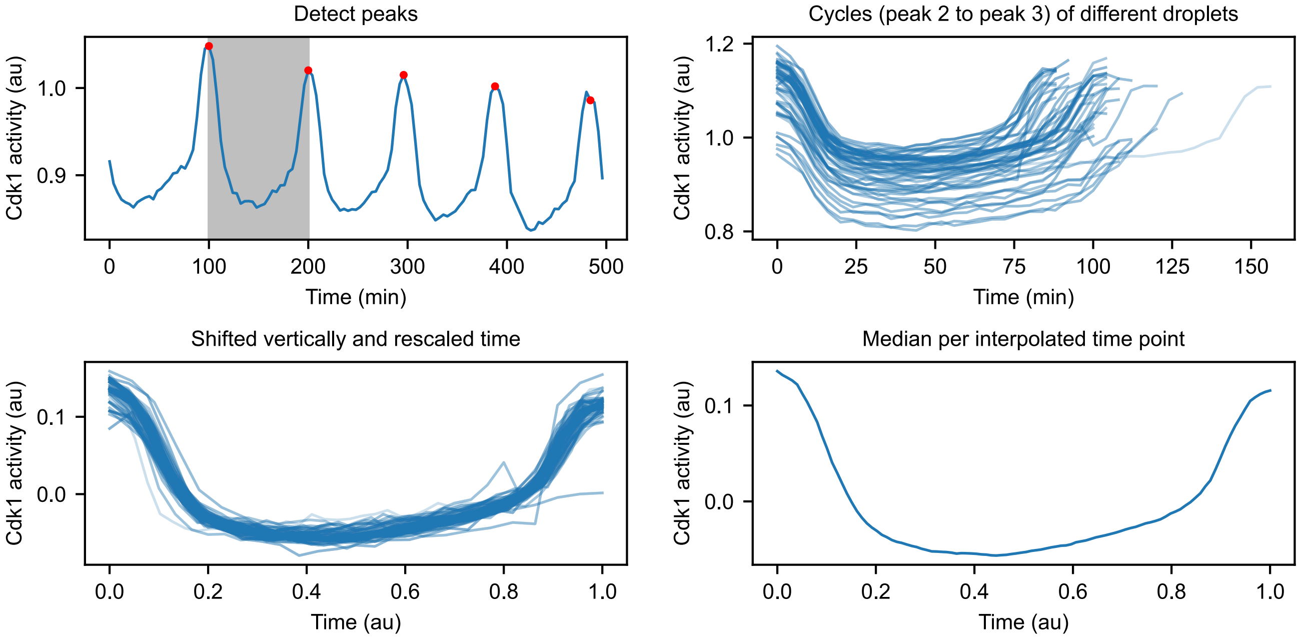


Fig. S13: **Determination of the average cycle shape.** Shows what is explained in Supplementary Note 5. In this example, $T = 22°\text{C}$ is shown. the bottom right panel shows the final average cycle.

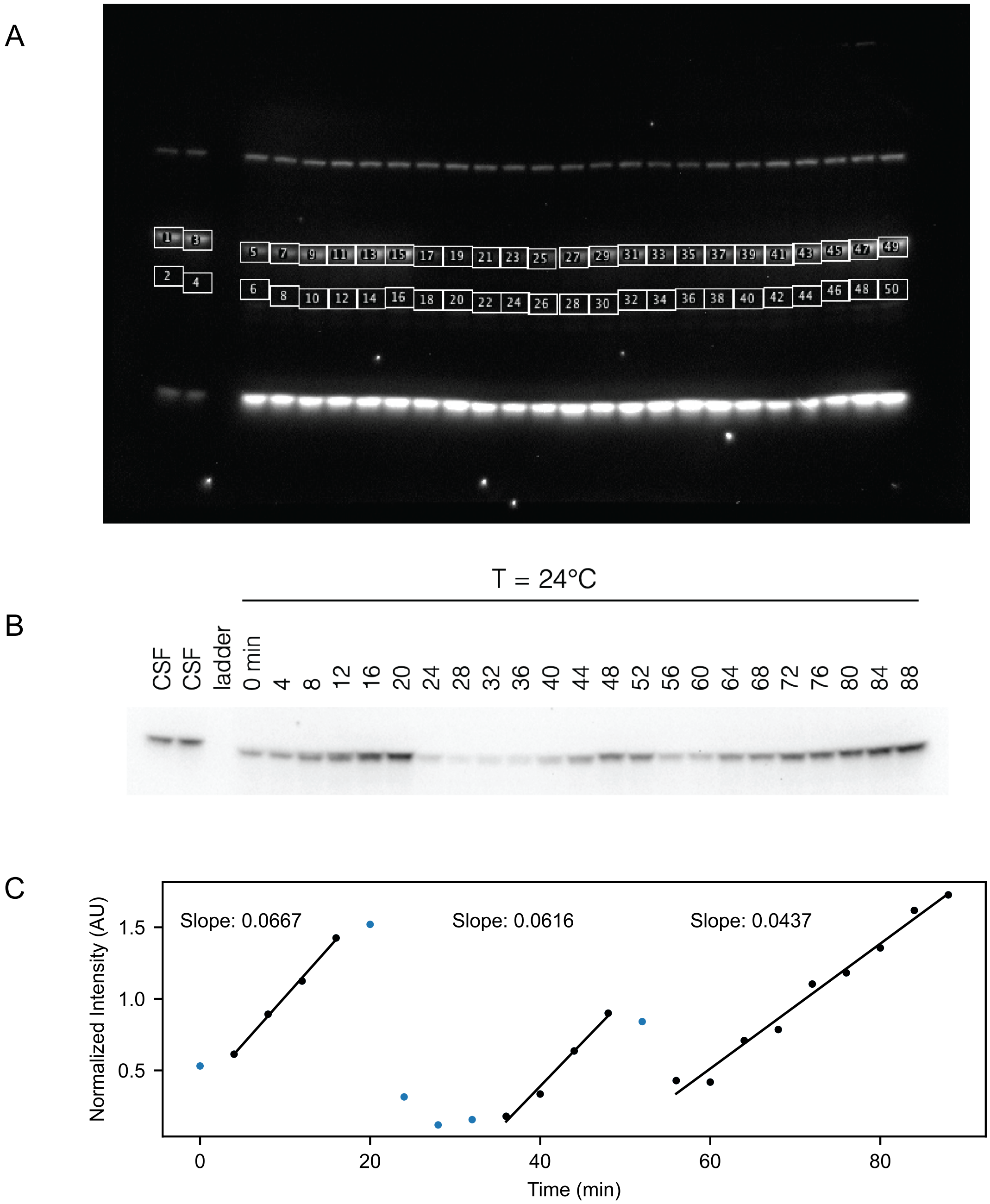


Fig. S14: **Measuring cyclin synthesis rates using quantitative Western blotting**. A. Time course of a representative Western blot using anti-cyclin B2 antibody on a cycling frog egg extract. B. Selected region of same Western blot as in A. C. Quantification of the Western blot by calculating the integrated density for each band and subtracting the background using FIJI, we obtain the intensity for each time point. This value is then divided by the average intensity for a CSF extract, to obtain the normalized intensities. To obtain the cyclin synthesis rates, we next fit the slopes for each of the cycles for the points indicated in black.

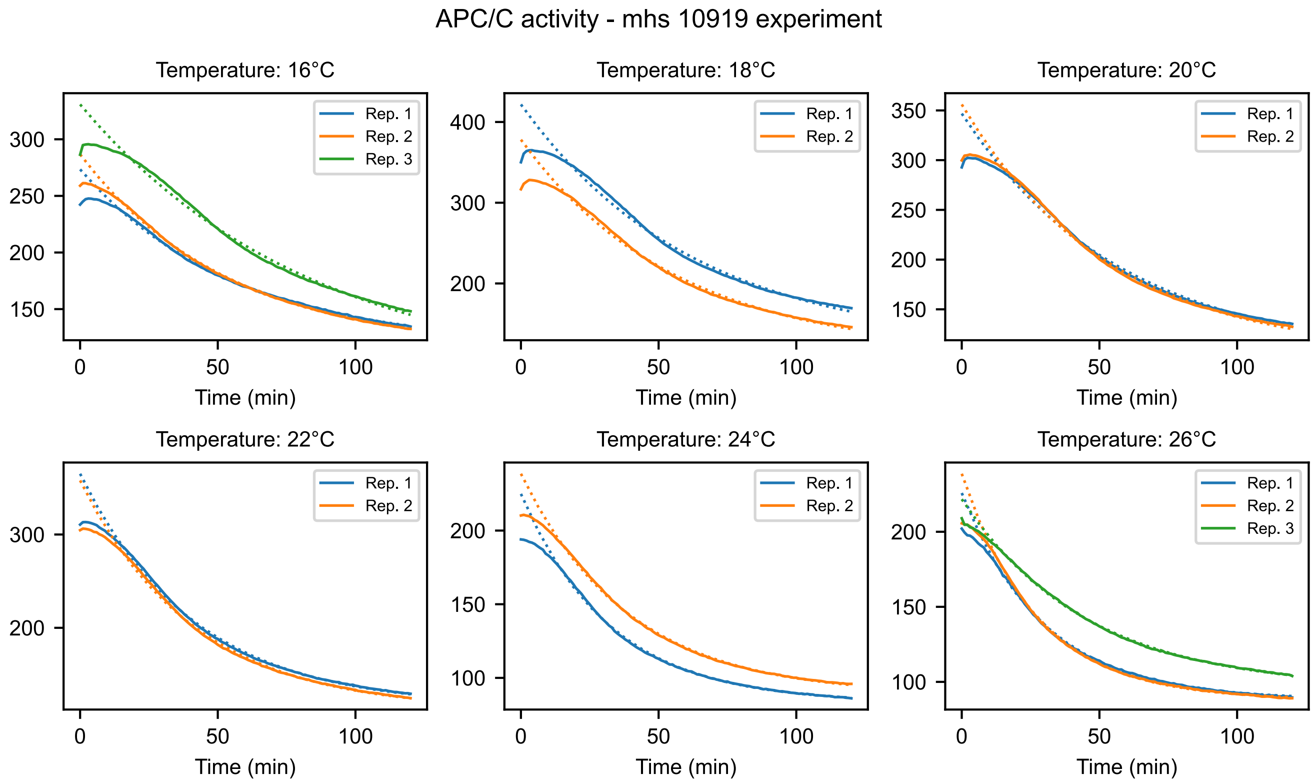


Fig. S15: **Time series for measuring APC activity.** Time series and fits, from which the rates in Fig. 6 are obtained. Dotted lines are fits of the form $y = Ae^{-kt} + B$. Details in Supplementary Note 6.

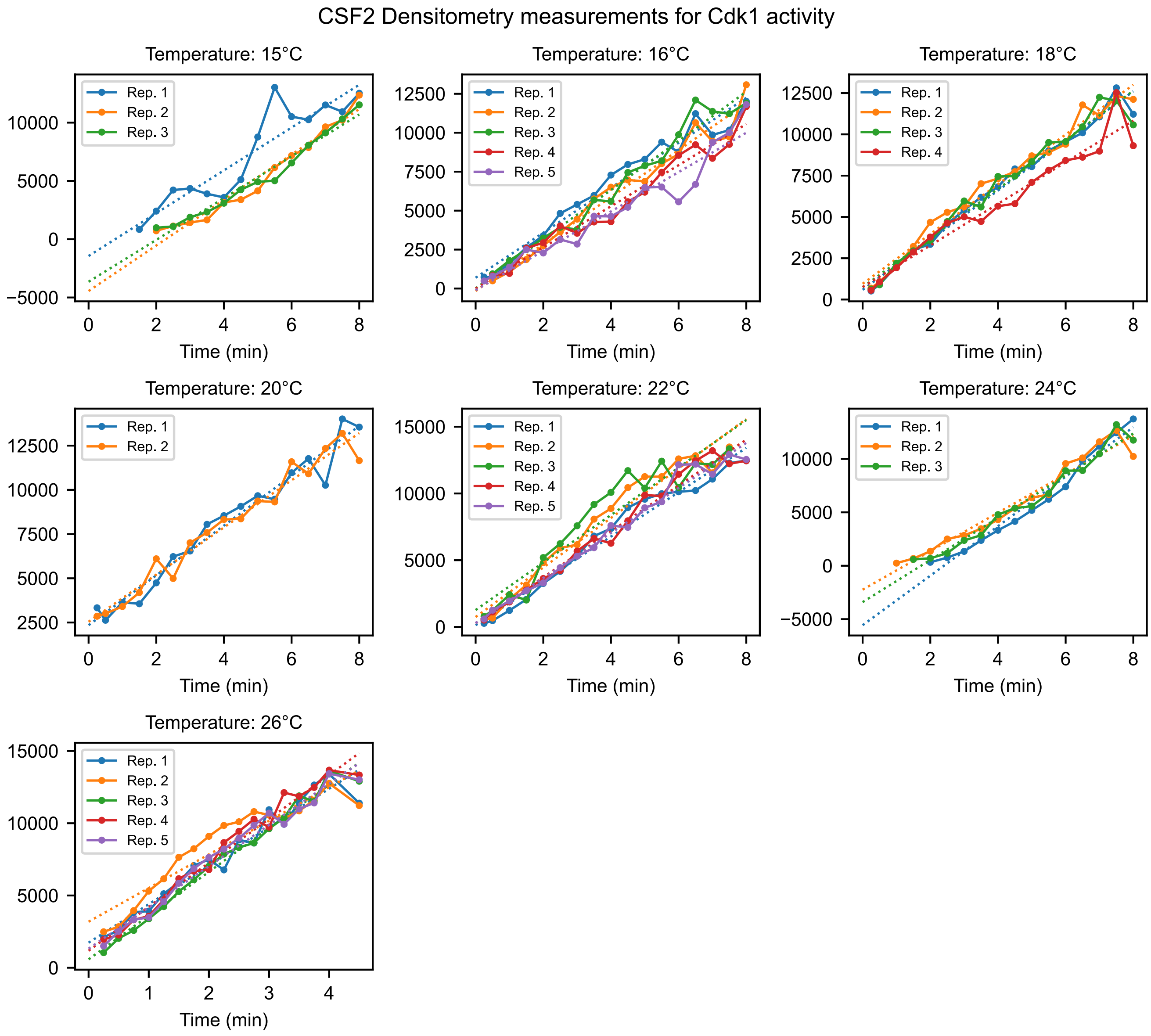


Fig. S16: **Time series for measuring Cdk1 activity.** Time series and fits, from which the rates in Fig. 6 are obtained. Dotted lines are fits of the form $y = at + b$. Details in Supplementary Note 6.

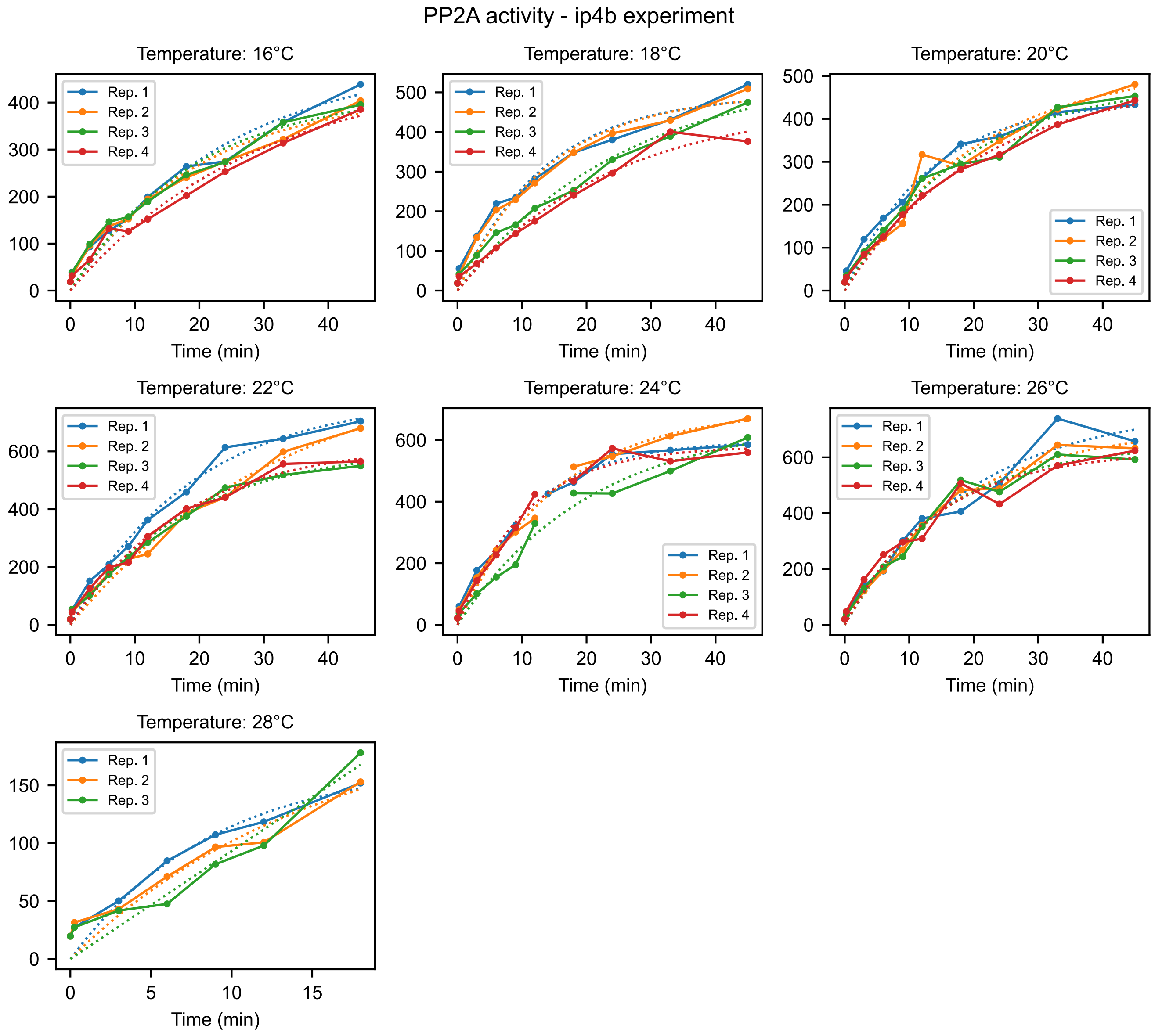


Fig. S17: **Time series for measuring PP2A activity.** Time series and fits, from which the rates in Fig. 6 are obtained. Dotted lines are fits of the form $y = A(1 - e^{-kt})$. Details in Supplementary Note 6.

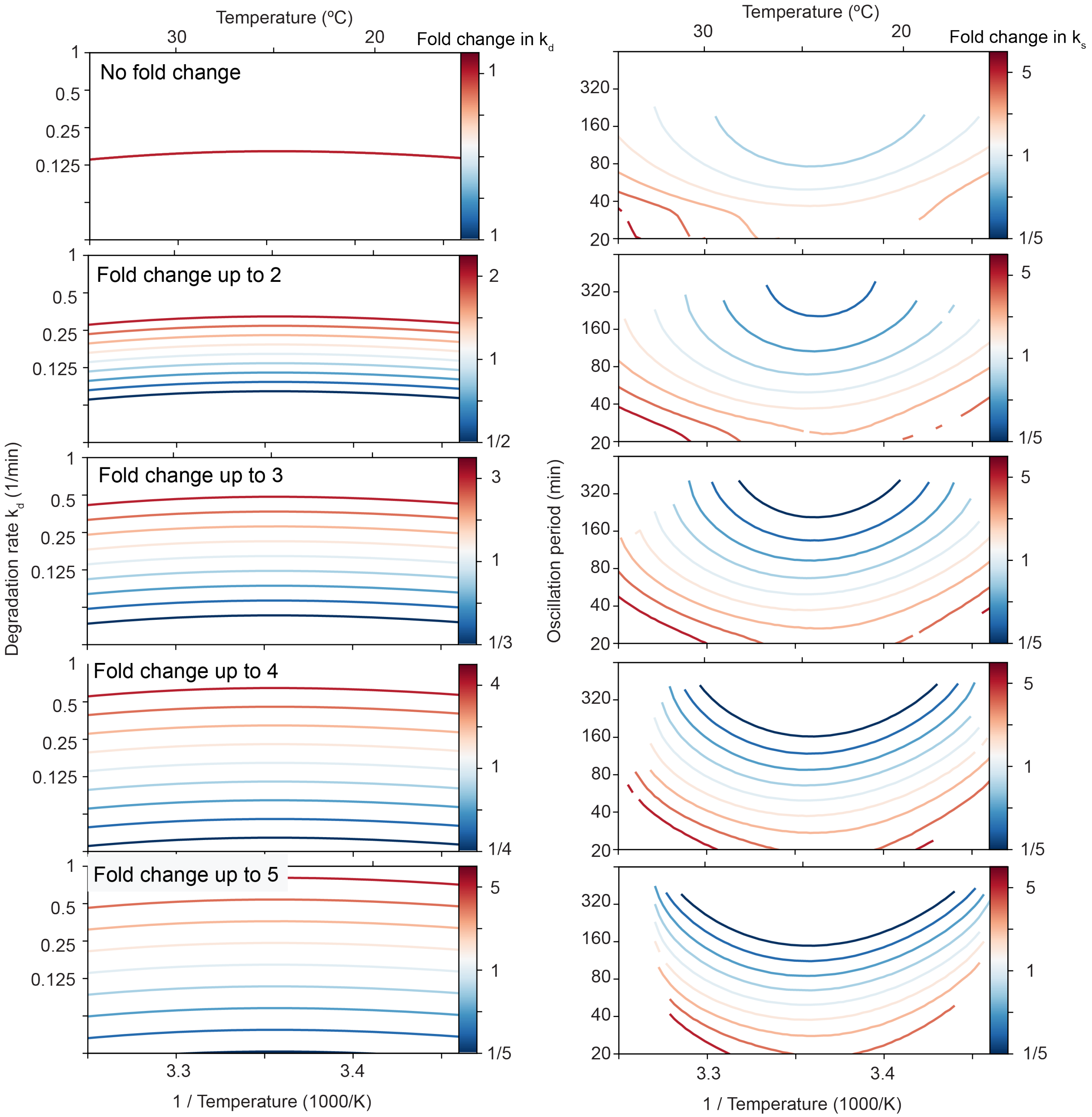


Fig. S18: **Decreasing the cyclin synthesis rate decreases the viable temperature range** Influence of changing the basal cyclin synthesis and the basal cyclin degradation rate by varying factors. In all simulations the basal cyclin synthesis rate is increased and decreased by a factor up to 5, similarly as shown in Fig. 7A. Additionally, from top to bottom we allow for increasing changes of the degradation rate as well. In the top panels, the basal degradation rate is kept constant as the basal cyclin synthesis rate is scaled. The lower panels show increasing fold changesin the basal degradation rate up to a scaling factor of 5, similar as for cyclin synthesis. Larger differences in cyclin synthesis and degradation rates (larger differences in scaling) lead to stronger reductions of the viable temperature range upon decreasing cyclin synthesis rate.
